# The Evolutionary Origins and Ancestral Features of Septins

**DOI:** 10.1101/2024.03.25.586683

**Authors:** Samed Delic, Brent Shuman, Shoken Lee, Shirin Bahmanyar, Michelle Momany, Masayuki Onishi

## Abstract

Septins are a family of membrane-associated cytoskeletal GTPases that play crucial roles in various cellular processes, such as cell division, phagocytosis, and organelle fission. Despite their importance, the evolutionary origins and ancestral function of septins remain unclear. In opisthokonts, septins form five distinct groups of orthologs, with subunits from multiple groups assembling into heteropolymers, thus supporting their diverse molecular functions. Recent studies have revealed that septins are also conserved in algae and protists, indicating an ancient origin from the last eukaryotic common ancestor. However, the phylogenetic relationships among septins across eukaryotes remained unclear. Here, we expanded the list of non-opisthokont septins, including previously unrecognized septins from rhodophyte red algae and glaucophyte algae. Constructing a rooted phylogenetic tree of 254 total septins, we observed a bifurcation between the major non-opisthokont and opisthokont septin clades. Within the non-opisthokont septins, we identified three major subclades: Group 6 representing chlorophyte green algae (6A mostly for species with single septins, 6B for species with multiple septins), Group 7 representing algae in chlorophytes, heterokonts, haptophytes, chrysophytes, and rhodophytes, and Group 8 representing ciliates. Glaucophyte and some ciliate septins formed orphan lineages in-between all other septins and the outgroup. Combining ancestral-sequence reconstruction and AlphaFold predictions, we tracked the structural evolution of septins across eukaryotes. In the GTPase domain, we identified a conserved GAP-like arginine finger within the G-interface of at least one septin in most algal and ciliate species. This residue is required for homodimerization of the single *Chlamydomonas* septin, and its loss coincided with septin duplication events in various lineages. The loss of the arginine finger is often accompanied by the emergence of the α0 helix, a known NC-interface interaction motif, potentially signifying the diversification of septin-septin interaction mechanisms from homo-dimerization to hetero-oligomerization. Lastly, we found amphipathic helices in all septin groups, suggesting that curvature-sensing is an ancestral trait of septin proteins. Coiled-coil domains were also broadly distributed, while transmembrane domains were found in some septins in Group 6A and 7. In summary, this study advances our understanding of septin distribution and phylogenetic groupings, shedding light on their ancestral features, potential function, and early evolution.

## INTRODUCTION

Septins are a family of paralogous cytoskeletal GTPases that associate with one another in defined stoichiometries to create nonpolar filaments. The first four septin genes (*CDC3*, *CDC10*, *CDC11*, and *CDC12*) were identified in a cell-cycle defective screen in *Saccharomyces cerevisiae* (Hartwell, 1971; Hartwell et al., 1974). Detailed molecular characterization of these septins showed that each gene encodes a distinct septin subunit that associates with other septin subunits to create filaments and other higher-order structures such as rings on the plasma membrane (Byers and Goetsch, 1976; Field et al., 1996; Longtine et al., 1996; McMurray and Thorner, 2008). It was later shown that septin assembly and filamentation are influenced by lipid composition of membranes (Bertin et al., 2010).

A septin subunit is comprised of a core GTPase domain and variable N- and C-terminal extensions (NTE and CTE). The GTPase domain is responsible for binding and hydrolyzing GTP, as well as mediating septin-septin interactions and polymerization (Sirajuddin et al., 2007; Hussain et al., 2023). The N-terminal domain of septins often contains a polybasic domain (PB1) directly upstream of the start of the GTPase domain, which plays critical roles in lipid recognition and septin polymerization (Omrane et al., 2019; Cavini et al., 2021). Depending on the septin subunit, the C-terminal domain can contain a coiled-coil domain which has been proposed to mediate lateral pairing of septin filaments (Leonardo et al., 2021). Additionally, some subunits also possess an amphipathic helix (AH) which has been shown to allow septins to recognize micron-scale curvature (Bridges et al., 2016; Cannon et al., 2019). The structure of septin protomers has been described using the human SEPT2/6/7 heterohexameric complex, which unequivocally identified two binding interfaces for septin subunits (Sirajuddin et al., 2007): The G-interface is defined as the face of the subunit with the GTP-binding pocket, where *trans* interactions with an opposing subunit stimulates GTP hydrolysis, whereas the NC-interface is the opposite face of the subunit. Both interfaces can be involved in homomeric and heteromeric dimerization events.

Previous phylogenetic analyses of opisthokont septins identified conserved residues within the G- and NC-interfaces (Pan et al., 2007; Auxier et al., 2019; Shuman and Momany, 2021). Additionally, these analyses provided an evolutionary basis for the modularity of septin paralogs in support of Kinoshita’s rule, which states that septins belonging to the same phylogenetic group can replace one another within the canonical protomer (Kinoshita, 2003b; Pan et al., 2007) For example, human SEPT3, 9, and 12 all belong to Group 1A and can replace one another within a protomer. Thus, these phylogenetic analyses can provide structural and biochemical insights into the assembly of septins.

Most of the cellular, biochemical, and phylogenetic characterizations of septin proteins have been from the opisthokont (animal & fungal) lineage. The presence of septins outside of opisthokonts was initially noted by Versele & Thorner, who mentioned the presence of bona fide septins in *Chlamydomonas reinhardtii* & *Nannochloris spp.* (Versele and Thorner, 2005). Subsequent studies in the green algae *Nannochloris bacillaris* and *Marvania geminata* and the ciliate *Tetrahymena thermophilus* characterized the localization of septins outside of the opisthokont paradigm. In the former, immunofluorescence studies using an antibody against the single septin in *N. bacillaris* showed its localization at the division site of both algae (Yamazaki et al., 2013). In the latter, septins were reported to localize to the mitochondria scission sites and proposed to regulate mitochondrial stability via autophagy pathways (Wloga et al., 2008). Additional septins have since been identified in some other algae and protists (Nishihama et al., 2011; Yamazaki et al., 2013; Onishi and Pringle, 2016); however, the phylogenetic relationship of these non-opisthokont septins remained unclear.

In this work, we provide an update to the distribution of septins across the eukaryotic tree of life and a rigorous phylogenetic analysis to compare their relationship to previously identified septin groups. We trace the evolution of structural motifs within the septin GTPase domains by combining ancestral sequence reconstruction and machine-learning 3D structural prediction. Lastly, we trace the gains and losses of septin-associated features in the NTE and CTE, such as the polybasic domain, coiled-coil, AH, and putative transmembrane domains to assess their evolutionary origins.

## MATERIALS AND METHODS

### Identification of New Septin Sequence**s**

To identify new non-opisthokont septin sequences, we utilized both the Joint Genome Institute Phycocosm webpage (https://phycocosm.jgi.doe.gov/) and the NCBI Genome database (https://blast.ncbi.nlm.nih.gov/). We used the initial set of queries consisting of *Chlamydomonas*, *Symbiodinium*, and *Paramecium* septins. These searches identified several septins in the phyla in which they have not been reported. To enhance the chance of finding new sequences in these and other divergent branches, we added *Porphyra*, *Ectocarpus*, and *Cyanophora* to the list of queries and performed additional searches (Table 1; Supplementary File 1). BLASTP searches were performed on November 14, 2021 using a BLOSUM62 matrix, E-value cutoff of 1x10^-5^, word size of 3, and filtered low complexity regions. The JGI database searches used proteomes from Excavata, Archeaplastida, Rhizaria, Heterokonta, and Alveolata (Supplementary File 2). Due to the limited availability of information for ciliate species on JGI, additional searches were performed using the NCBI database, specifically focusing on Alveolata (taxid:33630) (Supplementary File 2). Identified sequences were further examined manually for the presence of G-motifs (G1, G3, and G4) and S-motifs (S1-S4) to confirm that they are bona fide septins. Opisthokont septins were selected from (Auxier et al., 2019).

**Table 1:** Query sequences used in BLASTP searches Phylum Species Identifier.

| Phylum | Species | Identifier |
| --- | --- | --- |
| Chlorophyta<br>(green algae) | *Chlamydomonas reinhardtii* | Cre12.g556250 |
| Glaucophyta | *Cyanophora*<br>*paradoxa* | 13652g13185t1 |
| Rhodophyta<br>(red algae) | *Porphyra*<br>*umbilicalis* | 6951 |
| Phaeophyceae<br>(brown algae) | *Ectocarpus*<br>*siliculosus* | CBN74010 |
| Ciliophora<br>(ciliates) | *Paramecium*<br>*tetraurelia* | CAI38984 |
| Dinoflagellates | *Symbiodinium*<br>*minutum* | symbB1.v1.2.007989.t1<sup>a</sup> |

<sup>a</sup> This transcript encodes a very long 4484-aa predicted protein. See Onishi & Pringle (2016) for details. The 560-aa amino-terminal sequence containing the septin GTPase domain was used as query.

### Phylogenetic Analysis and Ancestral Sequence Reconstruction

Phylogenetic trees were constructed following the methodology described by (Auxier et al., 2019). A total of 131 opisthokont and 123 non-opisthokont septins were used; as an outgroup, several prokaryotic YihA proteins were also included (Supplementary File 3). Sequences were first aligned using the constraint-based alignment tool (COBALT) (Papadopoulos and Agarwala, 2007), which incorporates information about protein domains in a progressive multiple alignment. This tool biases the alignment within the septin GTPase domain. To remove regions of randomly similar sequences from the alignment, we employed ALISCORE and ALICUT (Misof and Misof, 2009; Kück et al., 2010; Kueck, 2017). ALISCORE identifies regions of ambiguous alignment, which were subsequently removed using ALICUT. This process resulted in a reduced MSA file containing highly conserved regions within the GTPase domain (Supplementary File 4), which was then used to generate the phylogenetic tree.

Tree generation was performed using the CIPRES gateway (Miller et al., 2010), employing RAxML-HPC v.8 on XSEDE with the PROTCAT substitution model and the LG protein matrix and a rapid 1000 bootstrap analysis. The generated trees were visualized using the Rstudio package “ggtree.” Bootstrap values displayed on the trees have been limited to values greater than 25.

For ancestral sequence reconstruction (ASR), we utilized the FASTML server for maximum-likelihood computing of the ancestral states (Ashkenazy et al., 2012). Due to limitations with the FASTML server, we reduced our list of septin sequences from 254 to 200 by removing some sequences from some fungal species and all sequences from the genus *Paramecium* except for the species *tetraurelia*. The resulting 200 sequences (Supplementary File 5) were aligned using COBALT alignment. As ASR provides meaningful interpretation when the entire protein sequence is provided, we did not utilize ALISCORE and ALICUT processing. To generate a new phylogenetic tree, we used the IQTree webserver (http://iqtree.cibiv.univie.ac.at/) with an automatic amino acid replacement matrix, 1000 ultrafast bootstraps, and all other default parameters (Trifinopoulos et al., 2016; Minh et al., 2020). This tree reproduced the same phylogenetic groupings and general branching patterns as our more rigorous ALISCORE and ALICUT processed tree. Nodes of interest, including parental nodes for the septin phylogenetic groups, opisthokont and protist divide, and the last eukaryotic common ancestor (LECA) node, were defined based on the joint reconstruction output file and labeled in Supplementary File 5. The protein sequences at these nodes were extracted and referred to as the ancestral septins.

### AlphaFold Predictions and Search for Polybasic Domains in N-terminal Extension

AlphaFold predictions were executed using the Colabfold Google notebook v1.3.0. The specific parameters can be found within the “config.json” file in each respective folder. Due to computational limitations of AlphaFold with extremely long sequences, some sequences required trimming. The objective of trimming was to preserve the entire GTPase domain and the CTE while reducing the sequence length to a manageable size (approximately 800 amino acids). Generally, the protein sequence was truncated from the N-terminal end. Predictions primarily used an MMseqs2 MSA. Five models with three recycles each were generated and the highest-ranking model was selected (Supplementary File 6). The resulting 3D structures were visualized using ChimeraX. Topology diagrams were drawn in Adobe Illustrator, following the convention used in (Cavini et al., 2021). For AlphaFold predictions of *K. flaccidum* and *I. multifiliis* septins, we used version 1.5.2 of the ColabFold notebook. The structures were visualized using ChimeraX and colored according to AlphaFold confidence.

To search for potential polybasic domains in the NTE of our reconstructed ancestral sequences, we developed a Python script that uses a sliding 10-amino-acid window to calculate the local average isoelectric point and plots this value against the first amino acid position across the entire protein length. To focus solely on the NTE, which is where PB1 in extant septins is primarily located, we aligned the ancestral septins to the GTPase domain of *S. cerevisiae* Cdc3 using CLUSTALω. Only residues before the start of the GTPase domain were plotted. To visualize the multiple sequence alignment (MSA) of the ancestral septins, a CLUSTALω alignment was performed without the Cdc3 GTPase domain to compare the amino acid composition between GTPase domain-adjacent polybasic domains. The MSA was visualized using the R package “ggmsa,” and the amino acids were colored according to their properties.

### Identification of amphipathic helices in extant septin sequences

For high-throughput prediction of amphipathic helices, we developed a Python script that consists of two steps of analysis: (1) secondary structure prediction by s4pred (Moffat and Jones, 2021) followed by (2) amphipathicity assessment of α-helices. In (1), secondary structure prediction was performed for the amino acid sequence of a given septin protein using the run_model.py script provided in https://github.com/psipred/s4pred. In (2), either a “fully-helical” or “partially-helical” segment of an amino-acid sequence was extracted by a sliding 18 amino-acid window. In a “partially-helical” segment, at least 6 amino acids at both ends of the 18 amino acid window must be fully helical. For example, while a segment with a prediction “HHHHHHCCCCCCHHHHHH” (6x H – 6x C – 6x H) was permitted, those with “HHHHHCCCCCCHHHHHHH” (5x H – 6x C – 7x H) were not. We included “partially-helical” segments for further assessment because some membrane-bound Ahs could be predicted as “partially helical,” where two helices are broken apart by non-helical sequence (e.g., Sun2 AH: Lee et al., 2023). For each helical segment, the amphipathicity was calculated and assessed similarly to HeliQuest software (Gautier et al., 2008), but with modifications. First, the mean hydrophobic moment value *<µH>* was calculated as previously described (Eisenberg et al., 1982) using the hydrophobicity scale values (Fauchere and Pliska, 1983) based on an assumption that all helices rotate with a 100 degree step. Then, the discriminant factor *D* = 0.944 x *<µH> +* 0.33 x *z* (where z is the net charge) was calculated accordingly to HeliQuest. Finally, the helical segment was considered amphipathic if all of the criteria below were satisfied: i) *D* > 0.68 OR (*<µH>* > 0.4 AND *z* = 0); ii) The hydrophobic face contains at least 3 consecutive bulky hydrophobic residues (L, V, F, I, W, M, Y) (e.g. a hydrophobic face “SYA<u>LLV</u>T” is satisfactory); iii) “Core” of the hydrophobic face does NOT contain any charged residue (”core”: the area of 90° centered around the pole). This search resulted in the identification of 4809 possible AH domains, with the vast majority showing overlap with one another (Supplementary File 7).

We then filtered the data to exclude AHs that are positioned inside of an septin GTPase domain. The GTPase domain of Cdc3 from *S. cerevisiae* was used as a reference to define the start and end residues for the GTPase domain of the other 254 extant sequences. The list of possible AHs of 18 amino acids in length was then screened by excluding those that overlapped with the GTPase domain. Sequences satisfying these criteria were considered to possess an AH (Supplementary File 8) and were highlighted in a cladogram generated using the R package “ggtree.” To generate helical wheel diagrams, individual AH sequences from the dataset were used as input to run the HeliQuest program (Gautier et al., 2008).

### Search for coiled-coil and putative transmembrane domains in extant septin sequences

To identify septins with coiled-coil domain and/or putative transmembrane domains in the set of 254 extant septins, we used the existing annotations on the UniProt database (The UniProt Consortium, 2023) release 2023_04. A BLASTP search using our list of 254 septins as query against the UniprotKB database retrieved 206 hits, for which “Coiled coil” and “Transmembrane” annotations were downloaded from the database. According to the UniProt documentation, these annotations are based on the COILS program (Lupas et al., 1991) with a minimum size of 28 amino acids for coiled-coil domains, and TMHMM and Phobius predictions (Krogh et al., 2001; Käll et al., 2004) for transmembrane domains. For the remaining 48 sequences, manual searches for coiled-coil and transmembrane domains were performed using Cocopred (Feng et al., 2022) and Phobius. These predictions are conservative and unlikely to identify all possible coiled-coil and transmembrane domains; for example, the present analysis identified fewer coiled-coil-containing septins than Auxier et al. (2019), which used the hidden-Markov-model-based Marcoil program. Results of these searches are summarized in Supplementary File 8.

## RESULTS

### Identification of new septin sequences

To search for septin sequences outside of opisthokonts, we compiled a small query list of previously identified septin sequences from algal and protist species (Table 1). These sequences were selected based on their evolutionary diversity, aiming to enhance the chance of identifying septins from various taxa. We conducted BLASTP searches using the BLOSUM62 matrix and an E-value cutoff of 1x10^-5^, utilizing the protein databases available on the Joint Genome Institute’s (JGI) Phycocosm webpage and the Alveolata database on the NCBI BLAST website (see Materials and Methods). These searches revealed previously unreported sequences in multiple taxa under the supergroups Archaeplastida and Chromista, including two species of rhodophyte red algae (*Porphyra umbilicalis* and *Pyropia yezeonsis*) and one species of glaucophyte algae (*Cyanophora paradoxa*) (Figure 1). Our searches also reproduced a previous failure to identify any septin sequences in the entire supergroups of Amoebozoa and Excavata (Fig. 1; Onishi & Pringle, 2016). At lower phylogenetic levels, septins were also not detected in Viridiplantae (land plants) (Fig. 1).

**Figure 1.**
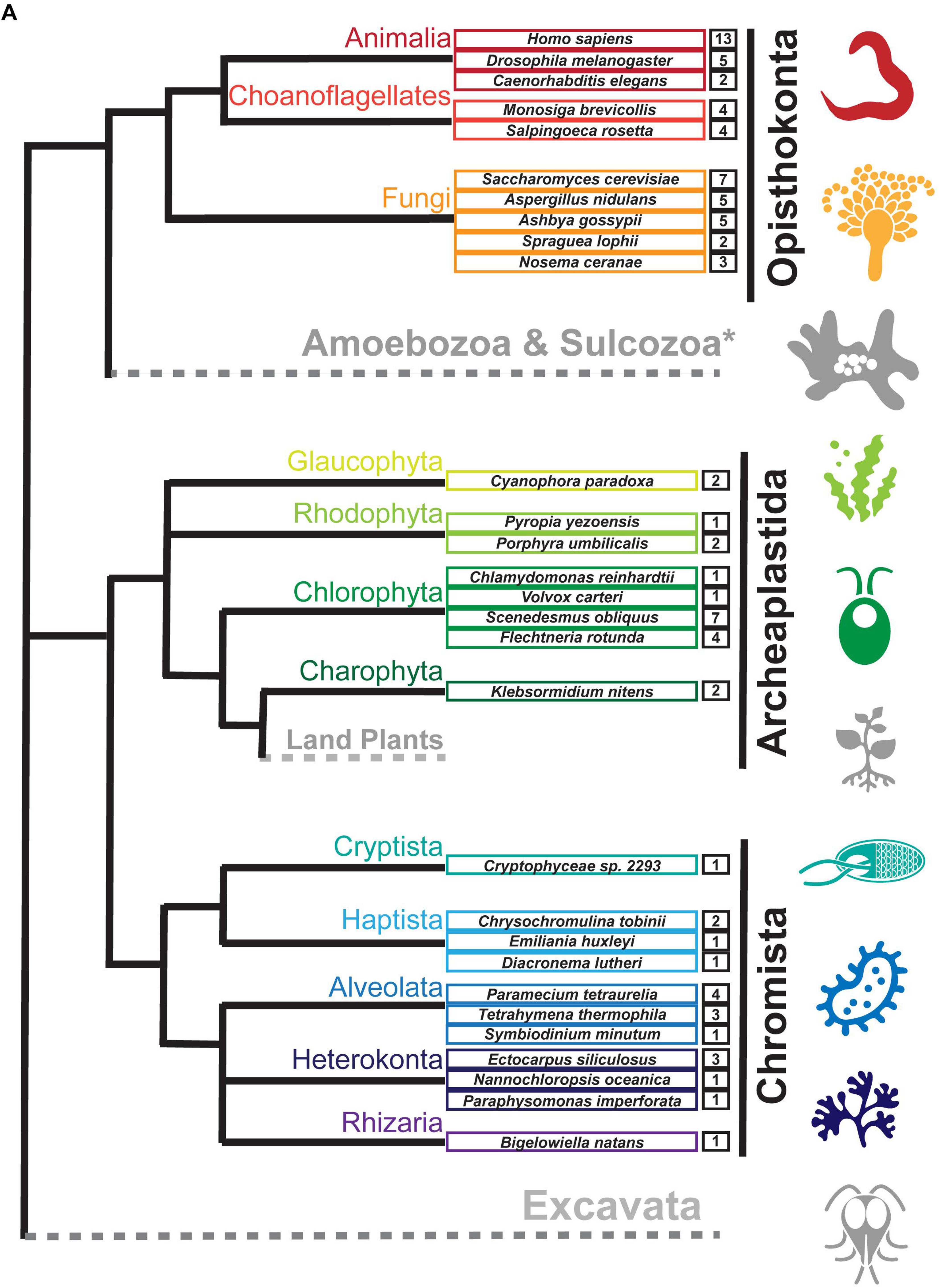
Distribution of septins in non-opisthokont phyla. (**A**) Unrooted taxonomic tree of eukaryotes (based on (Cavalier-Smith, 2018). Gray and dotted branches indicate lineages in which no septin sequence was identified, while black and colored branches represent lineages with identified septins. Representative species are shown and color-matched to their respective lineages, and the total numbers of septin paralogs identified in their genomes are indicated. *Possible septins were identified in *Planoprotostelium fungivorum*; because this is the only example of species with septins within Amoboezoa and Sulcozoa, we could not determine whether they are a result of unique gene retention, horizontal gene transfer, or contamination.

### New septin phylogenetic groups

The discovery of new septin sequences in distant branches of eukaryotes raised questions about their phylogenetic relationship with other septins. Previous studies have classified septins into five groups, but these groupings were defined predominantly based on septin sequences within the opisthokont lineage. We thus combined these new non-opisthokont septin sequences with a preexisting list of opisthokont septins (Auxier et al., 2019) and used the resulting 254 sequences to generate a consensus RAxML tree (Fig. S1) and a simplified cladogram (Fig. 2A). Briefly, the 254 sequences and four prokaryotic YihA NTPases (used here as an outgroup; (Weirich et al., 2008)) were aligned using NCBI’s COBALT alignment tool and processed using ALISCORE and ALICUT to remove ambiguous regions of alignment.

**Figure 2.**
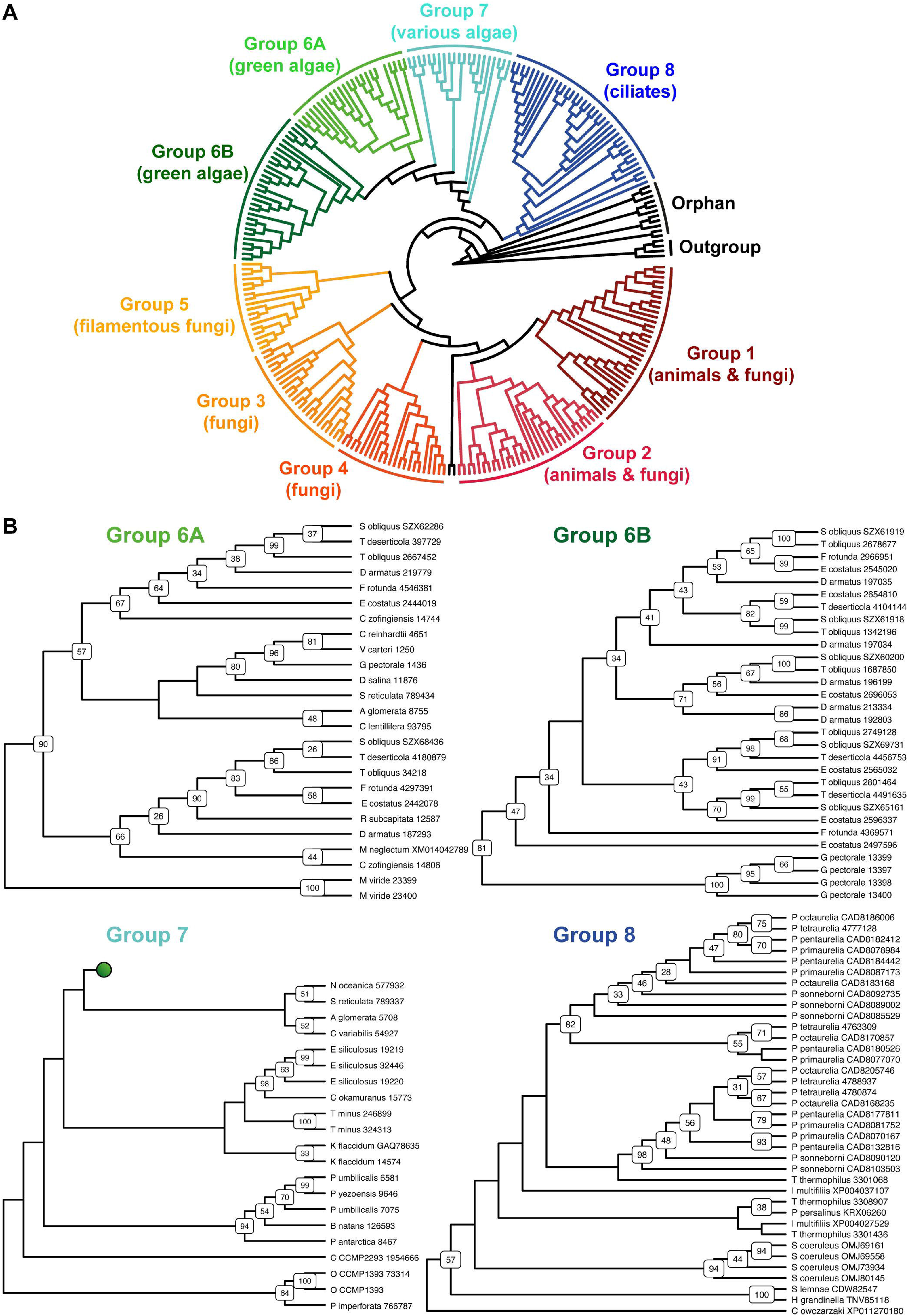
Identification of new septin groups in non-opisthokonts. (**A**) A simplified cladogram representation of a RAxML tree (Fig. S1) of 254 extant septin sequences across eukaryotic lineages. Individual septin phylogenetic clades are color-coded and labeled. The tree is rooted using four prokaryotic YihA proteins as an outgroup. (**B**) Magnified views of the four new phylogenetic clades. See Fig. S2 for the original RAxML trees. Bootstrap values greater than 25 are displayed at nodes.

Consistent with results from previous reports (Momany et al., 2001; Kinoshita, 2003a; Pan et al., 2007; Shuman and Momany, 2021), our phylogenetic analysis grouped the opisthokont septins into five distinct clades (Fig. 2A; Fig. S1): Groups 1 and 2 include septins from both animals and fungi, while Groups 3, 4, and 5 represent fungi-specific clades. Although limited sampling of non-opisthokont septins has previously placed some of them in Group 5 (Onishi and Pringle, 2016; Shuman and Momany, 2021), it is now clear that Group 5 septins are distinct from non-opisthokont septins, consistent with the proposal by Yamazaki et al. (2013).

The non-opisthokont septins themselves form three new groups (Groups 6-8) (Fig. 2B; Fig. S2). Group 6 is a monophyletic group of green algal species divided into two subgroups: Group 6A includes some septins that are encoded as a single gene in the genome, in species such as *C. reinhardtii* and *N. bacillaris* (Versele and Thorner, 2005; Yamazaki et al., 2013). Group 6B, in contrast, exclusively represents septins that appear to have emerged through gene duplication. For example, of five septins in the green alga *Gonium pectorale,* only one belongs to Group 6A while the remaining four belong to Group 6B (Fig. 2B; Fig. S2). The genes for these four septins form a cluster in the assembled *G. pectorale* genome. (Scaffold_65:140824 - 165695), suggesting a very recent gene duplication event. Similarly, of the seven septins in *Desmodesmus armatus,* five belong to Group 6B (Fig. 2B; Fig. S2). Group 7 is a paraphyletic group composed of septins from various groups of algae, such as additional green algae (e.g., *Symbiochloris reticulata*), heterokonts (*Ectocarpus siliculosus*), haptophytes (*Chrysochromulina tobinii*), cryptophytes (*Crytophyceae sp. CCMP2293*), chlorarachniophytes (*Bigelowiella natans*), and rhodophytes (*P. umbilicalis*) (Fig. 2B; Fig. S2). Finally, Group 8 is a monophyletic group comprised exclusively of septins from ciliates, except for one highly divergent sequence from the unicellular opisthokont *Capsaspora owczarzaki*. Within Group 8, septins from *Paramecium* and *Stentor coeruleus* formed genus-specific clades, suggesting recent expansion events of septin genes within their lineages (Fig. 2B; Fig. S2).

Several non-opisthokont sequences are currently not classified in Groups 6-8 because their phylogenetic positioning was sensitive to the programs and parameters used (Fig. 2A; Fig. S1). These include sequences from glaucophytes (*C. paradoxa*), dinoflagellates (*S. minutum, Pseudonitzschia multistrata*), and coccolithophores and related haptophytes (*Emiliania huxleyi, Phaeocystis globosa, C. tobinii, Diacronema lutheri*). Curiously, a septin from *Fonticula alba*, an opisthokont cellular slime mold, also belonged to this orphan group. Additional sampling of sequences from these and related species will likely help improve the confidence in their phylogenetic positioning.

### Conservation of G-interface residues in non-opisthokont septins

In previous studies, septins from Groups 1-5 were found to have several highly conserved regions in their GTPase domains (Fig. 3A) that participate in inter-subunit contacts across the G- and NC-interfaces (Fig. 3BC; Pan et al., 2007; Auxier et al., 2019; Shuman and Momany, 2021; Castro et al., 2023). To gain insights into the evolution of these interfaces in septins across the eukaryotic tree, we expanded the alignment to all 254 septins and generated a Weblogo representation for each septin group (Fig. 3D). In general, the GTPase-specific motifs (G1, G3, G4), septin-specific motifs (S2, S3, S4) except for the S1 motif (Pan et al., 2007; Auxier et al., 2019; Nishihama et al., 2011; Onishi & Pringle, 2016), and some key residues in the septin-unique element are all well conserved. More specifically, most of the key residues in the five G-interfaces (Gig1-Gig5) are all conserved, except for Gig2 which appears to be variable in Group 8 (Fig. 3D). In contrast, key residues in the four NC-interfaces (NCig1-4) are poorly conserved in Groups 6b, 7, and 8. These results suggest that non-opisthokont septins may primarily form homo- or hetero-dimers through the G-interface, and further addition of subunits through NC-interfaces may be limited to Group 6a. In support of this speculation, we found a unique arginine residue that is highly conserved in many Group 6-8 septins but not in Groups 1-5 (Fig. 3D); similar “arginine (R-) fingers” are found in other GTPases that form G-dimers (Koenig et al., 2008; Schwefel et al., 2013; see below).

**Figure 3.**
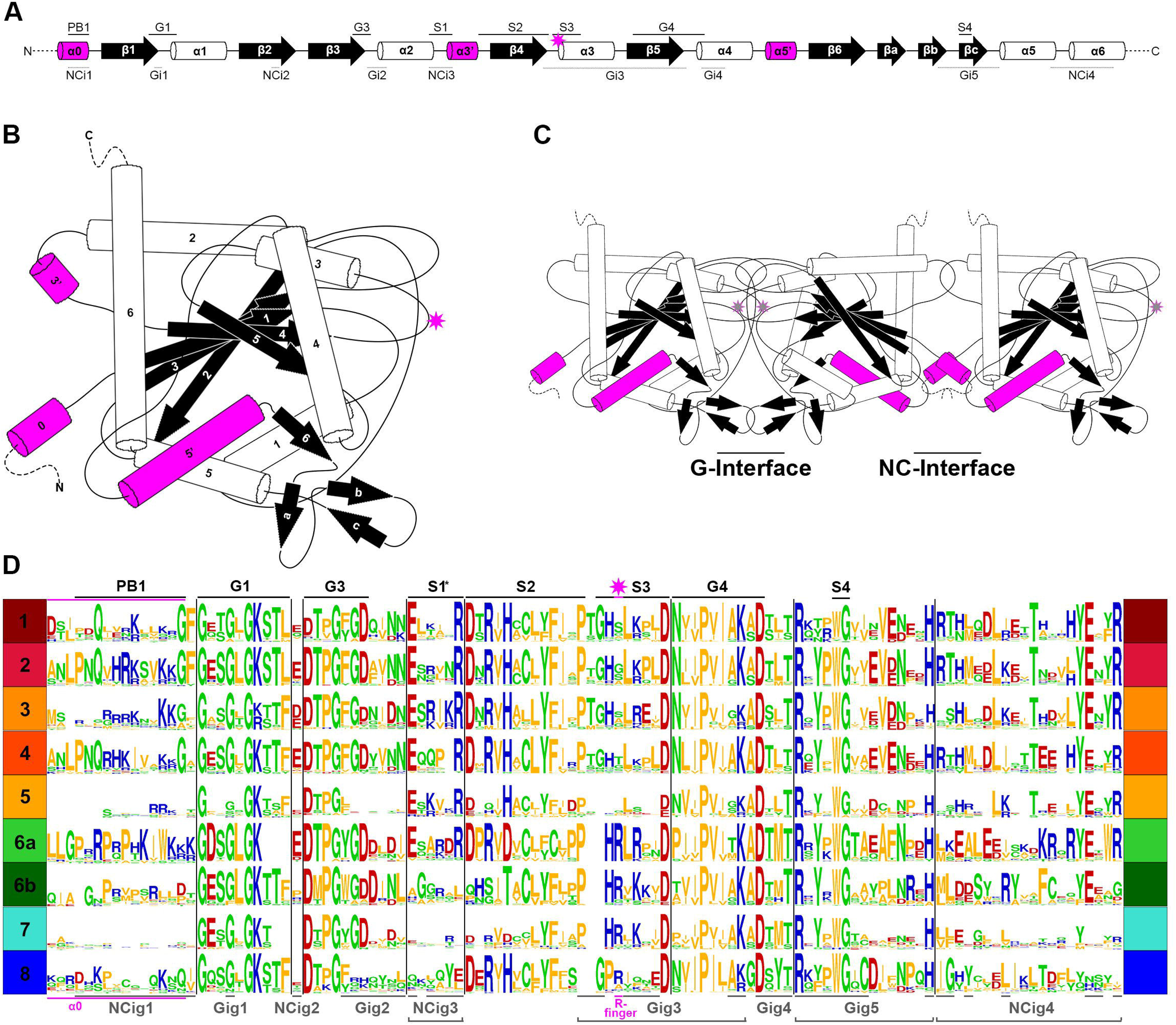
Patterns of conservation and diversity of interface motifs across septin Groups. (**A**) Topology diagram of the GTPase domain secondary structures from N to C-terminus. Conserved GTPase motifs and septin motifs are noted above by black lines (based on Grupp and Gronemeyer 2023) and the NC and G-interacting group regions are noted below by dashed lines (based on Auxier et al 2019). The typical position of the R-finger (when present) is indicated by the pink star. (**B**) A folded septin monomer. This aggregate depiction includes all predicted domains across eukaryote septins. Relative positions of secondary structures are based on PDB structures 7M6J and 8FWP (Mendonça et al., 2021; Grupp and Gronemeyer, 2023; Marques da Silva et al., 2023). (**C**) A septin trimer approximating interactions through their G- and NC-interfaces, based on PDB structure 7M6J. Grey stars with pink outline indicate the predicted positions of R-fingers if they are present in the subunits forming an interface. (**D**) Weblogo representation of select septin motifs, interacting groups, and structural elements across the eukaryotic septin groups. GTPase motifs and septin motifs are depicted above in black, and NC and G-interacting group regions are depicted below in grey. *Note, the location of S1 in groups 6a-8 was determined by relative position in the alignment to the beginning of S2. This loop region which resides between α2 and β4 has considerable sequence length variability and also includes a region where the α3’ helix is predicted.

### Reconstituted ancestral septins suggest that the arginine finger in the G-interface is an ancestral feature

To delve deeper into the evolution of the structural motifs within the septin GTPase domain, we used ancestral sequence reconstruction (ASR) (Ashkenazy et al., 2012) to resurrect ancestral septins. Due to the limitations of the program used, we reconstructed an IQTree of 200 of the 254 septins (Fig. 4A; Fig. S3). The grouping of septin clades and the overall topology of the tree were largely consistent with the RAxML tree (Fig. 2). Using this IQTree, ASR prediction was made for several key nodes representing Groups 1-8 and their parental nodes, and then AlphaFold2 (Jumper et al., 2021) was used to predict their 3D structures for the GTPase domain and the C-terminal extension (see Materials and Methods). Perhaps unsurprisingly given the conservation of the extant sequences (Fig. 3D), the tertiary structures of the ancestral sequences all appeared similar among themselves and with experimentally determined septin structures (Fig. S4).

**Figure 4.**
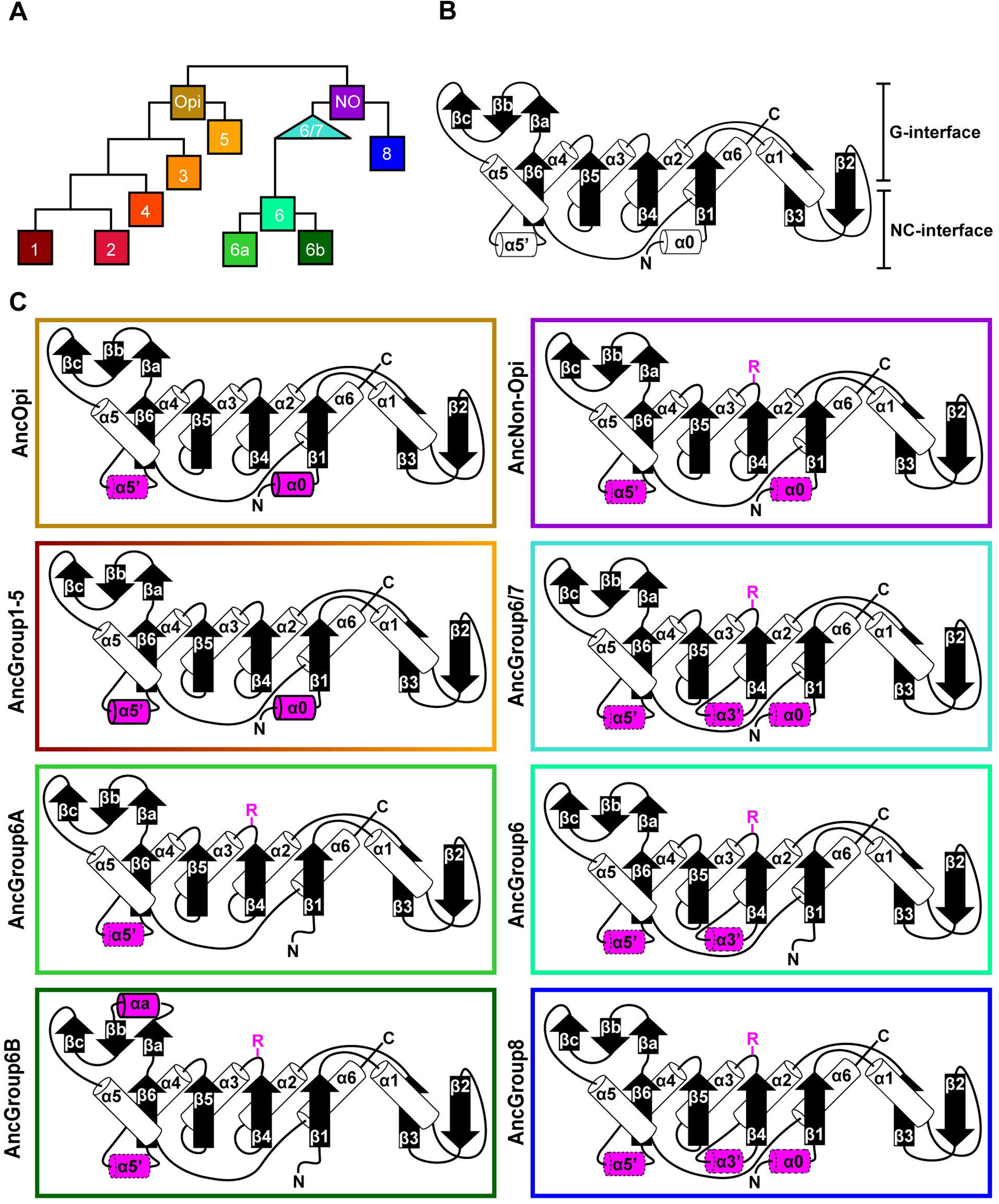
Ancestral sequence reconstruction of key evolutionary nodes throughout septin evolution. (**A**) Simplified tree diagram displaying the shape of the IQTree (Fig. S3) used in ancestral sequence reconstruction. Squares and triangle, key nodes with ancestral septins corresponding to interpretive diagrams shown in panel C. (**B**) Representative topology diagram of septin GTPase domain indicating both the G-interface and NC-interface. N and C represent the N-terminal and C-terminal end of the protein. α helices and β sheets are each numbered sequentially from the N- to C-termini, except for those in the SUE (βa-βc). (**C**) Interpretive topology diagrams of the reconstructed ancestral septins at the nodes labeled in panel A. Novel structural motifs found in this study are highlighted in magenta. Secondary structures outlined in bold solid lines and dotted lines represent motifs with higher (pLDDT >70) and lower (pLDDT <70) AlphaFold confidence scores, respectively. R, arginine finger.

To highlight gains and losses of sub-domain motifs during the evolution of ancestral septins, interpretive topology diagrams of the GTPase domains were generated based on the AlphaFold predictions (Fig. 4BC). This analysis revealed a largely consistent core structure of the GTPase domains consisting of six α-helices (α1-α6) and nine β-sheets (β1-β6 and βa-βc), as well as a few variable α-helices that emerged or were lost at specific ancestral nodes (see below). In addition to the helices and sheets, we identified an arginine residue positioned in the S3 motif of AncGroup 6-8 and LECA septins (Figs. 3BD and 4C). Although this residue is not found in the reconstructed in AncGroup 1-5 septins (Fig. 4C), some extant Group 5 septins, such as *A. nidulans* AspE, appear to have it (see below). Thus, this “R-finger” arginine is an ancestral feature of septin family proteins that has been lost in most opisthokonts. Intriguingly, it has been reported that this R-finger in the single septin of *C. reinhardtii* is required for its homo-dimerization across the G-interface (Pinto et al., 2017), where it reaches into the GTP-binding pocket of the opposite subunit to accelerate GTP hydrolysis (see Fig. 3C, G-interface). Thus, we suspected that the R-finger would invariably be conserved in single septins found in other species. This prediction was partially confirmed: 20 of the 23 single septins that were included in our analysis have an R-finger at the expected position (Fig. 5A), suggesting that the dimerization mechanism observed in *C. reinhardtii* may be ancestral and conserved in many algae and protists. Of the other three that lacked an R-finger, the sequence from the dinoflagellate *S. minutum* is an extremely large 4484-aa protein, with a septin-like domain near the N-terminus and some additional domains (e.g., SMC domain, HSP70) that are not found in other septins. The other two (from the ciliates *Halteria grandinella* and *Stylonychia lemnae*) have the arginine replaced by a histidine residue. It is unknown whether these single septins still form a G-dimer without an R-finger or have taken unique evolutionary paths to function without dimerizing through the G-interface.

**Figure 5.**
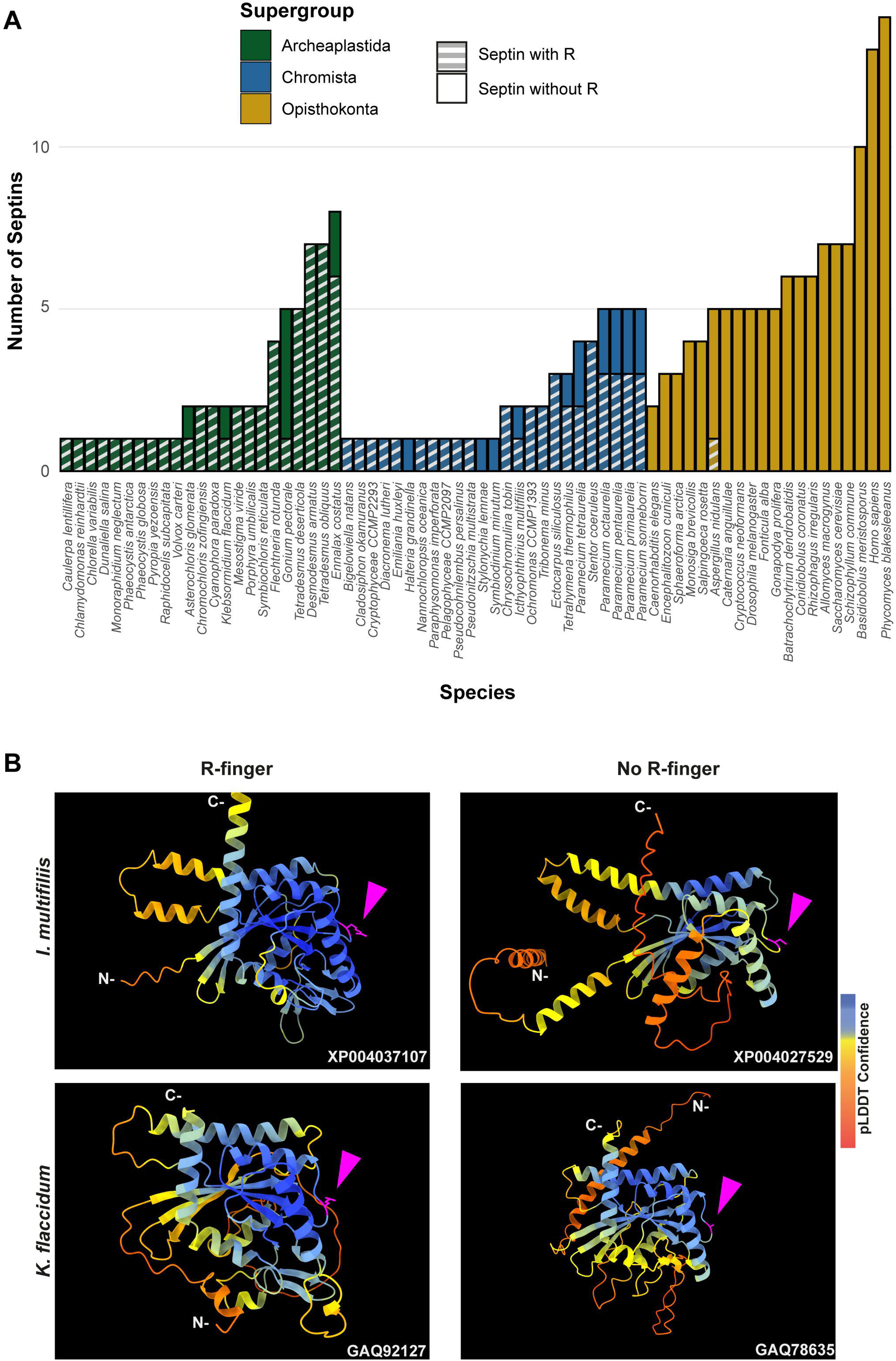
GAP-like R-finger is widely conserved in single septins. (**A**) Numbers of septins with and without R-finger in 68 species representing the three septin-harboring eukaryotic supergroups. (**B**) AlphaFold predictions of septins with and without R-finger in the species *I. multifiliis* (top row) and *K. flaccidum* (bottom row). N- and C-, amino-terminus and carbonyl-terminus, respectively. Magenta arrowheads indicate the positions with the presence or absence of R-finger. Structures are colored according to the AlphaFold pLDDT confidence scores.

Interestingly, in many algae and protists with multiple septin genes, a loss of the R-finger is observed in some of the duplicated genes (Fig. 5A). For example, the ciliate *Ichthyophthirius multifiliis* possesses two septins: XP004037107 with an R-finger and XP004027529 without (Fig. 5B). Similarly, the filamentous charophyte green alga *Klebsormidium flaccidum* has two proteins with and without an R-finger (GAQ92127 and GAQ78635, respectively; Fig. 5B). Given the apparent selective pressure against the loss of R-finger in single septins as well as the loss of R-finger in most opisthokont septins that are invariably encoded as multiple copies in a genome (see below), it is tempting to speculate that these septins may have lost their R-finger because of evolution to form hetero-oligomers. Biochemical characterization of these septins is needed to address this possibility.

Unlike the non-opisthokont counterparts, the vast majority of opisthokont septins do not possess an R-finger between the S2-S3 motifs (Figs. 3D and 4C). In Group 1-4 septins, the arginine residue is replaced by small uncharged amino acids such as serine, glycine, or alanine. Although there is an invariant histidine residue in the adjacent position (Fig. 3D) that could potentially be involved in GTP hydrolysis (Weirich et al., 2008), a mutation to this amino acid in human SEPT2 did not affect its GTPase activity (Sirajuddin et al., 2009). Thus, it is unlikely that the Group 1-4 opisthokont septins employ an R-finger-like molecular mechanism to interact through their G-interfaces. The R-finger is also absent in most filamentous-fungus-specific Group 5 septins (Figs. 3D and 4C), consistent with the previous observation that the S1-S4 motifs in septins in these groups are highly variable (Shuman and Momany, 2021). However, some septins, such as *Aspergillus nidulans* AspE (Fig. 5A), have an arginine residue located between the divergent S2-S3 motifs. Available data suggest that AspE is not incorporated into canonical septin complexes, although it interacts with them in a developmental-stage-specific manner (Hernandez-Rodriguez et al., 2014). It is interesting to speculate that AspE-type Group 5 septins have retained the ancestral trait to form a homomeric G-dimer using their R-fingers.

### Conservation of α0 and α5’ helices in opisthokont septins

In addition to the core helices and sheets, AncGroup 1-5 (opisthokont) septins displayed two additional invariant α-helices, both positioned in the NC-interface: α0 at the junction between the N-terminal extension and the GTPase domain, and α5’ that is positioned in-between α4 and β6 (Fig. 4C). Interestingly, however, these helices are not well conserved in Group 6-8 septins (Fig. 4C). In the human SEPT2/6/7 complex (and plausibly in many other opisthokont septins complexes), the α0 helix is an integral part of the NC interface where it forms an electrostatic inter-subunit interaction (Cavini et al., 2021). In addition, the α5’-helix contains a polyacidic region that is known to interact with the polybasic region 1 (PB1) within the α0 helix of a neighboring subunit across the NC interface (Fig. 3C; Cavini et al., 2021). Thus, it is conceivable that the α0 and α5’ helices evolved together in the opisthokont lineage as the positioning of PB1 was fixed in the former (see below).

The PB1 domain in α0 helix binds to phospholipids such as phosphatidylinositol 4-phosphate, 4,5-bisphosphate, and 3,4,5-triphosphate (Zhang et al., 1999; Casamayor and Snyder, 2003; Bertin et al., 2010; Onishi et al., 2010; Krokowski et al., 2018). The PB1 domain has been observed in some septins in non-opisthokont species such as in *C. reinhardtii* (Wloga et al., 2008; Nishihama et al., 2011; Pinto et al., 2017) despite the lack of α0 in the same proteins (Figs. 3D and 4B), raising the possibility that the emergence of PB1 precedes that of α0. To test this, we examined the NTEs of the reconstructed ASR sequences for the presence of PB1 by developing a Python script that calculates the isoelectric point of a 10 amino-acid window moving along protein sequences. We observed a basic region proximal to the beginning of the GTPase domain in AncGroup 1-5 septins (including in the very short NTE of AncGroup3 septin) (Fig. 6AB), consistent with the presence of PB1 in the majority of extant opisthokont septins (Nishihama et al., 2011; Shuman et al., 2021). Similarly, the regions immediately upstream of the G1 motif in AncGroup 6 and 6/7 septins are also highly basic (Fig. 6A). In contrast, the NTE of AncGroup8 is overall acidic (Fig. 6A), and a few basic residues found in this region are interdigitated by acidic residues (Fig. 6B), consistent with the reported ambiguity about the presence of polybasic regions in septins in *T. thermophila* and *P. tetraurelia* (Wloga et al., 2008). Interestingly, CLUSTALω alignment identified additional polybasic domains in AncGroup 6B and 6/7 septins at positions 339 and 214 aa upstream of the G1 motif, respectively, which exhibited higher homology to the proximal PB1 observed in AncGroup 1-5 septins (Fig. 6B), and the G1-proximal sequences (PB1’) are non-opisthokont-specific (Fig. 6B). Given the low overall sequence conservation of these regions in AncGroup 8 (Fig. 6B), it is not clear whether PB1’ is an ancestral feature that has been lost in opisthokont septins, or it was newly inserted adjacent to the G1 motif in the lineage leading to Group 6 and 7 septins. Overall, however, the presence of a polybasic region in the NTE appears to be an ancestral feature that predates the emergence of opisthokont-specific α0.

**Figure 6.**
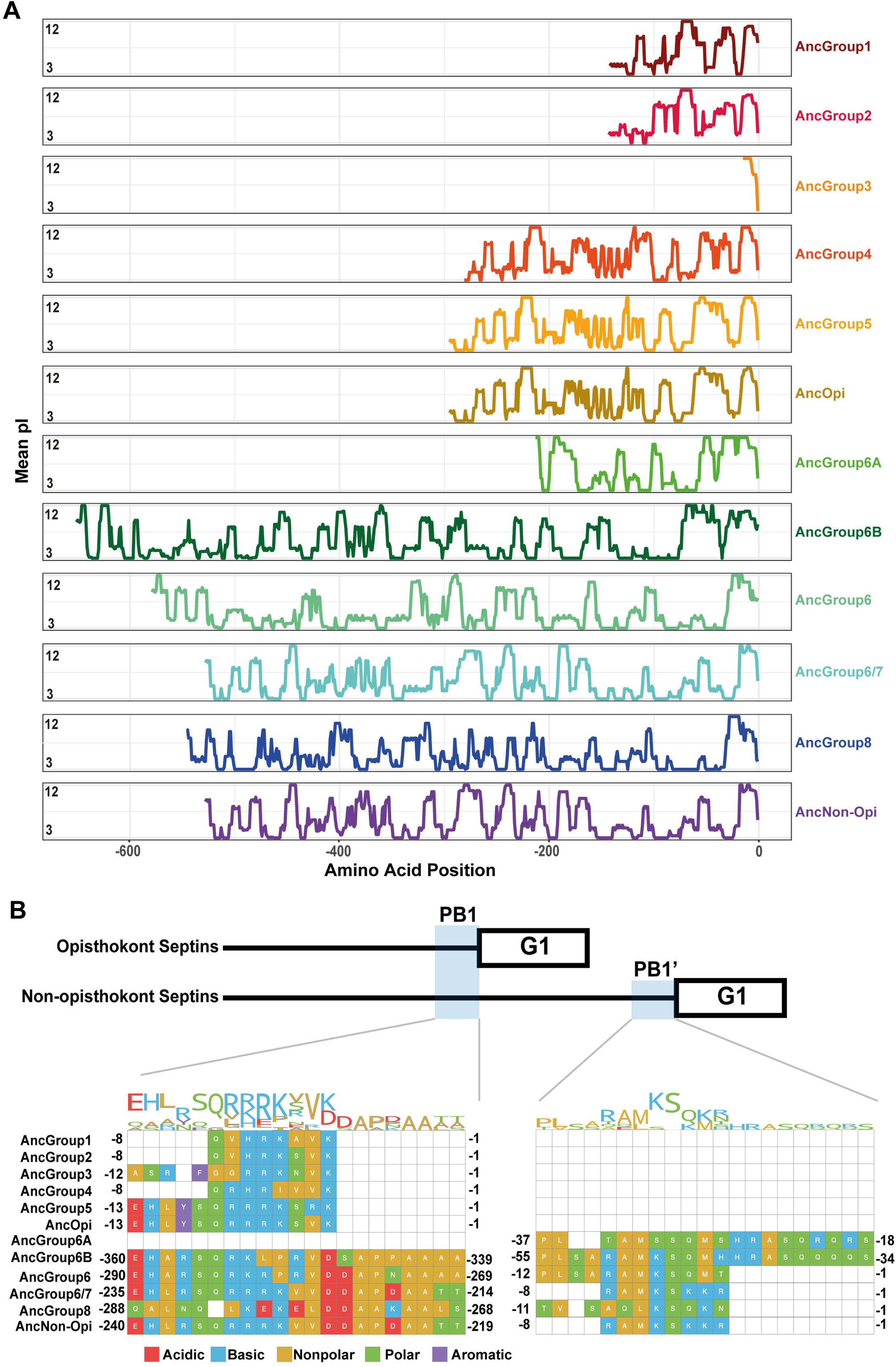
N-terminal polybasic domains across septins. (**A**) Calculation of isoelectric point windows across the NTE of reconstructed ancestral sequences. The average isoelectric point of a sliding 10 amino acid window is calculated across the NTE of reconstructed ancestral sequences. X=0 represents the start of the GTPase domain. (**B**) CLUSTALw multiple sequence alignment of reconstructed ancestral sequences displaying two polybasic domains in non-opisthokont lineages. Numbers indicate the amino acid positions from the start of the GTPase domain.

### Amphipathic helices are an ancestral feature of septins

Some opisthokont septins have the remarkable ability to recognize micron-scale membrane curvature through an amphipathic helix (AH) (Bridges et al., 2016; Cannon et al., 2019). Perturbation of these AHs can lead to abnormal subcellular localization of septin proteins (Cannon et al., 2019). To ask if AHs are found outside of opisthokonts and therefore can be an ancestral feature of septins, we developed a high-throughput pipeline to identify AH domains in a large number of polypeptide sequences by predicting alpha helices and then calculating their amphipathicity (see Materials & Methods), and applied it to the NTE and C-terminal extension (CTE) of our eukaryotic septin collection. This pipeline precisely identified previously reported AH domains in fungal and animal septins (Cannon et al., 2019; Lobato-Márquez et al., 2021; Woods et al., 2021), such as Cdc12 and Shs1 in *S. cerevisiae* and *Ashbya gossyppii*, human SEPT6, *Caenorhabditis elegans* UNC-61, and *Drosophila melanogaster* Sep1 (Fig. S5). In addition, our analysis revealed the presence of predicted AHs in septin sequences spanning all Groups (Fig. 7A; Table 2) with varying levels of conservation. In opisthokonts, for instance, predicted AHs were detected in 68% of Group 2 and Group 4 sequences, while only 13% of Group 3 sequences exhibited AHs. In Group 1, there is a striking difference between the two subclades: a predicted AH is completely absent in 1A (animals and fungi), while it is found in 75% of septins in 1B (animal-specific). This unexpected dichotomy suggests a potential connection between the evolution of AHs and the positioning of subunits within a canonical octameric protomer, in which 1A subunits occupy the central dimer. Like Group 3, only a small fraction of Group 5 septins (22%) have predicted AHs; unlike Group 1, there is no specific subgroup in which AHs are conserved, suggesting sporadic loss/gain of the domain within this group (Fig. 7A; Table 2). Notably, *A. nidulans* AspE has an unequivocal predicted AH with a large hydrophobic moment (Fig. 7E; Fig. S5), which may contribute to the highly cortical localization of this septin (Hernández-Rodríguez et al., 2014). In general, the AHs in Groups 1-5 displayed features consistent with stereotypical amphipathicity, with a large hydrophobic window and a hydrophilic face composed of both positively and negatively charged residues (Fig. S5).

**Figure 7.**
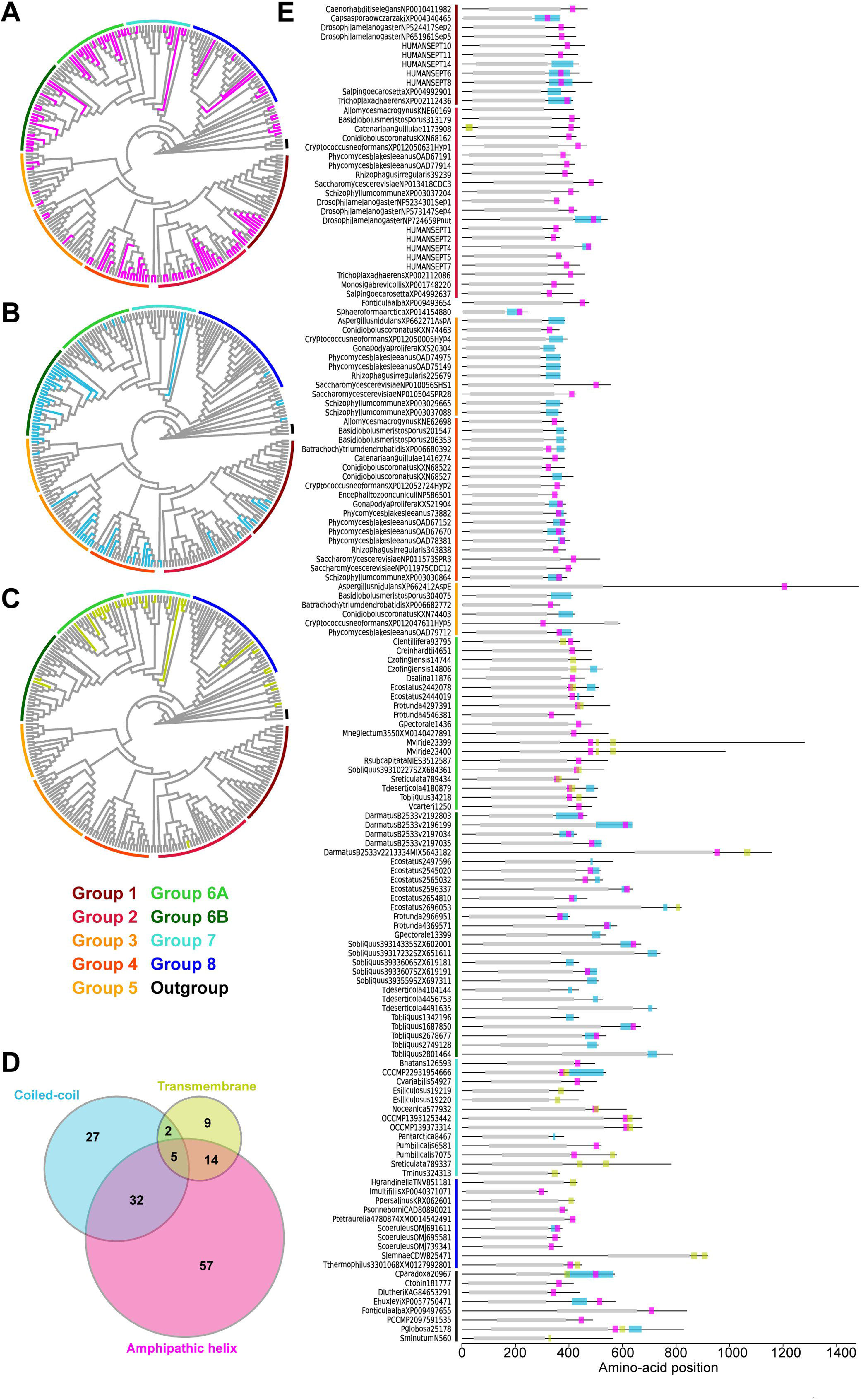
Distribution of AH, coiled-coil, and transmembrane domains across septin groups. (**A-C**) Simplified cladograms of the RAxML tree of 254 septins (see Fig. 2A), with individual sequences with AH (A, magenta), coiled-coil (B, blue), and transmembrane (C, green) domains highlighted. (**D**) Venn diagram showing the numbers of septins with AH, coiled-coil, and/or transmembrane domains. (**E**) Protein domain diagrams of septins with AH, coiled-coil, and/or transmembrane domains. Grey box, septin GTPase domain; magenta box, AH domain; blue box, coiled-coil domain; green box, transmembrane domain.

**Table 2.** Conservation of various features in septin groups.

| Group | Phylum | R-finger<br>(%) | $\alpha 0^a$ | PB1<sup>a</sup> | PB1'<sup>a</sup> | AH<br>(%)<sup>b</sup> | CC<br>(%)<sup>b</sup> | TM<br>(%)<sup>b</sup> |
| --- | --- | --- | --- | --- | --- | --- | --- | --- |
| 1 | Animals/fungi | 0 | Strong | Yes | No | 27<sup>c</sup> | 18 | 0 |
| 2 | Animals/fungi | 0 | Strong | Yes | No | **68** | 6.5 | 3.2 |
| 3 | Fungi | 0 | Strong | Yes | No | 13 | **33** | 0 |
| 4 | Fungi | 0 | Strong | Yes | No | **68** | **46** | 0 |
| 5 | Filamentous fungi | 5.6 | Strong | Yes | No | 22 | 17 | 0 |
| 6A | Green algae | 100 | None | No | Yes | **68** | 16 | **44** |
| 6B | Green algae | 80 | None | Yes | Yes | **50** | **87** | 6.7 |
| 7 | Various algae | 91 | Weak | Yes? | Yes?<sup>d</sup> | **38** | 9.5 | **43** |
| 8 | Ciliates | 60 | Weak | No | No | 18.9 | 2.7 | 11 |

<sup>a</sup>Based on AlphaFold predictions of ancestral protein structures.
<sup>b</sup>Based on analyses of extant sequences. Values greater than 30 are bold-faced. See Supplementary File 8 for details.
<sup>c</sup>0% in 1A, 75% in 1B.
<sup>d</sup>Because Group 7 is paraphyletic, we could not confidently infer the conservation of PB domains based on AngGroup6/7.

The wide distribution of AHs is also observed in all non-opisthokont groups (Fig. 7A; Table 2). Group 6A, consisting largely of single septins, has the highest rate of AH domains at 68%. In Group 6B, septins with predicted AHs were found in most subclades, with a total preservation rate of 50%. In Groups 7 and 8, septins with predicted AHs were found in at 38% and 19%, respectively. In the Heliquest visualization, both AHs present in Group 6B and Group 7 exhibited hydrophilic faces primarily composed of positively charged residues interspersed with small polar residues such as serines and threonines (Fig. S5). In some instances, weaker amphipathic helices were observed, as exemplified by *P. umbilicalis* 6581, which lacked a strongly pronounced hydrophilic face but still fulfilled the criteria of our search because of their high net charges that raised the *D-*factor (Fig. 7E; Fig. S5). Some Group 6A and Group 8 septins have predicted AHs similar to those observed in Groups 1-5 with a large hydrophobic window opposite the cluster of both positively and negatively charged residues.

### Selective distribution of coiled-coil and transmembrane domains in specific septin groups

Many animal and fungal septins contain a coiled-coil (CC) motif in the CTE which is thought to be involved in polymer stabilization and the formation of bundles and filament pairs (Sirajuddin et al., 2007; Bertin et al., 2010; Cavini et al., 2021). We utilized the existing annotation of coiled-coil domains in the Uniprot database to identify them in our list of 254 extant septins. Interestingly, we observed the presence of CCs in Groups 1B, 3, 4, and 6B (Fig 7B; Table 2). The majority of these sequences were also positive for AH domains (Fig. 7D), with AH domains residing within CC domains in many cases, such as in *S. cerevisiae* Cdc12 (Fig 7E; Cannon et al., 2019). Interestingly, CC domains were almost entirely excluded from non-opisthokont Groups 6A, 7, and 8 (Fig. 7B; Table 2), suggesting that the CC domains observed in Group 6B were a result of convergent molecular evolution. It is interesting to speculate that septin gene duplication in some green algae (Fig. 5A) and the formation of heterooligomeric complexes may have led to the emergence of lateral pairing between septin subunits.

Lastly, it has previously been reported that some non-opisthokont septins possess putative transmembrane (TM) domains or short hydrophobic patches (Wloga et al., 2008; Nishihama et al., 2011). Thus, we searched for the presence of potential TM domains in our list of 254 extant septin sequences. Except for one sequence from the parasitic fungus *Catenaria anguillulae* (A0A1Y2I4M7, Group 2A, 46% identical to *S. cerevisiae* Cdc3) that has a unique N-terminal TM domain, all septins with a TM domain were found in the non-opisthokont lineages, with notable enrichment in Groups 6A and 7 (Fig. 7CE; Table 2). This distribution of TM domains in our dataset seems to suggest that they emerged early in the non-opisthokont branch after its split with opisthokonts and were subsequently lost in many species in Group 6B and 8. [See, however, Discussion for another possibility given a recent report by (Perry et al., 2023).] It is interesting to note that there is little overlap between the distributions of CC and TM domains in Group 6 septins (Fig. 7D), perhaps suggesting that the evolution of septin-septin interactions through CC domains necessitated a concomitant loss of TM that would otherwise restrict the accessibility of CTE.

In summary, our searches for α-helix-based structures that are often associated with septin CTE suggest that the AH and TM domains may have ancient origins in septin evolution, while the CC domain may have evolved independently in multiple lineages.

## DISCUSSION

Septins have been reported in a variety of eukaryotic lineages outside of opisthokonts (Versele and Thorner, 2005; Wloga et al., 2008; Nishihama et al., 2011; Yamazaki et al., 2013; Onishi and Pringle, 2016; Shuman and Momany, 2021), although their phylogenetic relationships have not been fully explored. Here, we performed an updated search for septins in non-opisthokont lineages and found that septins are widely spread in two distinct non-opisthokont eukaryotic supergroups: Archaeplastida and Chromista. Because these two supergroups and opisthokonts share the ancestry only at the LECA level, our results strongly support the idea that the first septin appeared in an early eukaryotic ancestor. We inferred structural features related to septin-septin interactions, membrane binding, and curvature sensing across eukaryotic evolution, and hypothesized functions related to ancestral septins.

Septins in Archaeplastida and Chromista form new phylogenetic clades outside of the previously defined Groups 1-5, herein named Groups 6A, 6B, 7, and 8. Group 6A and 6B are composed exclusively of septins from various green algae, while septins in Groups 7 and 8 belong to other various algae (some other green algae, red algae, heterokonts, haptophytes, cryptophytes, chlorarachniophytes) and ciliates, respectively. It is peculiar that these septins in algae from diverse groups formed a single clade separate from the ciliate septins, which is inconsistent with the general taxonomical classification of these species (compare Fig. 1 and Fig. 2A). It is tempting to speculate that these algal septins may have spread through horizontal transfer of nuclear genes, when ancestral red and green algae were taken up by other eukaryotes to form secondary and tertiary endosymbiosis (Keeling and Palmer, 2008; Archibald, 2012).

In this study, we found that the majority (but not all) of non-opisthokont septins have a conserved arginine residue within the G-interface. This arginine is predicted to act similarly to other R-fingers in GTPase-activating proteins (GAPs). Because R-fingers are also found in other “paraseptin” GTPases such as TOC34/TOC159 and AIG1/GIMAP (Leipe et al., 2002; Weirich et al., 2008), it is likely an ancestral feature that has been lost in some lineages. Biochemical and structural studies on the single Group 6A septin from *C. reinhardtii* have shown that this arginine is critical for the very high GTPase activity of this septin (40 times higher than human SEPT9, the most active septin GTPase in opisthokonts) and its homo-dimerization through the G-interface (Pinto et al., 2017). Interestingly, while Group 6A septins invariably have an R-finger, some Group 6B septins have lost this residue. It appears that the loss of R-finger is a crucial evolutionary step associated with septin gene duplication in many eukaryotic lineages, including Group 6 (green algae), Group 8 (ciliates), and the transition from ancestral septin to opisthokonts.

Suppose we imagine an ancestral septin dimer with subunits possessing two potential interaction interfaces (G and NC). In that case, we predict that the presence of an R-finger strongly biases the interaction to the G-interface, suggesting that most ancestral septins formed a dimer across their G-interface. Upon gene duplication, some septins lost the R-finger and gained the NC-interface interaction motif, α0. These evolutionary events then would shift the equilibrium to favor the NC-interface, allowing for the formation of septin heterocomplex protomers. In some cases, evolution of non-opisthokont septin complexes may have involved further mutations in the GTP-binding pocket and the G-interface, causing some septins to be locked in apo-nucleotide or GTP-bound state, as seen in some opisthokont septins (Hussain et al., 2023).

When hypothesizing about the potential ancestral functions of septins, we sought to identify motifs that are crucial for septin function. We observe the presence of a polybasic domain immediately preceding the GTPase domain in all septins except for Group 8. Previous studies have implicated this domain to be important for membrane recognition, as well as stabilizing an NC-interaction interface (Bertin et al., 2010; Cavini et al., 2021). The wide distribution of the polybasic domain, but not an α0 helix in which it is found in opisthokonts, suggests that the role of ancestral septins involved their binding to lipid bilayers. In support of this, we found that AH domains were also present across many of the septin phylogenetic groups, suggesting that they are also an ancestral septin feature. By comparing helical wheel diagrams of these AH domains across species, we begin to see some level of heterogeneity in the amino acid composition. Models to distinguish curvature sensing peptides highlight the importance of specific amino acid composition in either being a membrane sensor versus a membrane binder (van Hilten et al., 2023). It could be that the variation in amino acid composition confers distinct membrane binding properties, such as curvature sensing or subcellular localization. Within Groups 1-5, AH domains often had large hydrophobic faces and a large hydrophobic moment due to the presence of acidic and basic residues along the hydrophilic face. In contrast, in some lineages, particularly in group 6B and group 7, we observe the reduction of charged residues and often find threonine and serine residues. These residues may act as potential phosphorylation sites to adaptively regulate the functional properties of these helices (Byeon et al., 2022). Future biochemical studies of the AH domains of diverse septins would provide additional context to the ancestral role of this domain.

We identified the presence of CC and putative TM domains in the CTE of septins across various phylogenetic groups. In non-opisthokonts, we observed an almost exclusive and ubiquitous conservation of CC domains in Group 6B, while TM domains are highly enriched in Group 6A. Considering that Group 6B is composed of septins that have undergone recent gene duplication, it raises an interesting possibility that septins utilize CC to form interactions between subunits and filaments only after the emergence of heterocomplexes. In this scenario, gene duplication and subsequent diversification would be a prerequisite for this specialization of function among subunits. It is important to note that our classification of septin groups was based solely on the sequences of the GTPase domain, independently of the CTE sequence. Therefore, the strong correlation between Group 6A/TM and Group6B/CC suggests a co-evolution between the GTPase and CTE.

In addition to Group 6, TM domains were found sporadically in the CTE of some Group 7 and 8 septins but largely missing from the opisthokont sequences we used in our analysis. We initially interpreted this as evidence that the TM domain emerged after the opisthokont/non-opisthokont split and was subsequently lost in some lineages. However, a recent study by Perry et al. (2023) reported the presence of TM domains in a transcript isoform of *C. elegans UNC-61* (Group 1) as well as many other opisthokont proteins currently annotated as septins on the Uniprot database (but were not included in our list of 254 septins). Interestingly, many of these TM domains are found in the NTE, as seen in *C. anguillulae* A0A1Y2I4M7 (Fig. 7E). Thus, we provide two possible interpretations: The N- and C-terminal TM domains evolved independently in opisthokonts and non-opisthokonts, respectively. Alternatively, the LECA septin possessed a TM in the C-terminus, which was inherited by some progeny in all septin groups; in opisthokonts, domain movement within a gene (Furuta et al., 2011) shifted the position of TM from C- to N-terminus.

For future studies of septin evolution and general principles of evolutionary constraints, two approaches appear particularly appealing. First, a comparative approach using green algae with single vs. multiple septins seems to provide a unique opportunity to understand the evolution of septin duplication and the formation of heterocomplexes. For example, while *C. reinhardtii* possesses a single Group 6A septin with R-finger, PB1/PB1’, AH, and possible TM ((Wloga et al., 2008; Nishihama et al., 2011); though it is not currently annotated as such on Uniprot), a related green alga in the same Chlamydomonadales order, *G. pectorale*, has a total of five septins (one Group 6A and four 6B) with various combinations of septin features (Supplementary File 8). The Kinoshita rule (Kinoshita, 2003a) of opisthokont septins highlights the modularity and redundancy of opisthokont septin subunits at each position of a canonical protomer, where a septin from the same group can replace one another. Biochemical and cell biological experiments of Group 6A and Group 6B septins can shed light on whether this rule also applies to non-opisthokont septins.

Second, to understand how – parsimoniously – a single septin with R-finger evolved into a highly variable family of five septin groups in opisthokonts, some filamentous fungi possessing Group 5 with putative R-fingers seem to be an ideal model. One such protein, AspE in *A. nidulans*, has been shown to be excluded from the heterooligomeric complex formed by other subunits (Hernández-Rodríguez et al., 2014). Perhaps this septin has an extremely high GTPase activity, forms a G-dimer, and works independently of canonical filaments or binds to filaments in a substoichiometric fashion.

Finally, although our study provided a general overview of septin evolution, it is important to consider these evolutionary events in the context of the cellular processes the ancestral septins were involved in. Given the near-universal role of animal and fungal septins in cytokinesis, it is tempting to speculate that ancestral septins had similar roles. In support of this, the single septin in the green alga *N. bacillaris* showed its localization at the division site (Yamazaki et al., 2013). However, the two and only other reports on non-opisthokont septins did not show division-site localization: in another green alga *C. reinhardtii,* a septin was found at the flagella-base region, and in the ciliate *T. thermophila,* septins were found associated with mitochondria (Wloga et al., 2008; Pinto et al., 2017). Further functional studies of septins in non-opisthokonts are necessary to reveal the ancestral and fundamental functions of septins.

## ACKNOWLEDGMENTS

We thank Jenna Perry and Amy Maddox for sharing unpublished information. This work was supported by National Institutes of Health Grant R01 GM131004 (to S.B.), National Science Foundation Grant MCB 1818383 (to M.O. and John R. Pringle), Duke University Department of Biology, and The Franklin College of Arts and Sciences at the University of Georgia.

---

**Supplementary Figure 1.**
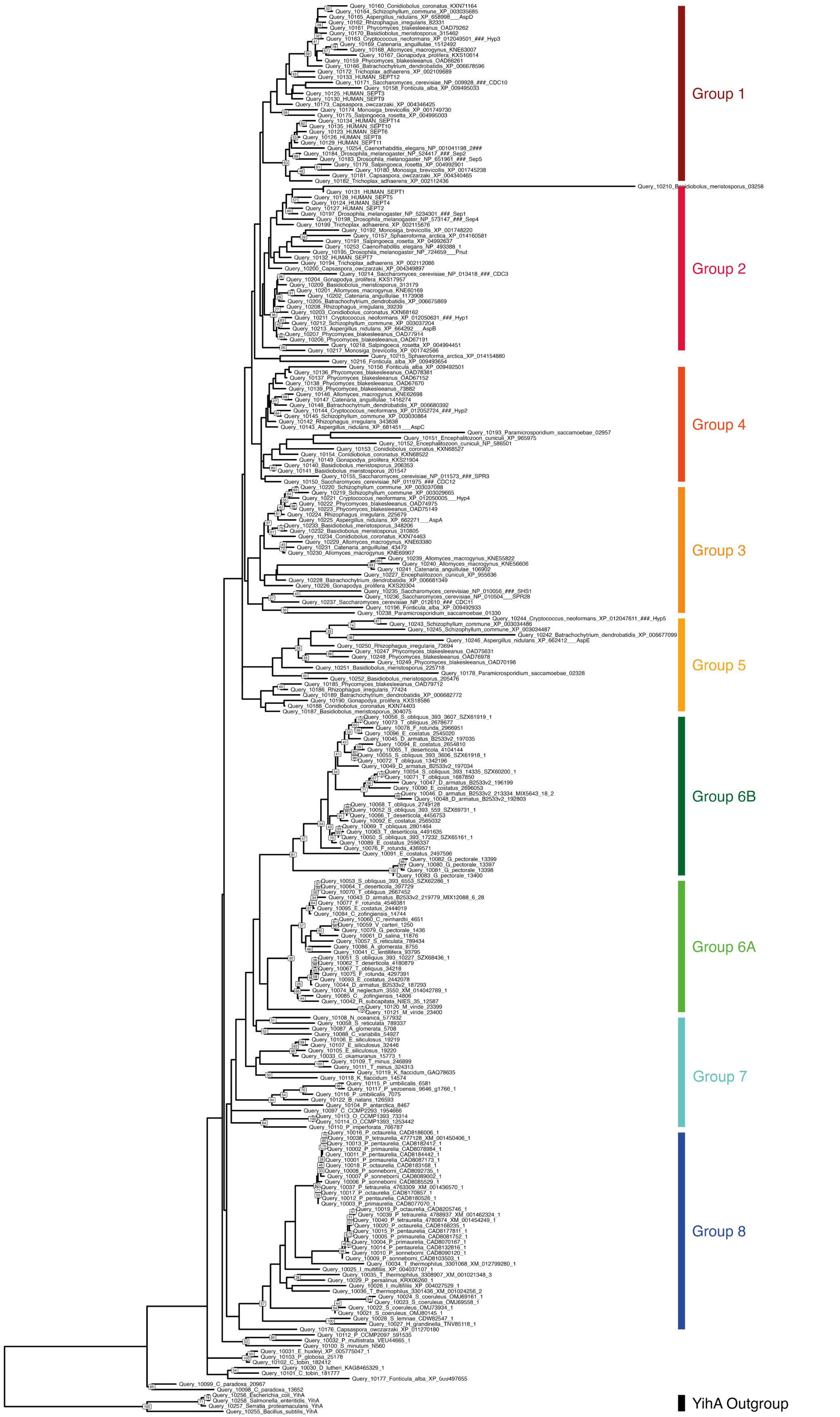
RAxML tree of all eukaryotic septins with 1000 bootstraps and YihA family as outgroup. Bootstrap values <25 are not shown. Defined phylogenetic groups are colored and displayed adjacent to tree tips.

**Supplementary Figure 2.**
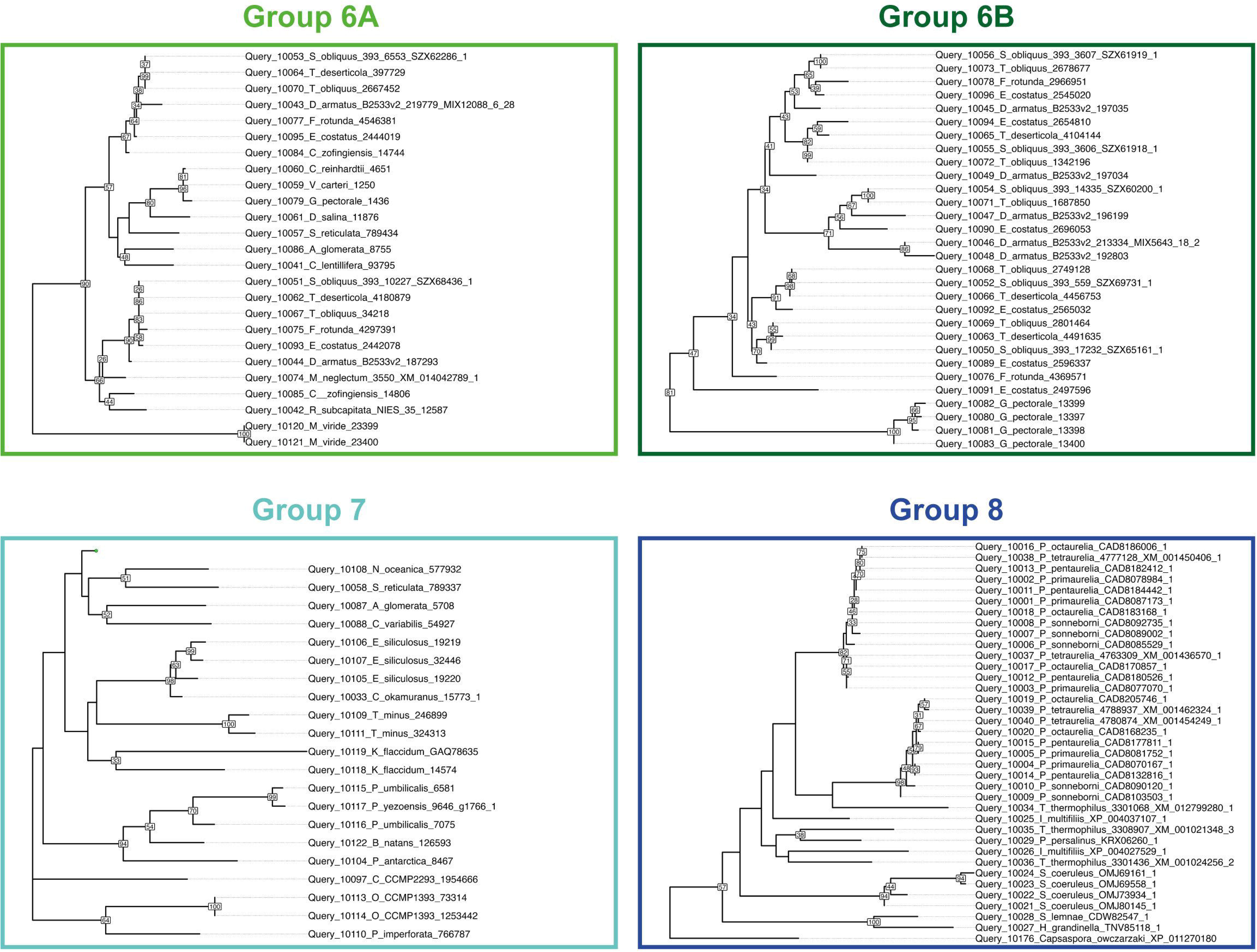
Magnified views of septin groups 6-8.

**Supplementary Figure 3.**
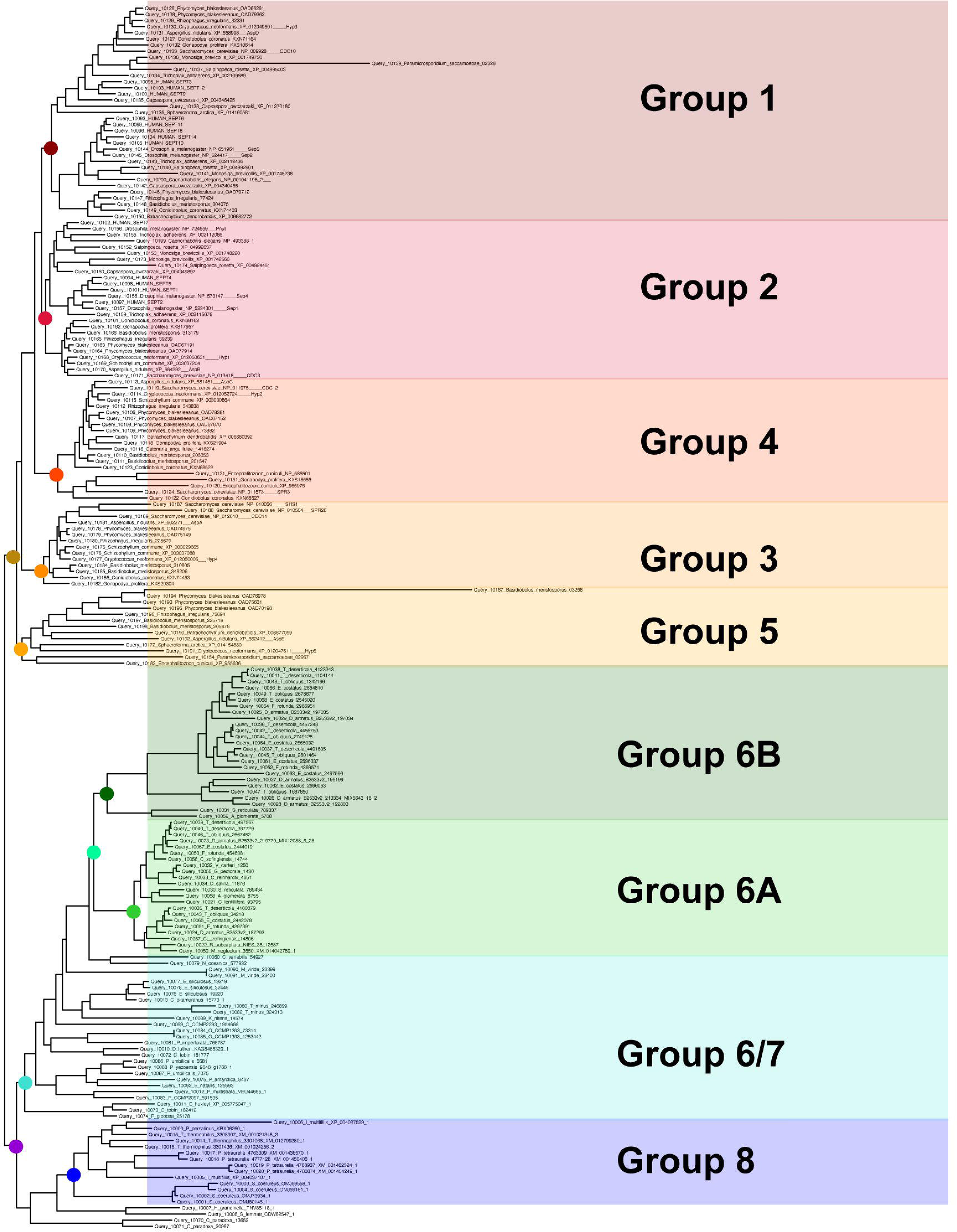
IQTree tree 200 septin sequences used in ancestral sequence reconstitution. Groups as defined in Figure 2 are redefined adjacent to branch tips. Colored nodes represent select ancestral sequences used for AlphaFold prediction.

**Supplementary Figure 4.**
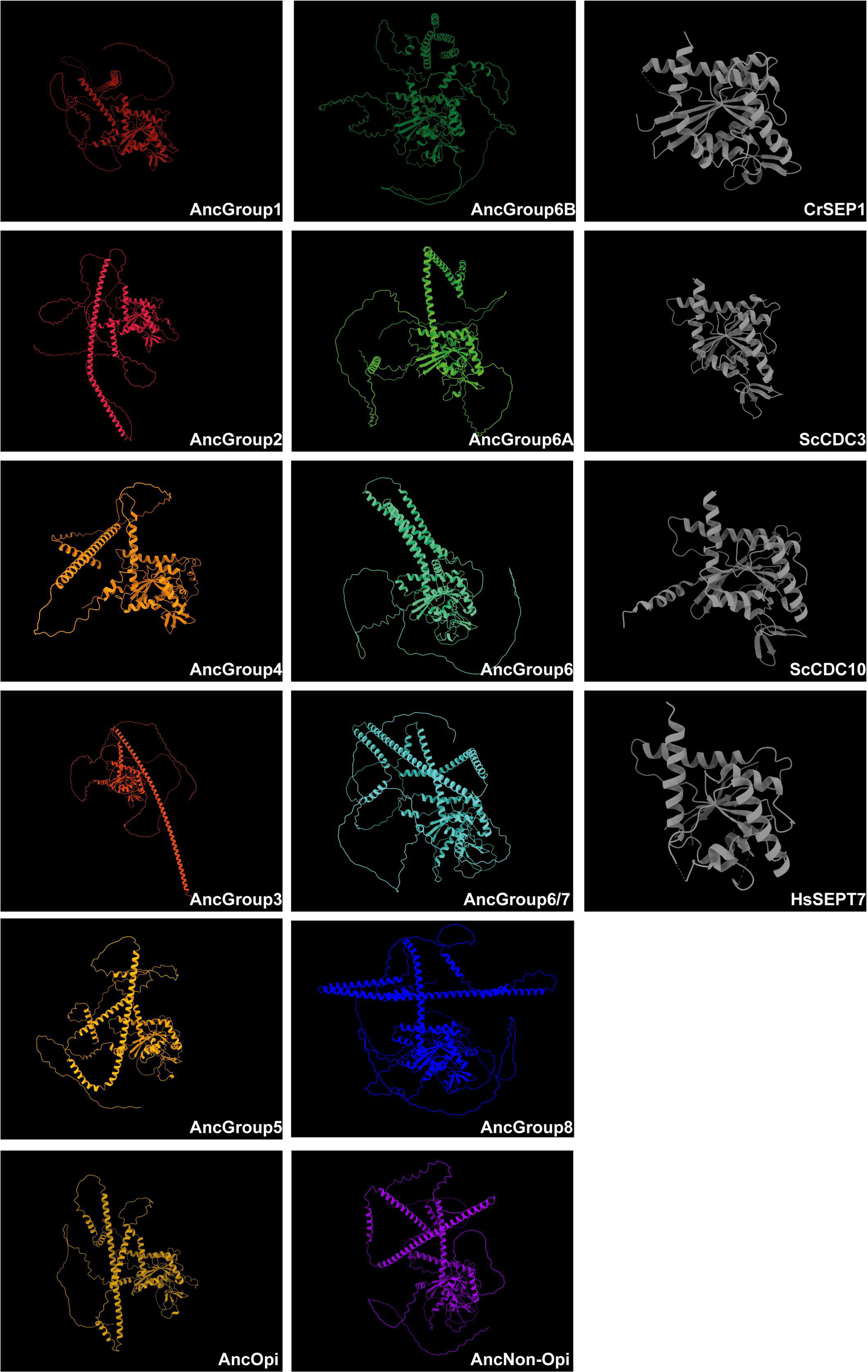
AlphaFold-predicted 3D structures of ancestral septins. Structures in grey are experimentally determined Protein Data Bank (PDB) files of septin GTPase domains, included here as references. Structures are orientated such that the NC-interface is towards the left of the monomer and the G-interface is towards the right.

**Supplementary Figure 5.**
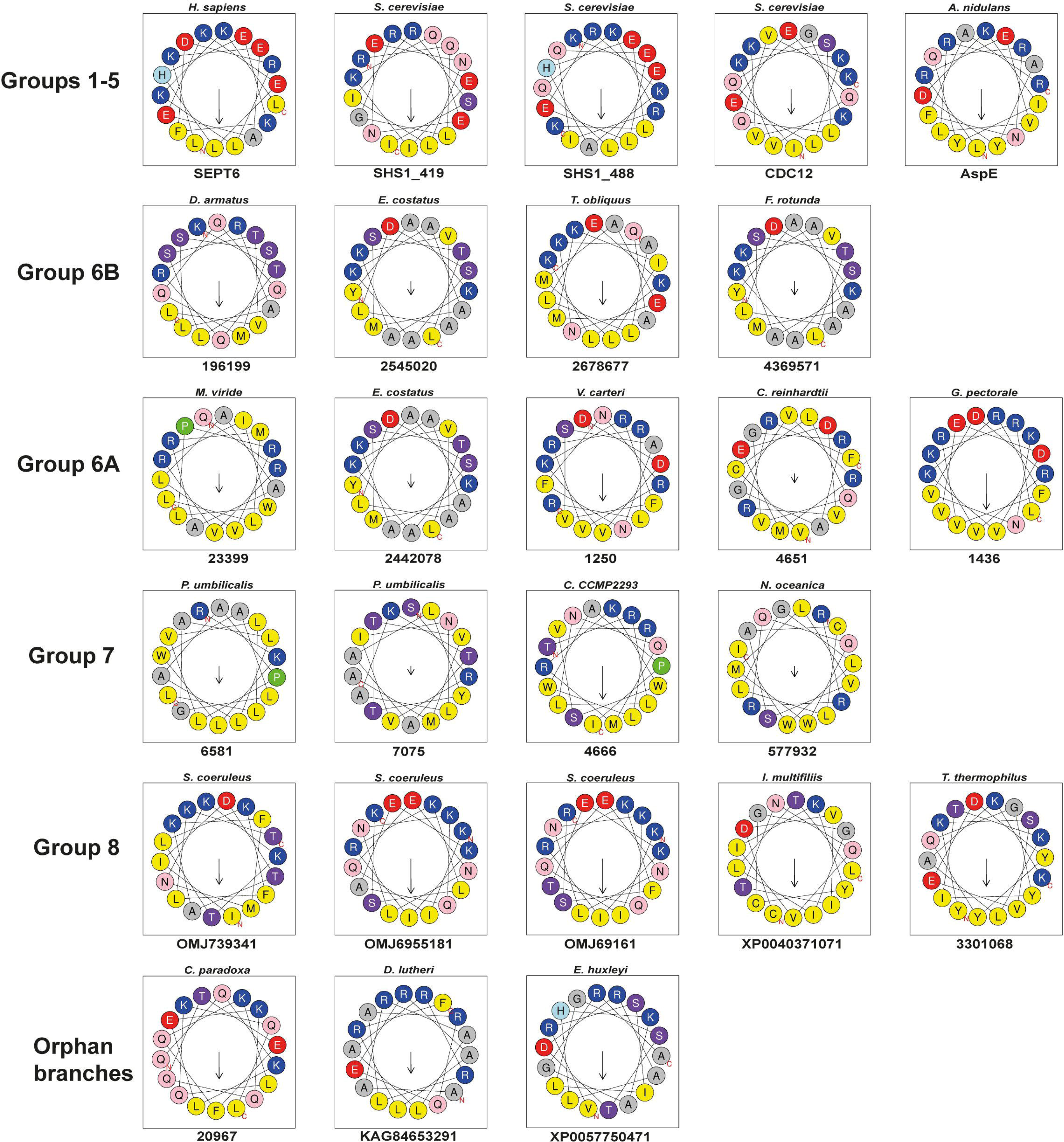
Representative helical wheel diagrams of predicted AHs across septin phylogenetic groups. Arrow represents the hydrophobic moment vector. Amino acids are colored according to their chemistry: yellow, hydrophobic; purple, Ser/Thr residues; grey, Gly/Ala residues; blue, basic residues; red, acidic residues; pink, Asp; green, Pro.

## Supplementary File/Data

**Supplementary File 1.** Query Septin FASTA sequences used in this study.

**Supplementary File 2.** List of Searched JGI Genomes + NCBI Taxa.

**Supplementary File 3.** Septin + YihA Fasta File.

**Supplementary File 4**. ALISCORE & ALICUT Processed File.

**Supplementary File 5**. IQTREE Input and Output files.

**Supplementary File 6**. AlphaFold Prediction Files.

**Supplementary File 7**. AH 18-aa Raw Output.

**Supplementary File 8**. Table summarizing properties of septins used in this study (Phylogenetic group, R-finger, AH, CC, TM).

## REFERENCES

Archibald, J. M. (2012). “Chapter Three - The Evolution of Algae by Secondary and Tertiary Endosymbiosis,” in Advances in Botanical Research, ed. G. Piganeau (Academic Press), 87–118. doi: 10.1016/B978-0-12-391499-6.00003-7

Ashkenazy, H., Penn, O., Doron-Faigenboim, A., Cohen, O., Cannarozzi, G., Zomer, O., et al. (2012). FastML: a web server for probabilistic reconstruction of ancestral sequences. Nucleic Acids Research 40, W580–W584. doi: 10.1093/nar/gks498

Auxier, B., Dee, J., Berbee, M. L., and Momany, M. (2019). Diversity of opisthokont septin proteins reveals structural constraints and conserved motifs. BMC Evol Biol 19, 4. doi: 10.1186/s12862-018-1297-8

Bertin, A., McMurray, M. A., Thai, L., Garcia, G., Votin, V., Grob, P., et al. (2010). Phosphatidylinositol-4,5-bisphosphate Promotes Budding Yeast Septin Filament Assembly and Organization. Journal of Molecular Biology 404, 711–731. doi: 10.1016/j.jmb.2010.10.002

Bridges, A. A., Jentzsch, M. S., Oakes, P. W., Occhipinti, P., and Gladfelter, A. S. (2016). Micron-scale plasma membrane curvature is recognized by the septin cytoskeleton. J Cell Biol 213, 23–32. doi: 10.1083/jcb.201512029

Byeon, S., Werner, B., Falter, R., Davidsen, K., Snyder, C., Ong, S.-E., et al. (2022). Proteomic Identification of Phosphorylation-Dependent Septin 7 Interactors that Drive Dendritic Spine Formation. Frontiers in Cell and Developmental Biology 10. Available at: https://www.frontiersin.org/articles/10.3389/fcell.2022.836746 (Accessed December 28, 2023).

Byers, B., and Goetsch, L. (1976). A highly ordered ring of membrane-associated filaments in budding yeast. Journal of Cell Biology 69, 717–721. doi: 10.1083/jcb.69.3.717

Cannon, K. S., Woods, B. L., Crutchley, J. M., and Gladfelter, A. S. (2019). An amphipathic helix enables septins to sense micrometer-scale membrane curvature. Journal of Cell Biology 218, 1128–1137. doi: 10.1083/jcb.201807211

Casamayor, A., and Snyder, M. (2003). Molecular dissection of a yeast septin: distinct domains are required for septin interaction, localization, and function. Mol Cell Biol 23, 2762– 2777. doi: 10.1128/MCB.23.8.2762-2777.2003

Castro, D. K. S. V., Rosa, H. V. D., Mendonça, D. C., Cavini, I. A., Araujo, A. P. U., and Garratt, R. C. (2023). Dissecting the Binding Interface of the Septin Polymerization Enhancer Borg BD3. Journal of Molecular Biology 435, 168132. doi: 10.1016/j.jmb.2023.168132

Cavalier-Smith, T. (2018). Kingdom Chromista and its eight phyla: a new synthesis emphasising periplastid protein targeting, cytoskeletal and periplastid evolution, and ancient divergences. Protoplasma 255, 297–357. doi: 10.1007/s00709-017-1147-3

Cavini, I. A., Leonardo, D. A., Rosa, H. V. D., Castro, D. K. S. V., D’Muniz Pereira, H., Valadares, N. F., et al. (2021). The Structural Biology of Septins and Their Filaments: An Update. Frontiers in Cell and Developmental Biology 9. Available at: https://www.frontiersin.org/articles/10.3389/fcell.2021.765085 (Accessed May 29, 2023).

Eisenberg, D., Weiss, R. M., and Terwilliger, T. C. (1982). The helical hydrophobic moment: a measure of the amphiphilicity of a helix. Nature 299, 371–374. doi: 10.1038/299371a0

Fauchere, J., and Pliska, V. (1983). Hydrophobic parameters II of amino acid side-chains from the partitioning of N-acetyl-amino acid amides. Eur. J. Med. Chem. 18.

Feng, S.-H., Xia, C.-Q., and Shen, H.-B. (2022). CoCoPRED: coiled-coil protein structural feature prediction from amino acid sequence using deep neural networks. Bioinformatics 38, 720–729. doi: 10.1093/bioinformatics/btab744

Field, C. M., al-Awar, O., Rosenblatt, J., Wong, M. L., Alberts, B., and Mitchison, T. J. (1996). A purified Drosophila septin complex forms filaments and exhibits GTPase activity. The Journal of cell biology 133, 605–616. doi: 10.1083/jcb.133.3.605

Furuta, Y., Kawai, M., Uchiyama, I., and Kobayashi, I. (2011). Domain movement within a gene: a novel evolutionary mechanism for protein diversification. PLoS One 6, e18819. doi: 10.1371/journal.pone.0018819

Gautier, R., Douguet, D., Antonny, B., and Drin, G. (2008). HELIQUEST: a web server to screen sequences with specific alpha-helical properties. Bioinformatics 24, 2101–2102. doi: 10.1093/bioinformatics/btn392

Grupp, B., and Gronemeyer, T. (2023). A biochemical view on the septins, a less known component of the cytoskeleton. Biological Chemistry 404, 1–13. doi: 10.1515/hsz-2022-0263

Hartwell, L. H. (1971). Genetic control of the cell division cycle in yeast: IV. Genes controlling bud emergence and cytokinesis. Experimental Cell Research 69, 265–276. doi: 10.1016/0014-4827(71)90223-0

Hartwell, L. H., Culotti, J., Pringle, J. R., and Reid, B. J. (1974). Genetic Control of the Cell Division Cycle in Yeast. Science 183, 46–51. doi: 10.1126/science.183.4120.46

Hernández-Rodríguez, Y., Masuo, S., Johnson, D., Orlando, R., Smith, A., Couto-Rodriguez, M., et al. (2014). Distinct Septin Heteropolymers Co-Exist during Multicellular Development in the Filamentous Fungus Aspergillus nidulans. PLOS ONE 9, e92819. doi: 10.1371/journal.pone.0092819

Hussain, A., Nguyen, V. T., Reigan, P., and McMurray, M. (2023). Evolutionary degeneration of septins into pseudoGTPases: impacts on a hetero-oligomeric assembly interface. Front Cell Dev Biol 11, 1296657. doi: 10.3389/fcell.2023.1296657

Jumper, J., Evans, R., Pritzel, A., Green, T., Figurnov, M., Ronneberger, O., et al. (2021). Highly accurate protein structure prediction with AlphaFold. Nature 596, 583–589. doi: 10.1038/s41586-021-03819-2

Käll, L., Krogh, A., and Sonnhammer, E. L. L. (2004). A combined transmembrane topology and signal peptide prediction method. J Mol Biol 338, 1027–1036. doi: 10.1016/j.jmb.2004.03.016

Keeling, P. J., and Palmer, J. D. (2008). Horizontal gene transfer in eukaryotic evolution. Nat Rev Genet 9, 605–618. doi: 10.1038/nrg2386

Kinoshita, M. (2003a). Assembly of mammalian septins. J Biochem 134, 491–496. doi: 10.1093/jb/mvg182

Kinoshita, M. (2003b). The septins. Genome Biol 4, 1–9. doi: 10.1186/gb-2003-4-11-236

Koenig, P., Oreb, M., Rippe, K., Muhle-Goll, C., Sinning, I., Schleiff, E., et al. (2008). On the Significance of Toc-GTPase Homodimers*. Journal of Biological Chemistry 283, 23104–23112. doi: 10.1074/jbc.M710576200

Krogh, A., Larsson, B., von Heijne, G., and Sonnhammer, E. L. (2001). Predicting transmembrane protein topology with a hidden Markov model: application to complete genomes. J Mol Biol 305, 567–580. doi: 10.1006/jmbi.2000.4315

Krokowski, S., Lobato-Márquez, D., Chastanet, A., Pereira, P. M., Angelis, D., Galea, D., et al. (2018). Septins Recognize and Entrap Dividing Bacterial Cells for Delivery to Lysosomes. Cell Host & Microbe 24, 866–874.e4. doi: 10.1016/j.chom.2018.11.005

Kück, P., Meusemann, K., Dambach, J., Thormann, B., von Reumont, B. M., Wägele, J. W., et al. (2010). Parametric and non-parametric masking of randomness in sequence alignments can be improved and leads to better resolved trees. Frontiers in Zoology 7, 10. doi: 10.1186/1742-9994-7-10

Kueck, P. (2017). ALICUT: a Perlscript which cuts ALISCORE identified RSS. Available at: https://github.com/PatrickKueck/AliCUT/blob/master/ALICUT_V2.31.pl

Lee, S., Carrasquillo Rodrı Guez, J. W., Merta, H., and Bahmanyar, S. (2023). A membrane-sensing mechanism links lipid metabolism to protein degradation at the nuclear envelope. J Cell Biol 222, e202304026. doi: 10.1083/jcb.202304026

Leipe, D. D., Wolf, Y. I., Koonin, E. V., and Aravind, L. (2002). Classification and evolution of P-loop GTPases and related ATPases11Edited by J. Thornton. Journal of Molecular Biology 317, 41–72. doi: 10.1006/jmbi.2001.5378

Leonardo, D. A., Cavini, I. A., Sala, F. A., Mendonça, D. C., Rosa, H. V. D., Kumagai, P. S., et al. (2021). Orientational Ambiguity in Septin Coiled Coils and its Structural Basis. Journal of Molecular Biology 433, 166889. doi: 10.1016/j.jmb.2021.166889

Lobato-Márquez, D., Xu, J., Güler, G. Or., Ojiakor, A., Pilhofer, M., and Mostowy, S. (2021). Mechanistic insight into bacterial entrapment by septin cage reconstitution. Nat Commun 12, 4511. doi: 10.1038/s41467-021-24721-5

Longtine, M. S., DeMarini, D. J., Valencik, M. L., Al-Awar, O. S., Fares, H., De Virgilio, C., et al. (1996). The septins: roles in cytokinesis and other processes. Current Opinion in Cell Biology 8, 106–119. doi: 10.1016/S0955-0674(96)80054-8

Lupas, A., Van Dyke, M., and Stock, J. (1991). Predicting Coiled Coils from Protein Sequences. Science 252, 1162–1164. doi: 10.1126/science.252.5009.1162

Marques da Silva, R., Christe dos Reis Saladino, G., Antonio Leonardo, D., D’Muniz Pereira, H., Andréa Sculaccio, S., Paula Ulian Araujo, A. et al. (2023). A key piece of the puzzle: The central tetramer of the *Saccharomyces cerevisiae* septin protofilament and its implications for self-assembly. Journal of Structural Biology 215, 107983. doi: 10.1016/j.jsb.2023.107983

McMurray, M. A., and Thorner, J. (2008). “Biochemical Properties and Supramolecular Architecture of Septin Hetero-Oligomers and Septin Filaments,” in The Septins, (John Wiley & Sons, Ltd), 47–100. doi: 10.1002/9780470779705.ch3

Mendonça, D. C., Guimarães, S. L., Pereira, H. D., Pinto, A. A., de Farias, M. A., de Godoy, A. S., et al. (2021). An atomic model for the human septin hexamer by cryo-EM. Journal of Molecular Biology 433, 167096. doi: 10.1016/j.jmb.2021.167096

Miller, M. A., Pfeiffer, W., and Schwartz, T. (2010). Creating the CIPRES Science Gateway for inference of large phylogenetic trees., in 2010 Gateway Computing Environments Workshop (GCE), 1–8. doi: 10.1109/GCE.2010.5676129

Minh, B. Q., Schmidt, H. A., Chernomor, O., Schrempf, D., Woodhams, M. D., von Haeseler, A., et al. (2020). IQ-TREE 2: New Models and Efficient Methods for Phylogenetic Inference in the Genomic Era. Molecular Biology and Evolution 37, 1530–1534. doi: 10.1093/molbev/msaa015

Misof, B., and Misof, K. (2009). A Monte Carlo Approach Successfully Identifies Randomness in Multiple Sequence Alignments : A More Objective Means of Data Exclusion. Systematic Biology 58, 21–34. doi: 10.1093/sysbio/syp006

Moffat, L., and Jones, D. T. (2021). Increasing the accuracy of single sequence prediction methods using a deep semi-supervised learning framework. Bioinformatics 37, 3744– 3751. doi: 10.1093/bioinformatics/btab491

Momany, M., Zhao, J., Lindsey, R., and Westfall, P. J. (2001). Characterization of the Aspergillus nidulans septin (asp) gene family. Genetics 157, 969–977. doi: 10.1093/genetics/157.3.969

Nishihama, R., Onishi, M., and Pringle, J. R. (2011). New insights into the phylogenetic distribution and evolutionary origins of the septins. Biol Chem 392, 681–687. doi: 10.1515/BC.2011.086

Omrane, M., Camara, A. S., Taveneau, C., Benzoubir, N., Tubiana, T., Yu, J., et al. (2019). Septin 9 has Two Polybasic Domains Critical to Septin Filament Assembly and Golgi Integrity. iScience 13, 138–153. doi: 10.1016/j.isci.2019.02.015

Onishi, M., Koga, T., Hirata, A., Nakamura, T., Asakawa, H., Shimoda, C., et al. (2010). Role of septins in the orientation of forespore membrane extension during sporulation in fission yeast. Mol Cell Biol 30, 2057–2074. doi: 10.1128/MCB.01529-09

Onishi, M., and Pringle, J. R. (2016). The nonopisthokont septins: How many there are, how little we know about them, and how we might learn more. Methods Cell Biol 136, 1–19. doi: 10.1016/bs.mcb.2016.04.003

Pan, F., Malmberg, R. L., and Momany, M. (2007). Analysis of septins across kingdoms reveals orthology and new motifs. BMC Evol Biol 7, 103. doi: 10.1186/1471-2148-7-103

Papadopoulos, J. S., and Agarwala, R. (2007). COBALT: constraint-based alignment tool for multiple protein sequences. Bioinformatics 23, 1073–1079. doi: 10.1093/bioinformatics/btm076

Perry, J. A., Werner, M. E., Heck, B. W., Maddox, P. S., and Maddox, A. S. (2023). Septins throughout phylogeny are predicted to have a transmembrane domain, which in Caenorhabditis elegans is functionally important. 2023.11.20.567915. doi: 10.1101/2023.11.20.567915

Pinto, A. P. A., Pereira, H. M., Zeraik, A. E., Ciol, H., Ferreira, F. M., Brandão-Neto, J., et al. (2017). Filaments and fingers: Novel structural aspects of the single septin from Chlamydomonas reinhardtii. J Biol Chem 292, 10899–10911. doi: 10.1074/jbc.M116.762229

Schwefel, D., Arasu, B. S., Marino, S. F., Lamprecht, B., Köchert, K., Rosenbaum, E., et al. (2013). Structural Insights into the Mechanism of GTPase Activation in the GIMAP Family. Structure 21, 550–559. doi: 10.1016/j.str.2013.01.014

Shuman, B., and Momany, M. (2021). Septins From Protists to People. Front Cell Dev Biol 9, 824850. doi: 10.3389/fcell.2021.824850

Sirajuddin, M., Farkasovsky, M., Hauer, F., Kühlmann, D., Macara, I. G., Weyand, M., et al. (2007). Structural insight into filament formation by mammalian septins. Nature 449, 311–315. doi: 10.1038/nature06052

The UniProt Consortium (2023). UniProt: the Universal Protein Knowledgebase in 2023. Nucleic Acids Research 51, D523–D531. doi: 10.1093/nar/gkac1052

Trifinopoulos, J., Nguyen, L.-T., von Haeseler, A., and Minh, B. Q. (2016). W-IQ-TREE: a fast online phylogenetic tool for maximum likelihood analysis. Nucleic Acids Res 44, W232–W235. doi: 10.1093/nar/gkw256

van Hilten, N., Methorst, J., Verwei, N., and Risselada, H. J. (2023). Physics-based generative model of curvature sensing peptides; distinguishing sensors from binders. Science Advances 9, eade8839. doi: 10.1126/sciadv.ade8839

Versele, M., and Thorner, J. (2005). Some assembly required: yeast septins provide the instruction manual. Trends Cell Biol 15, 414–424. doi: 10.1016/j.tcb.2005.06.007

Weirich, C. S., Erzberger, J. P., and Barral, Y. (2008). The septin family of GTPases: architecture and dynamics. Nat Rev Mol Cell Biol 9, 478–489. doi: 10.1038/nrm2407

Wloga, D., Strzyzewska-Jówko, I., Gaertig, J., and Jerka-Dziadosz, M. (2008). Septins stabilize mitochondria in Tetrahymena thermophila. Eukaryot Cell 7, 1373–1386. doi: 10.1128/EC.00085-08

Woods, B. L., Cannon, K. S., Vogt, E. J. D., Crutchley, J. M., and Gladfelter, A. S. (2021). Interplay of septin amphipathic helices in sensing membrane-curvature and filament bundling. Mol Biol Cell 32, br5. doi: 10.1091/mbc.E20-05-0303

Yamazaki, T., Owari, S., Ota, S., Sumiya, N., Yamamoto, M., Watanabe, K., et al. (2013). Localization and evolution of septins in algae. The Plant Journal 74, 605–614. doi: 10.1111/tpj.12147

Zhang, J., Kong, C., Xie, H., McPherson, P. S., Grinstein, S., and Trimble, W. S. (1999). Phosphatidylinositol polyphosphate binding to the mammalian septin H5 is modulated by GTP. Current Biology 9, 1458–1467. doi: 10.1016/S0960-9822(00)80115-3

