## Supplementary figures and images for "The Evolutionary Origins and Ancestral Features of Septins"

### AncChlorophyte_450_1250_60852_coverage.png

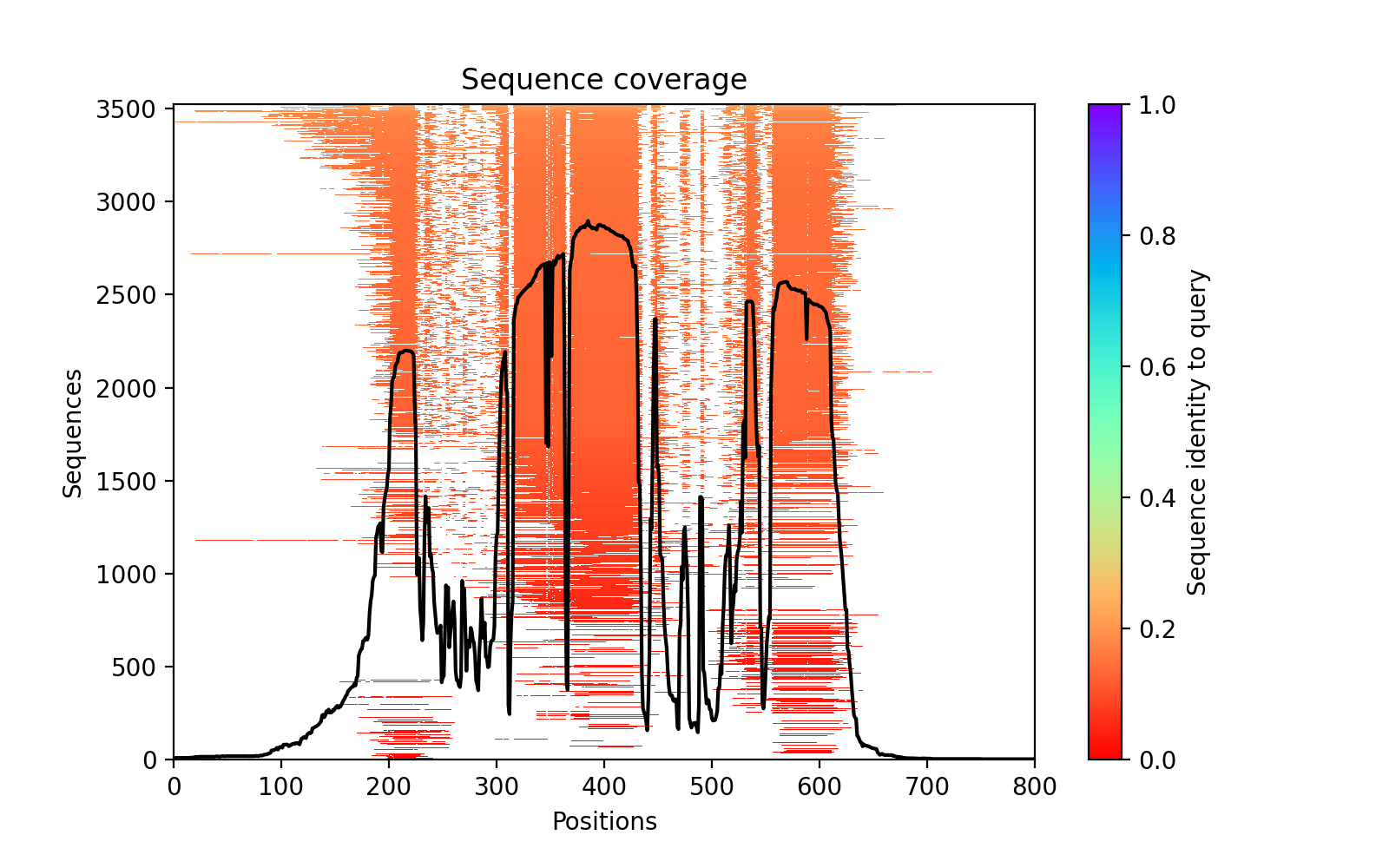

### AncChlorophyte_450_1250_60852_PAE.png

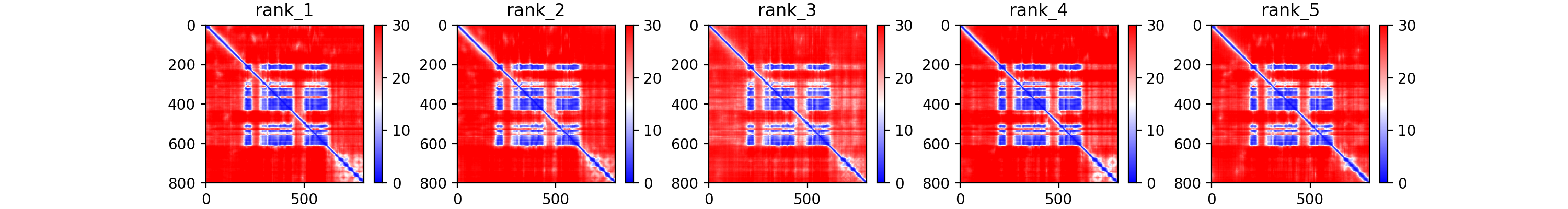

### AncChlorophyte_450_1250_60852_plddt.png

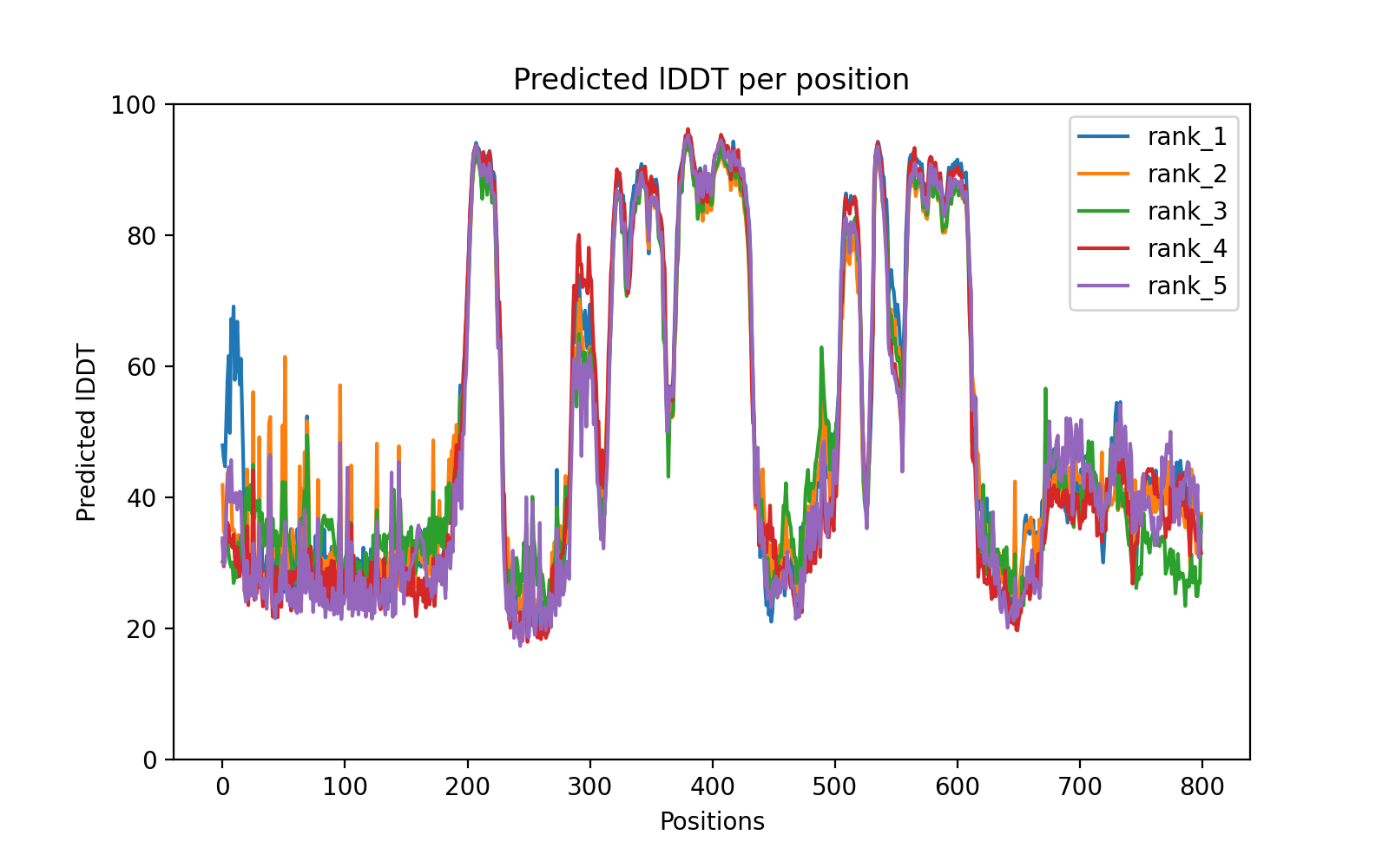

### AncChlorophyte_Volvo_2173e_coverage.png

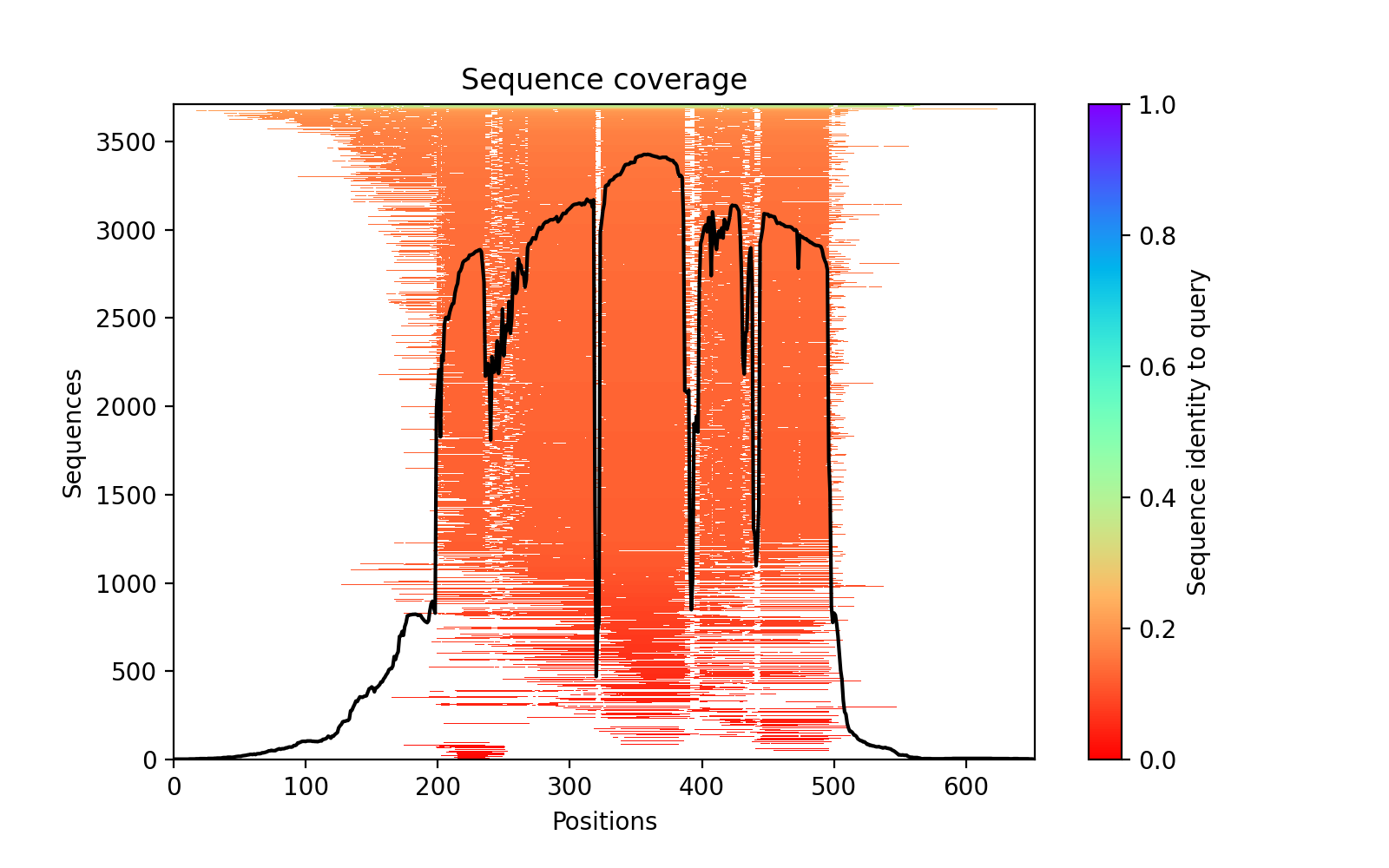

### AncChlorophyte_Volvo_2173e_PAE.png

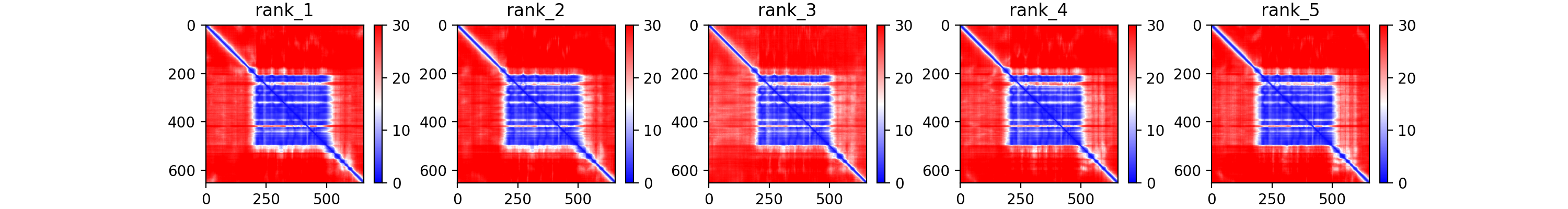

### AncChlorophyte_Volvo_2173e_plddt.png

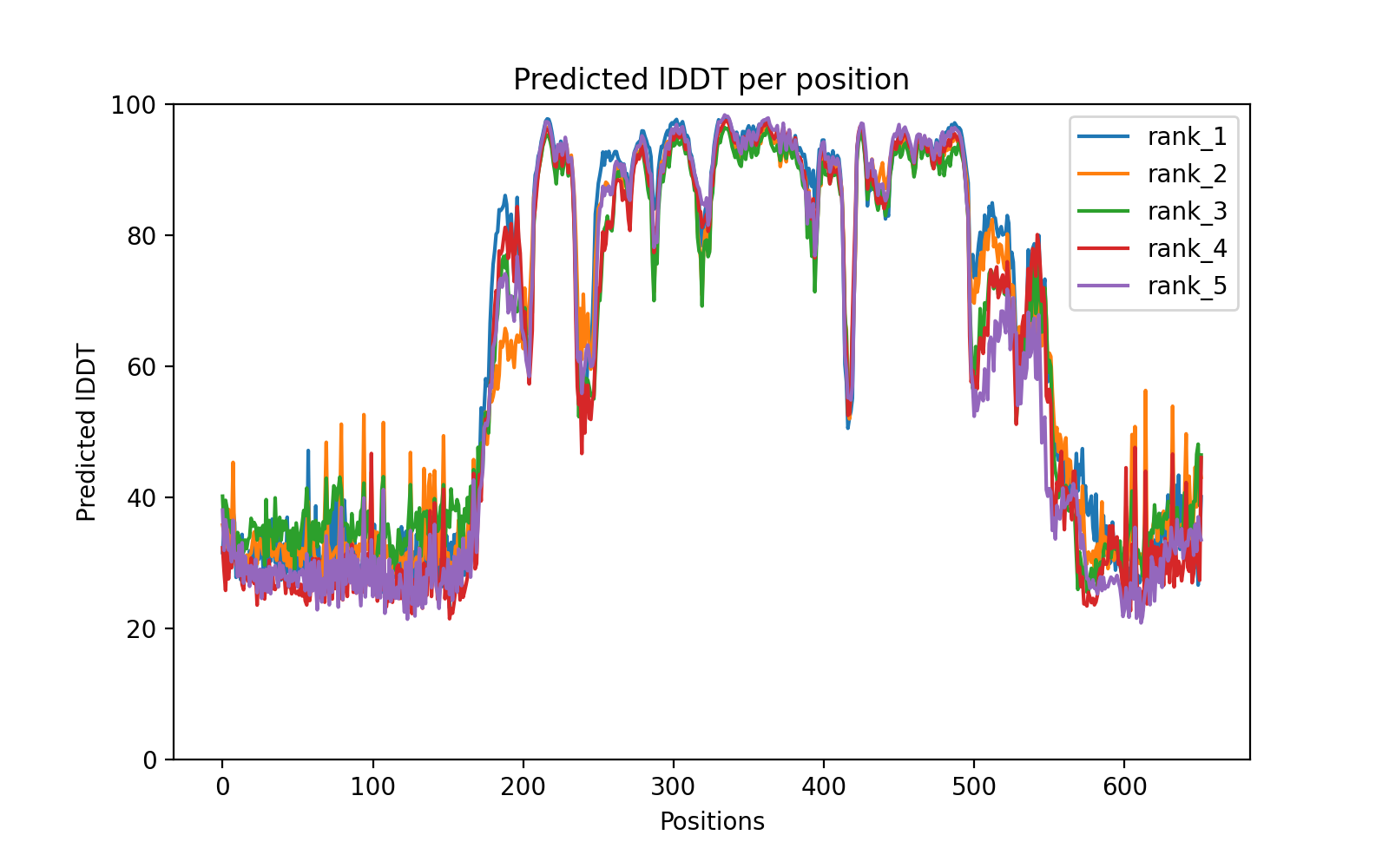

### AncCiliates_200_978_9bb52_coverage.png

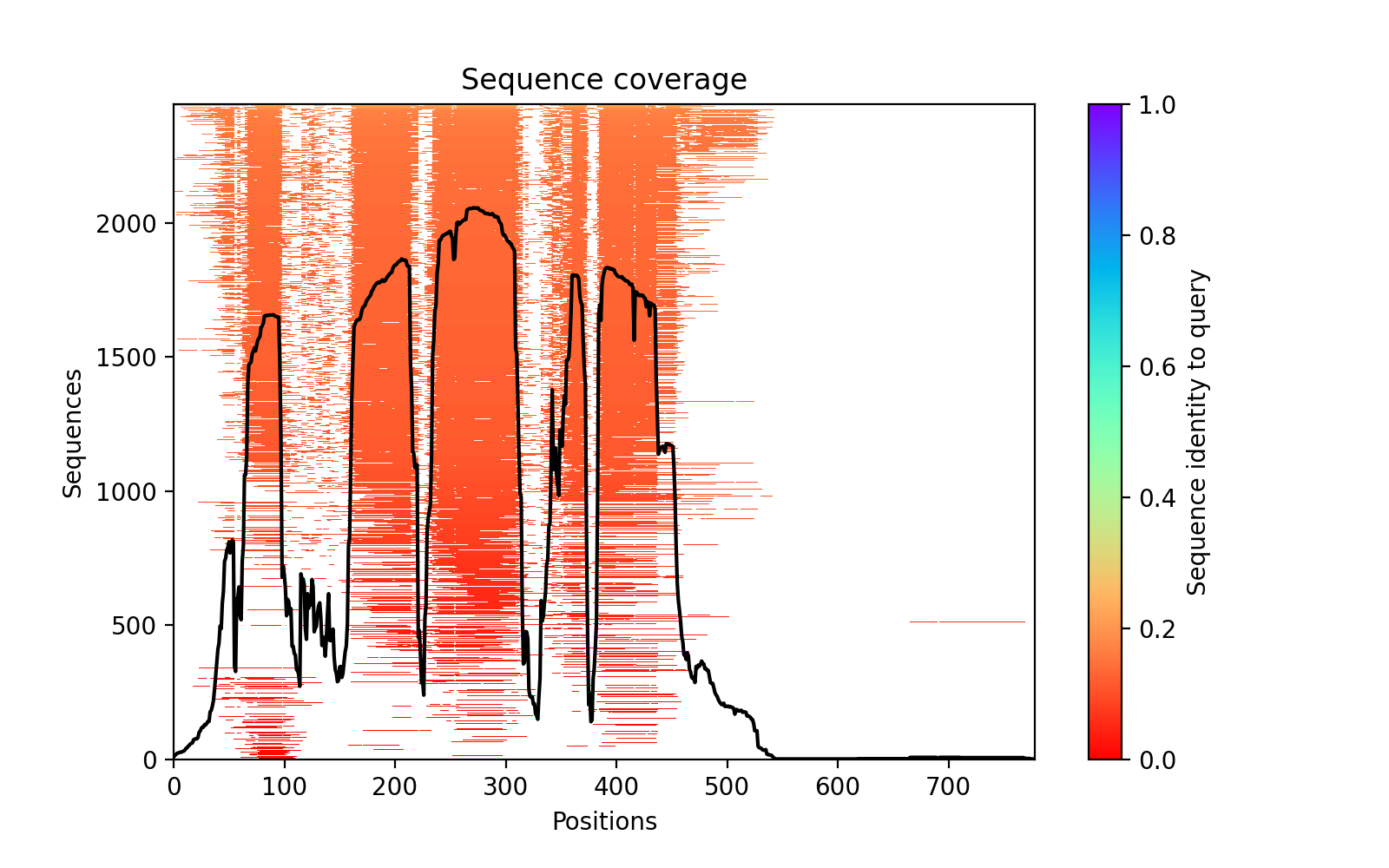

### AncCiliates_200_978_9bb52_PAE.png

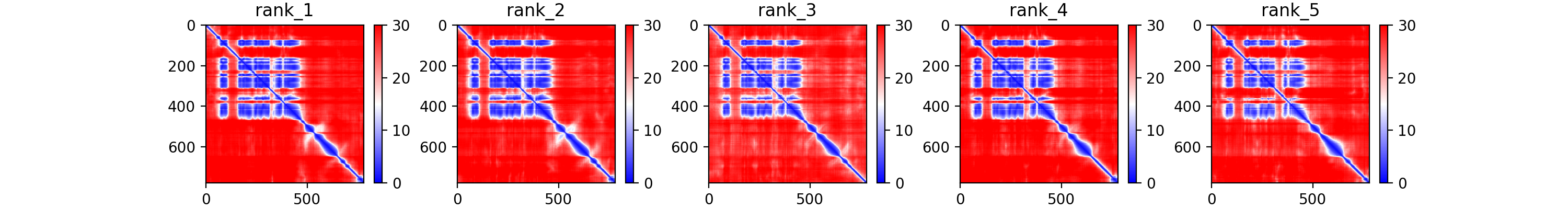

### AncCiliates_200_978_9bb52_plddt.png

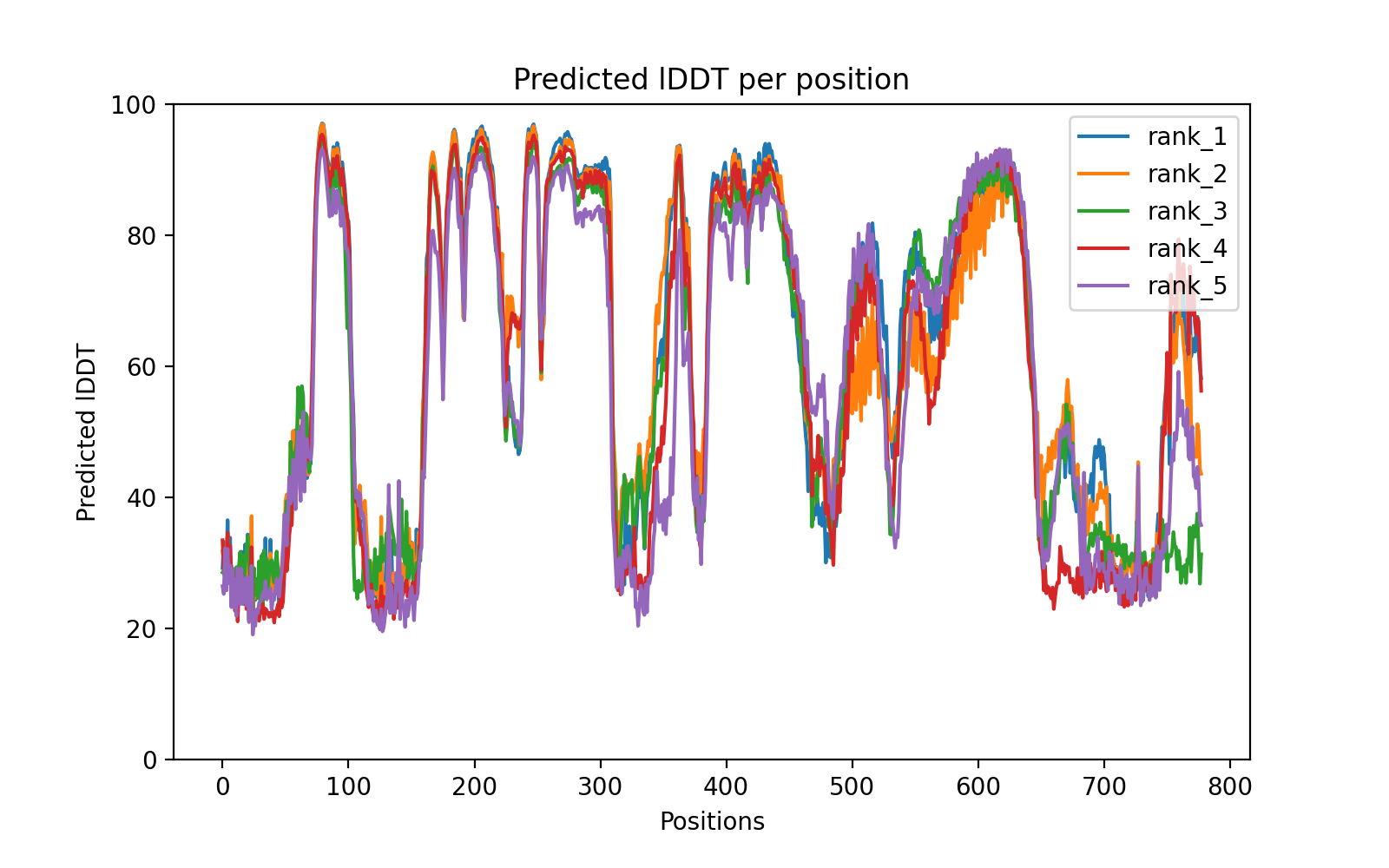

### AncGeneralChlorophyte_450_1150_73d48_coverage.png

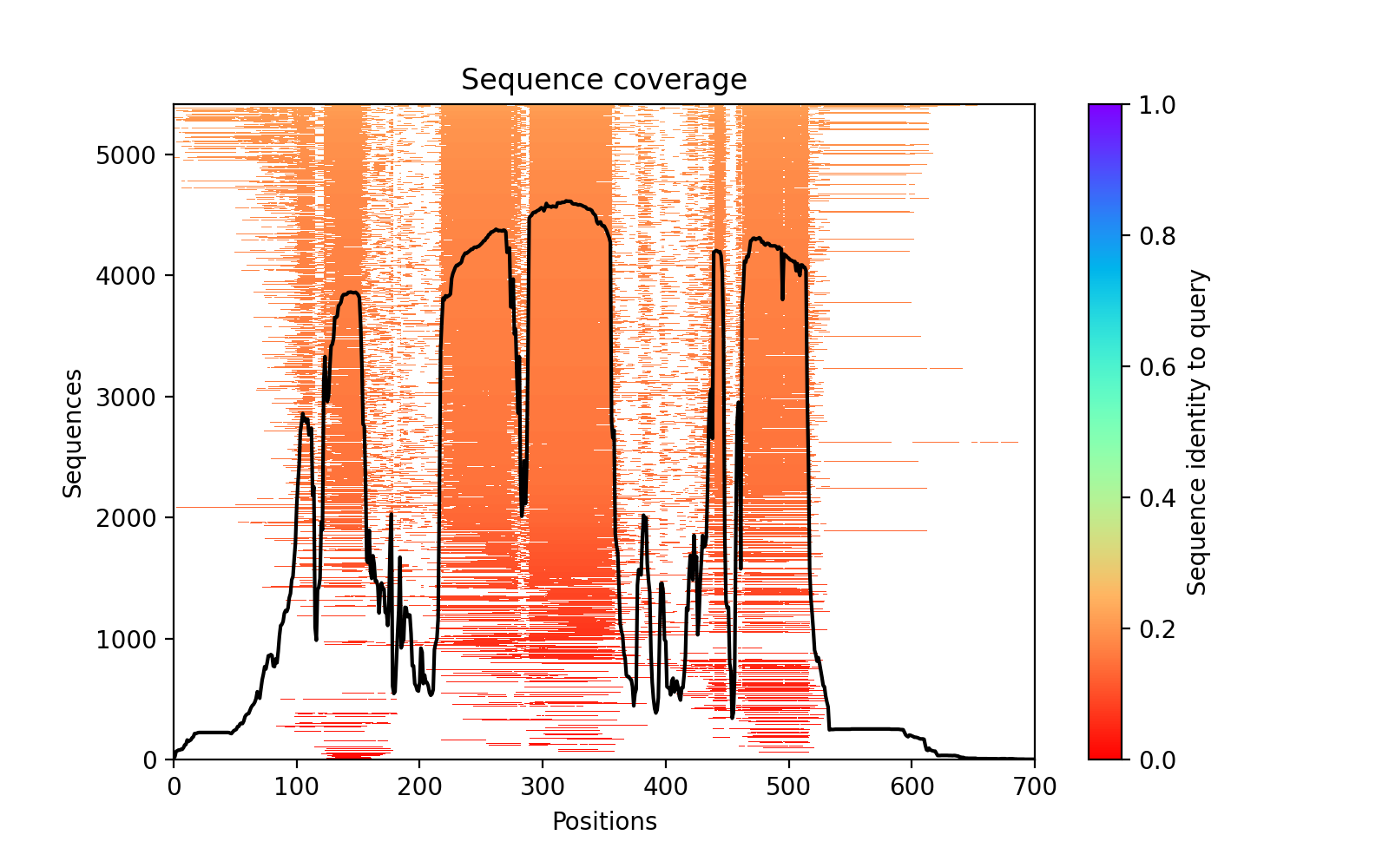

### AncGeneralChlorophyte_450_1150_73d48_PAE.png

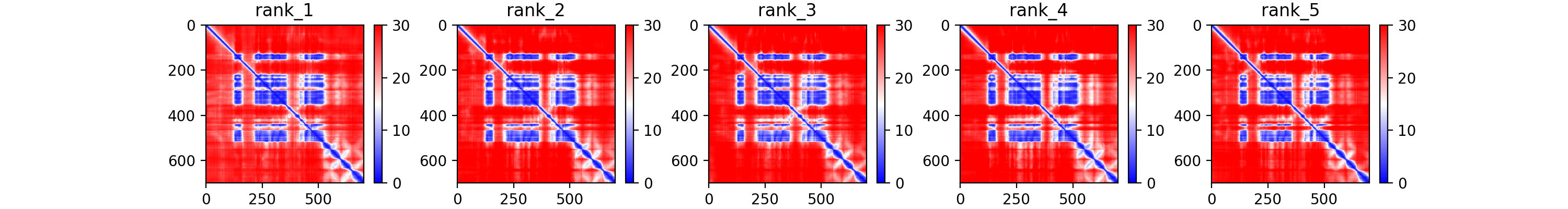

### AncGeneralChlorophyte_450_1150_73d48_plddt.png

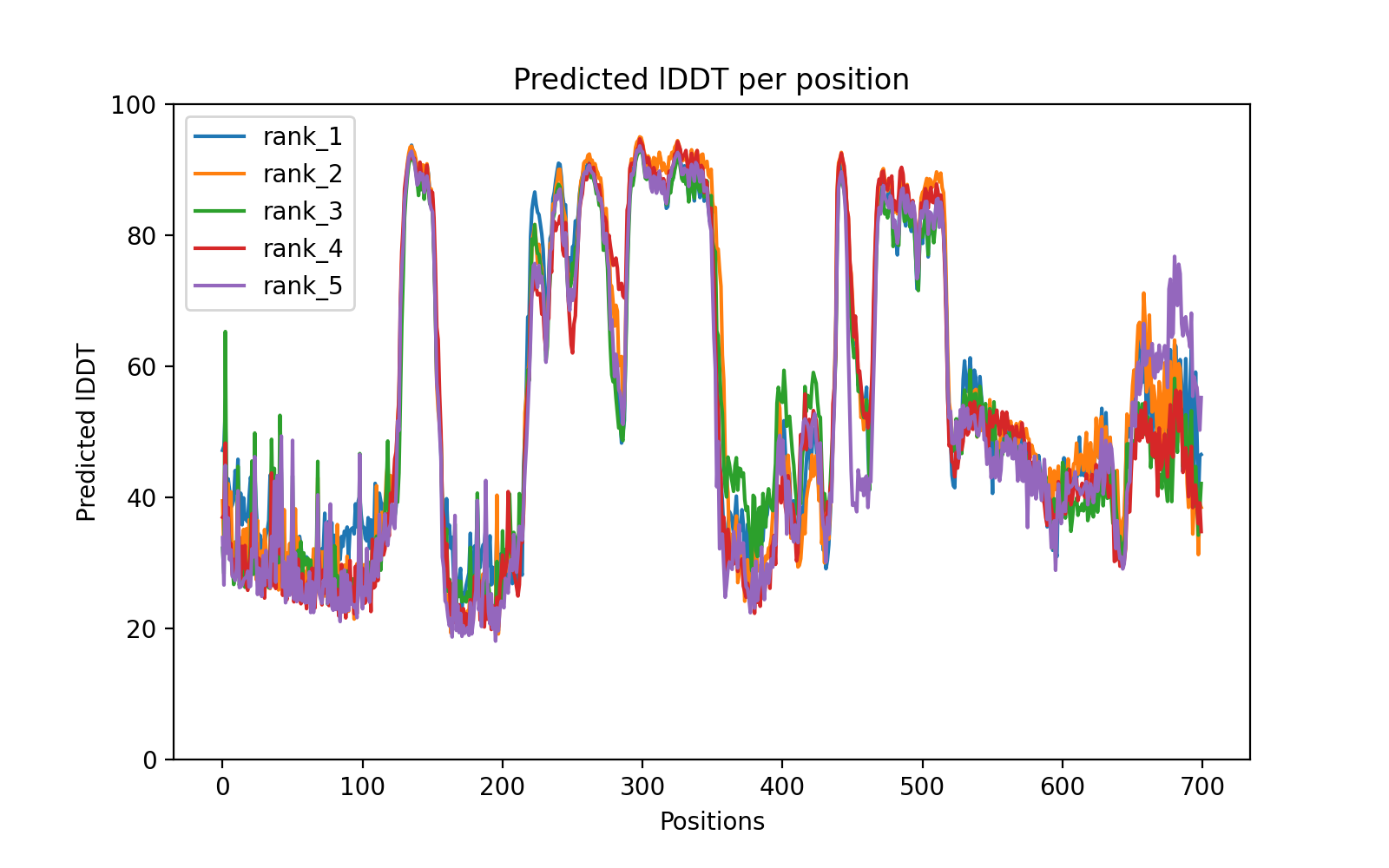

### AncGroup1_1a383_coverage.png

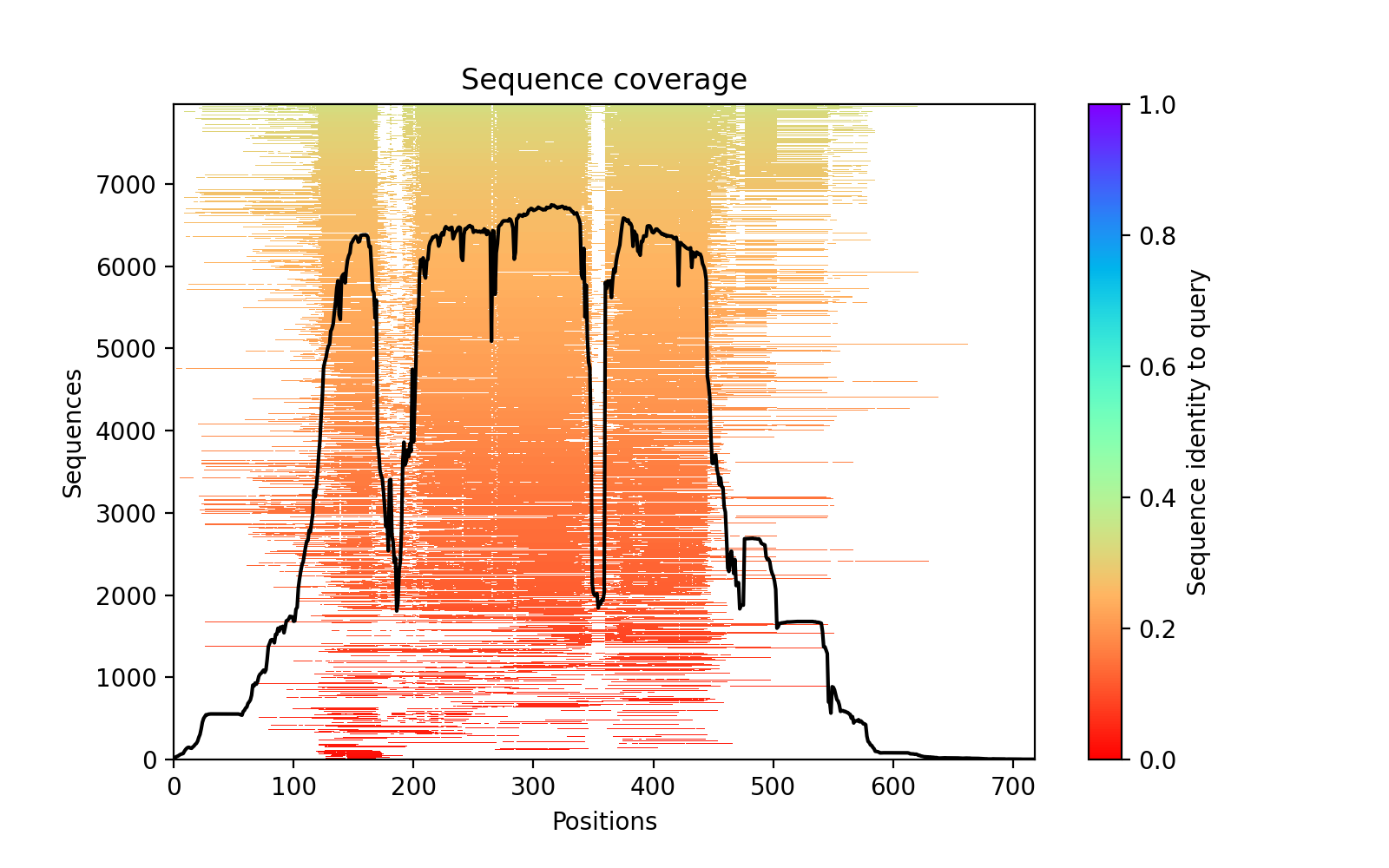

### AncGroup1_1a383_PAE.png

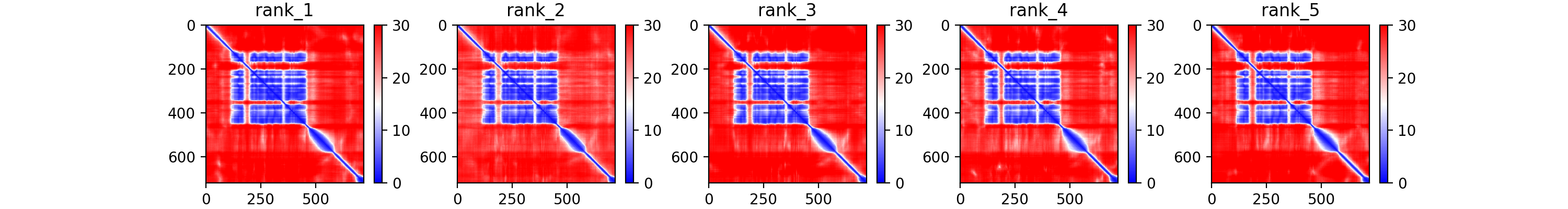

### AncGroup1_1a383_plddt.png

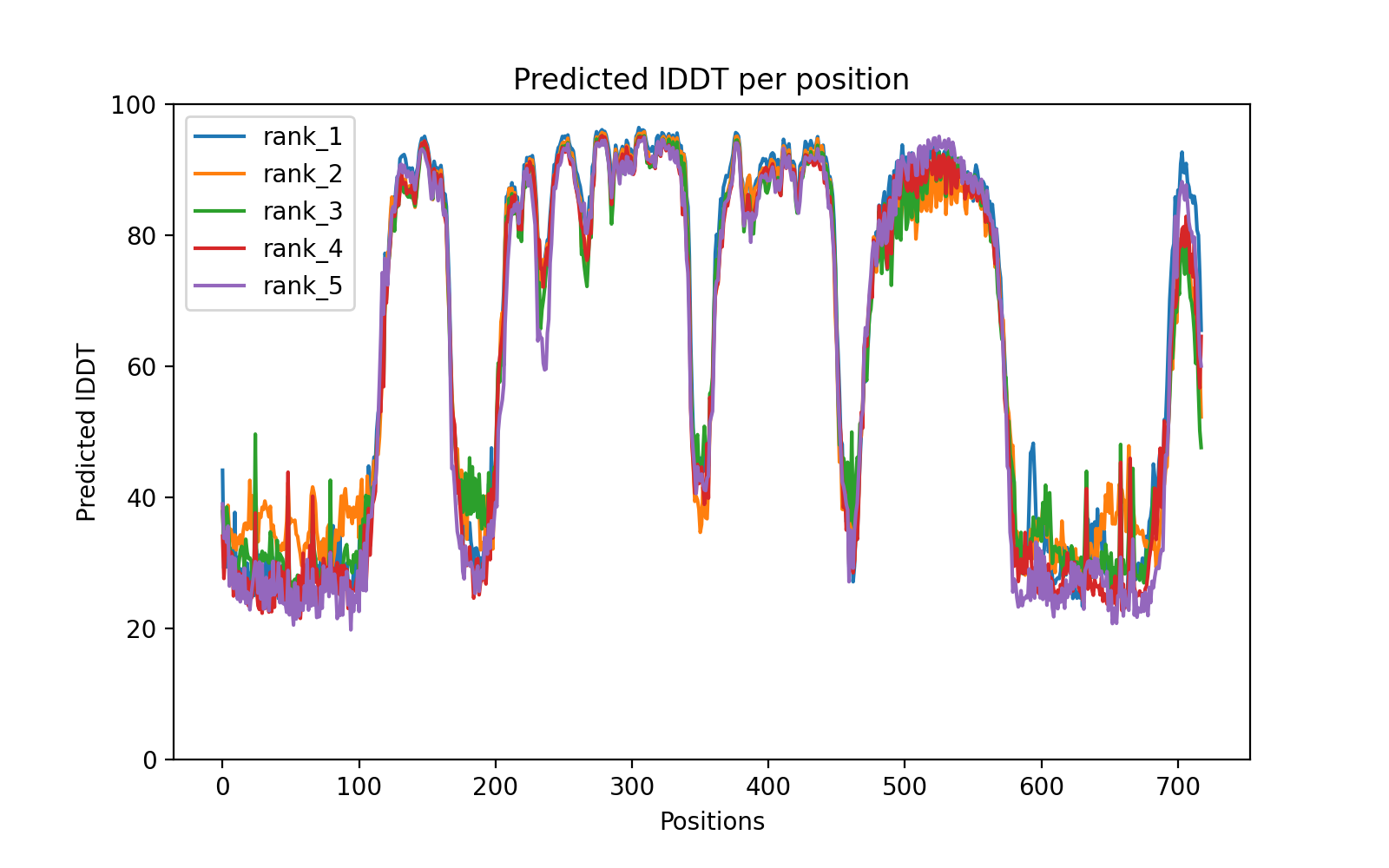

### AncGroup2_3fced_coverage.png

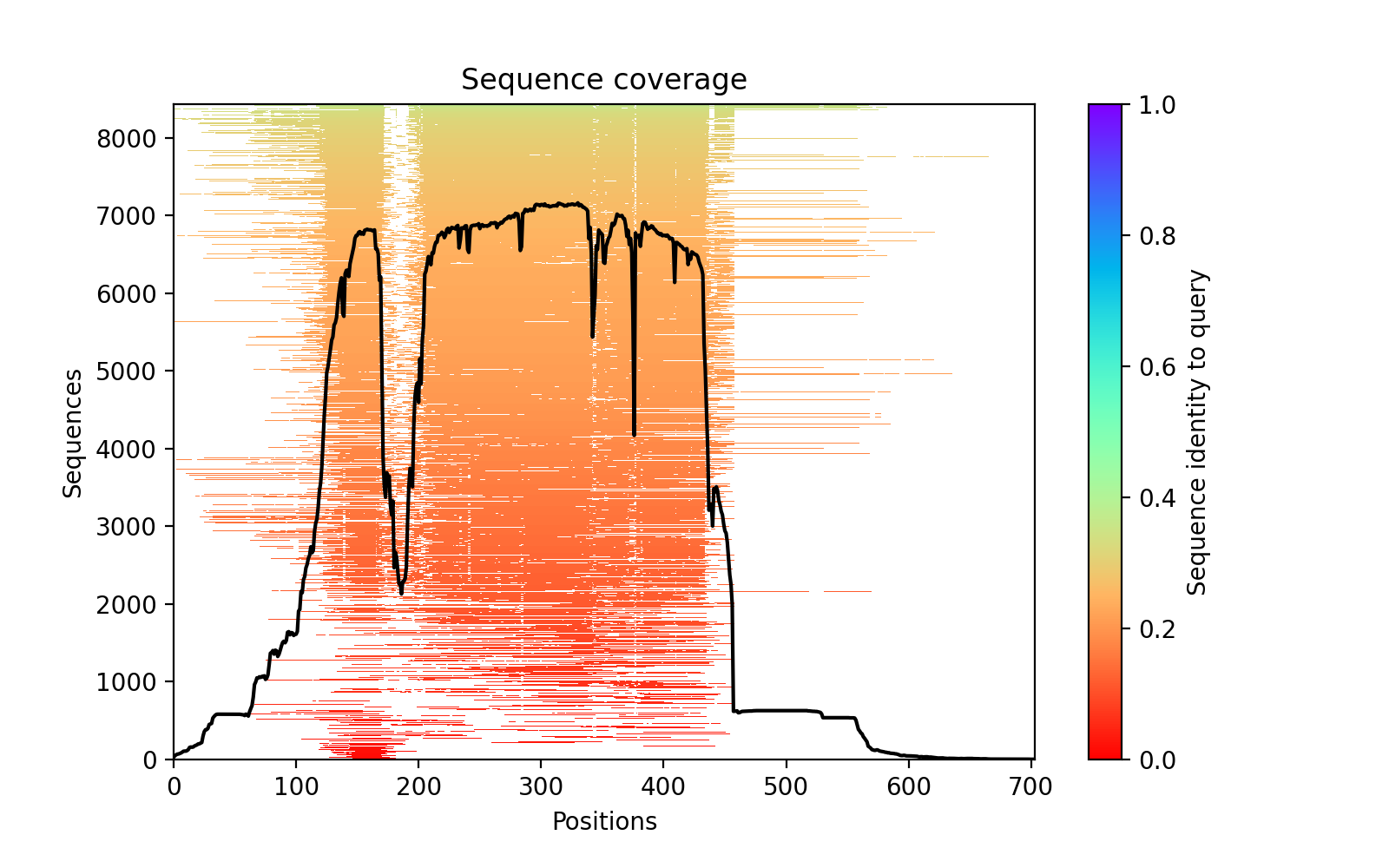

### AncGroup2_3fced_PAE.png

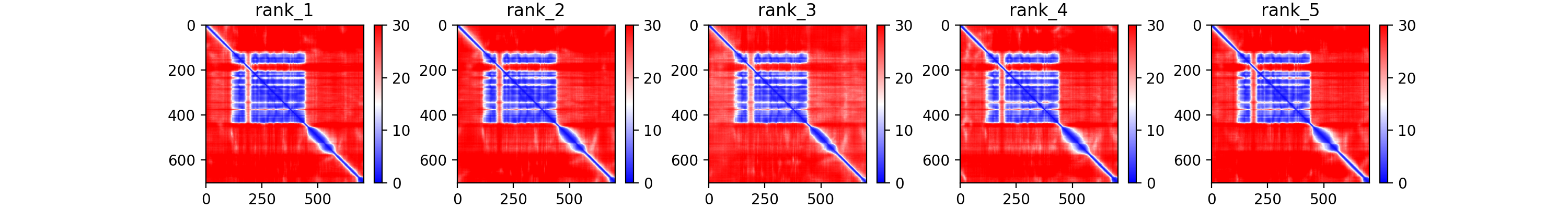

### AncGroup2_3fced_plddt.png

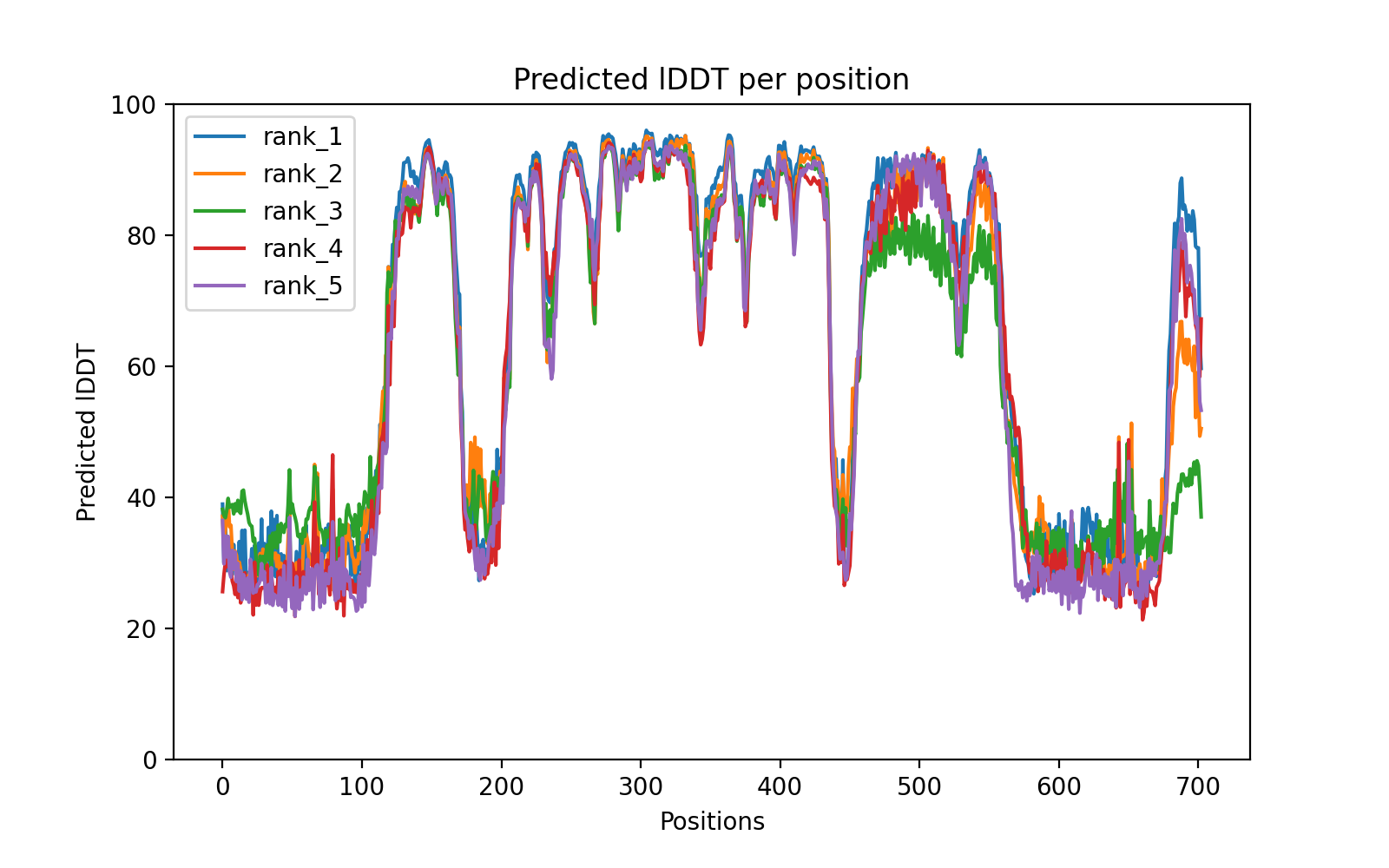

### AncGroup3_2f3f8_coverage.png

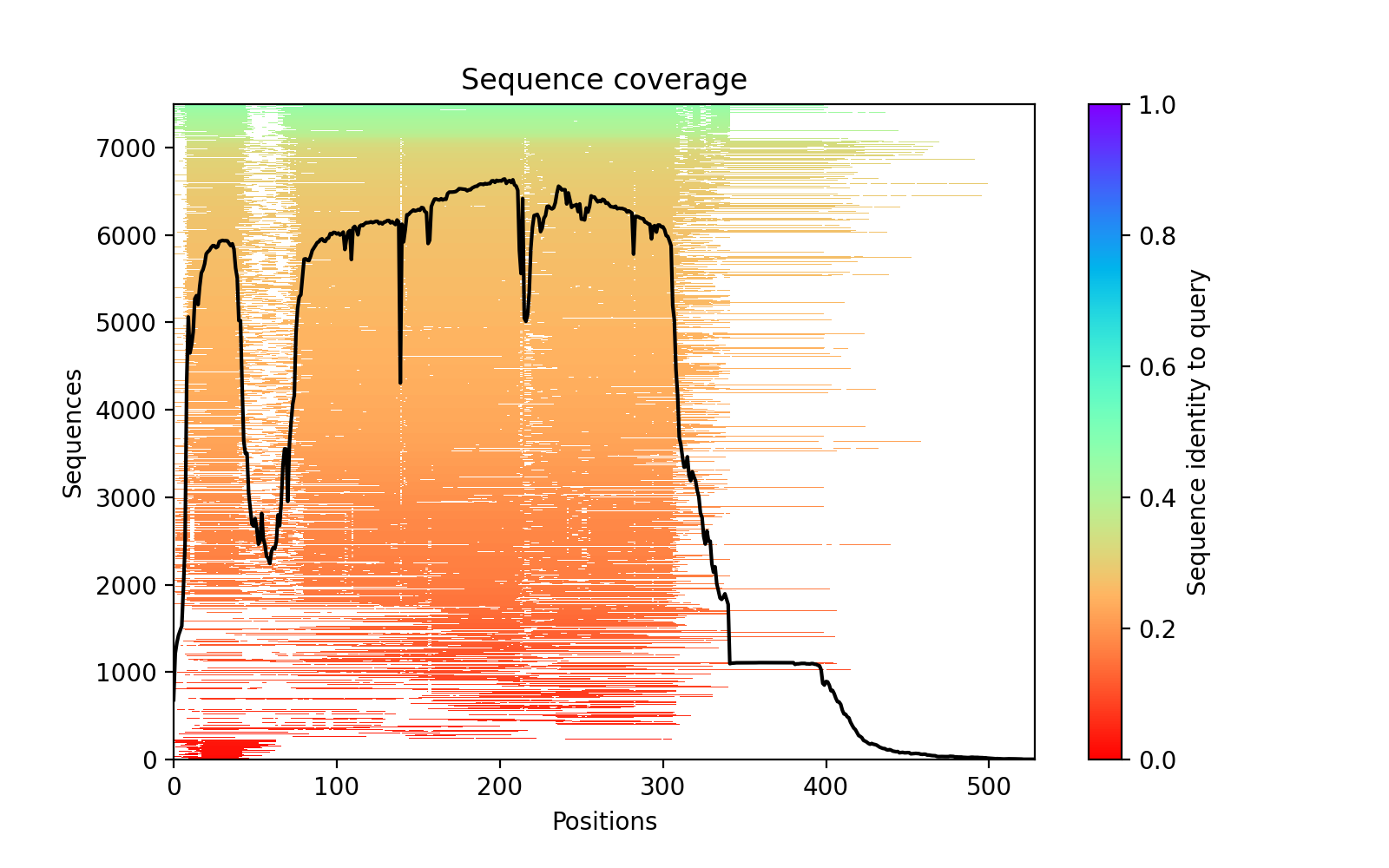

### AncGroup3_2f3f8_PAE.png

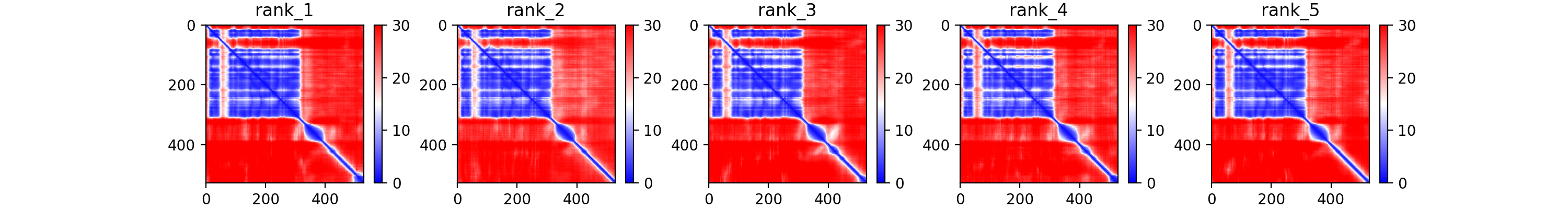

### AncGroup3_2f3f8_plddt.png

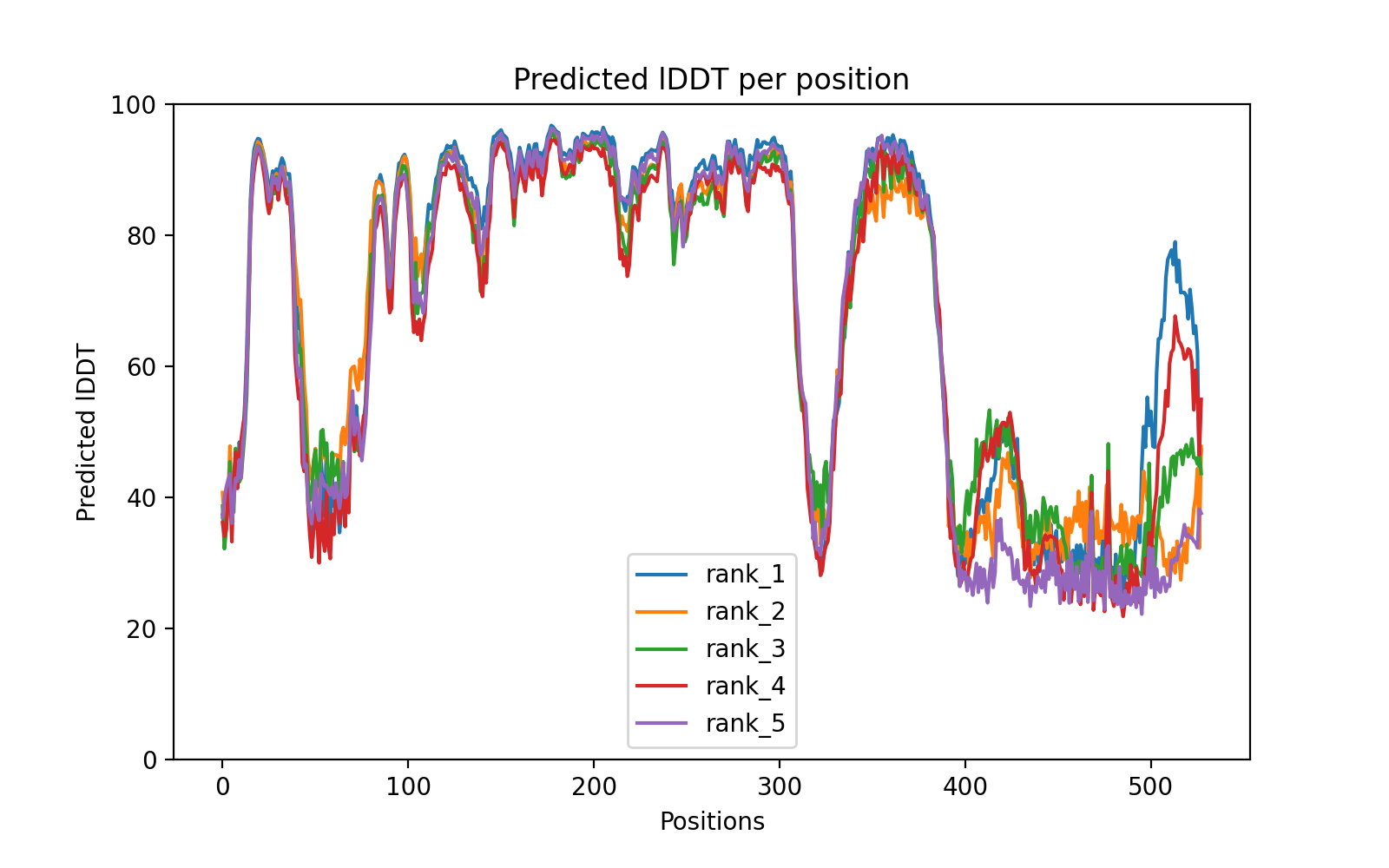

### AncGroup4_1_750_91333_coverage.png

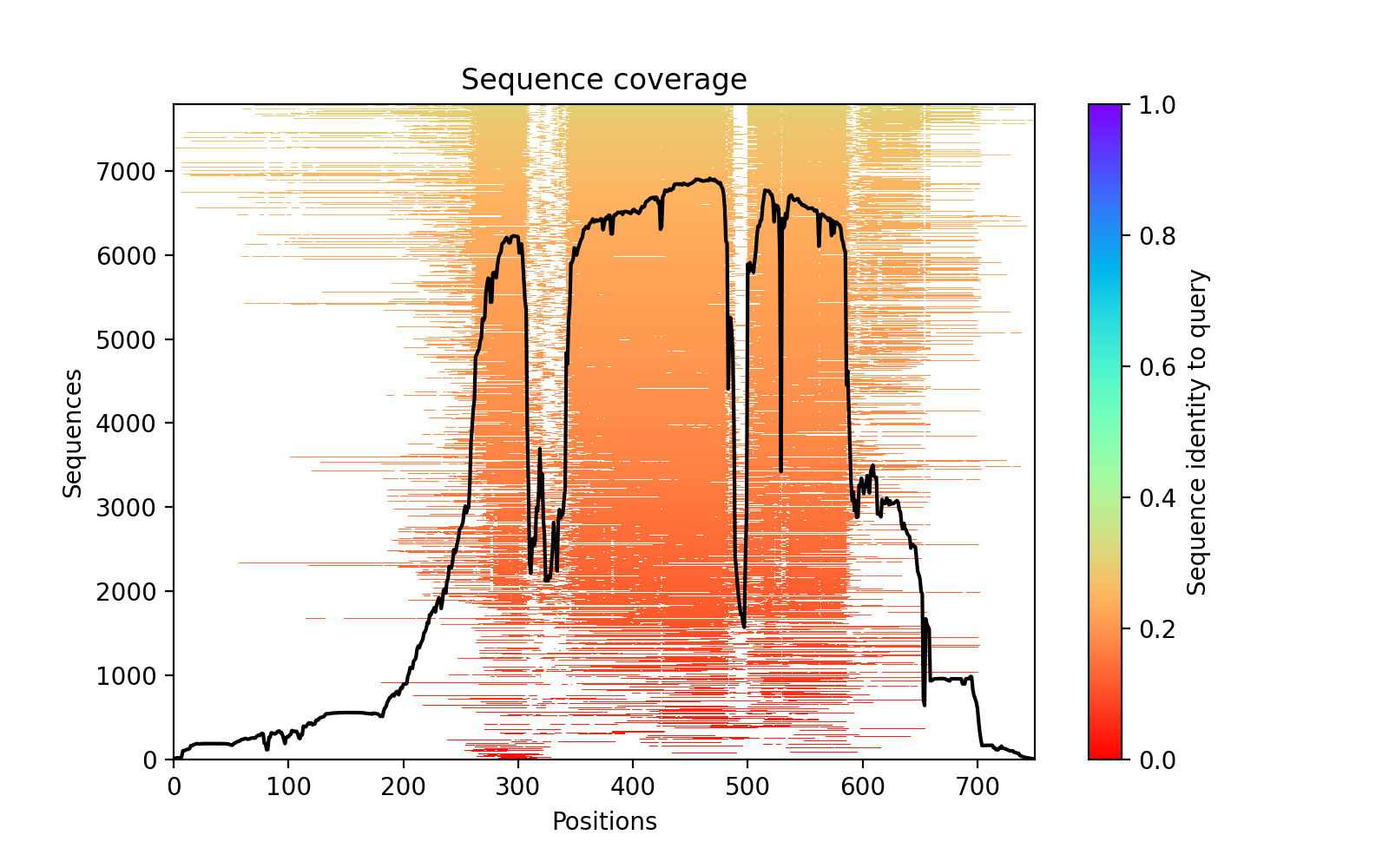

### AncGroup4_1_750_91333_PAE.png

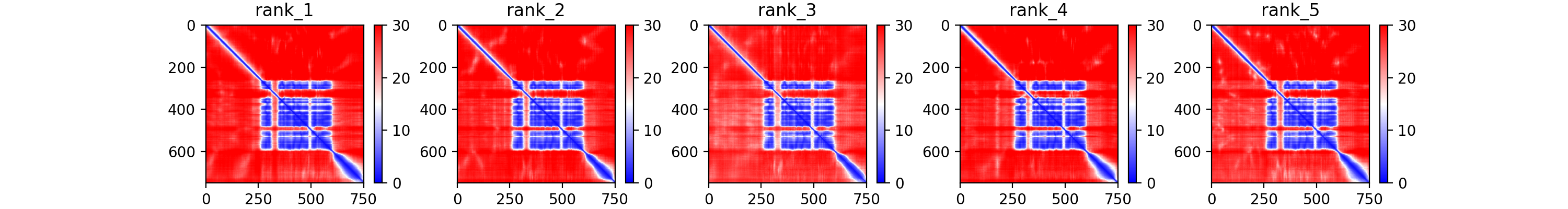

### AncGroup4_1_750_91333_plddt.png

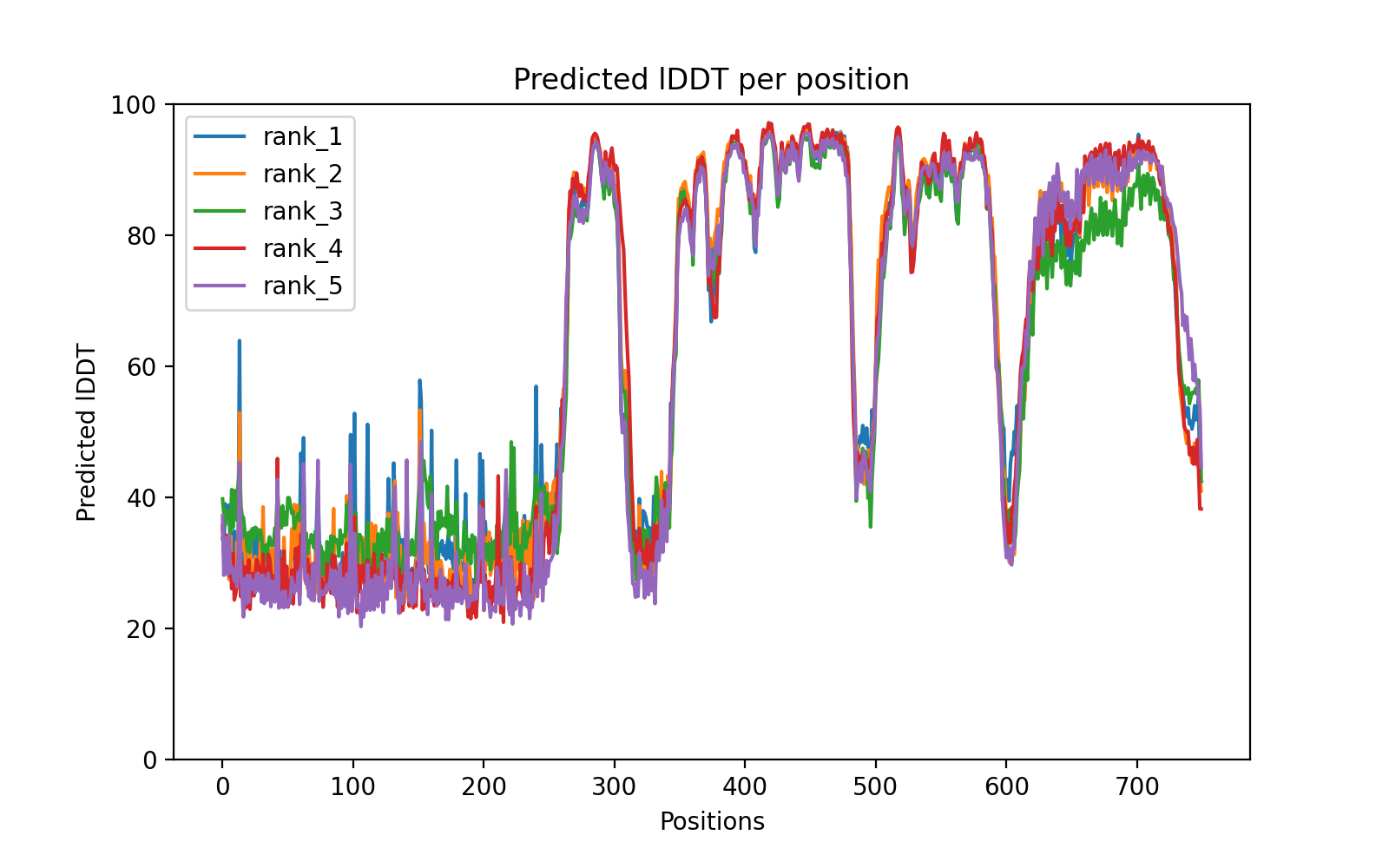

### AncGroup5_200_954_98a30_coverage.png

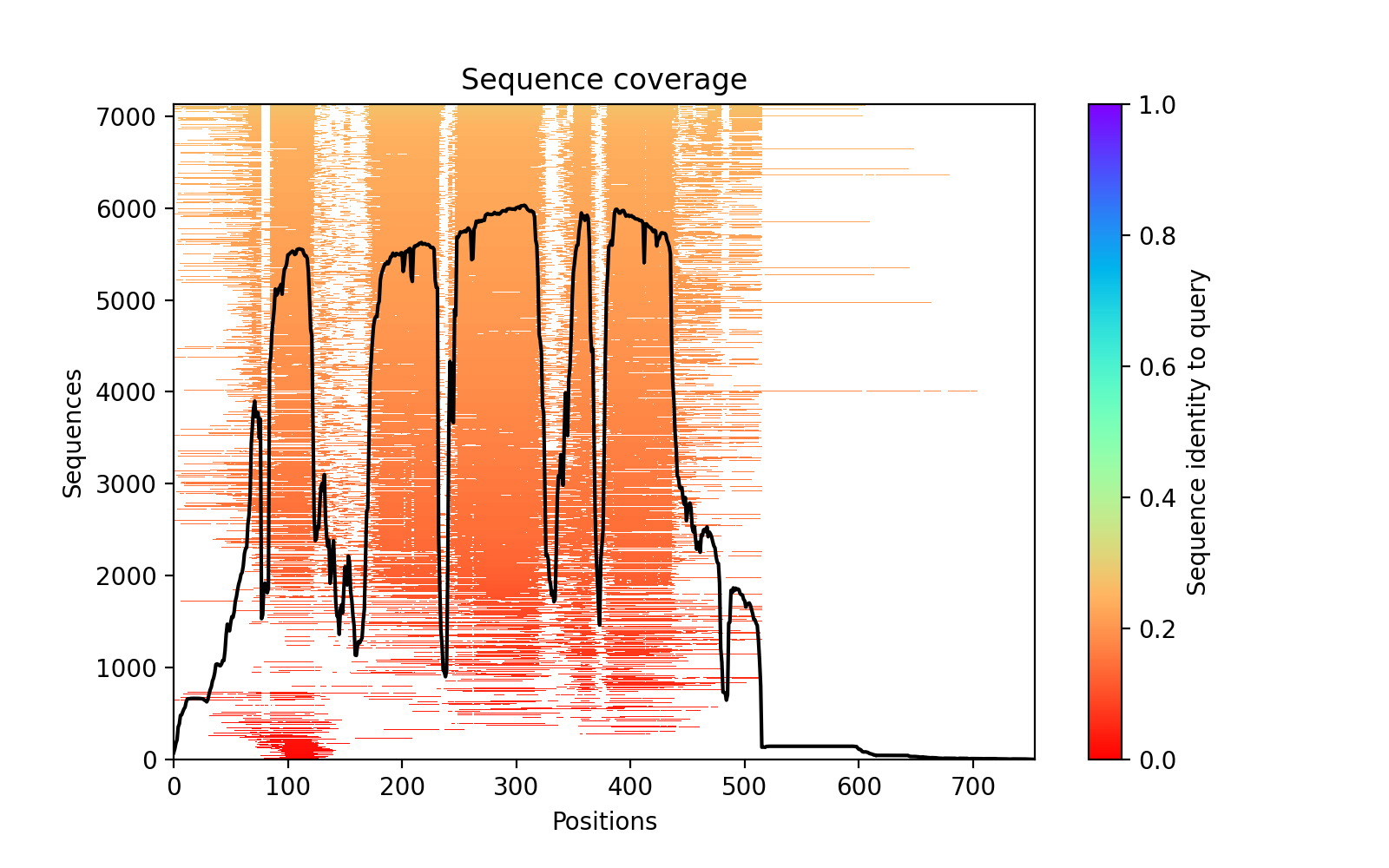

### AncGroup5_200_954_98a30_PAE.png

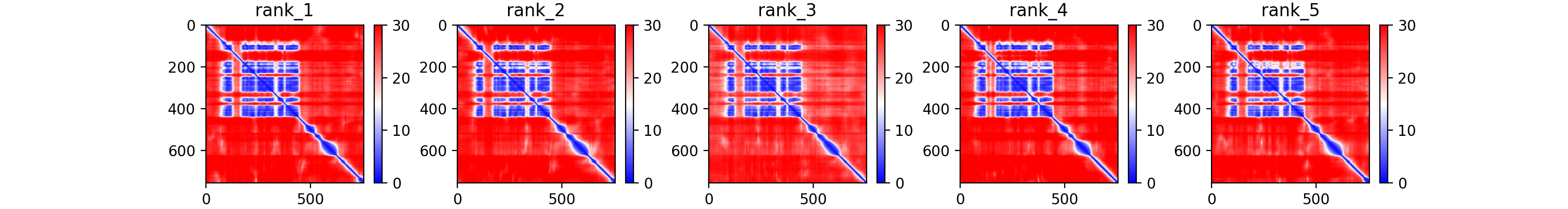

### AncGroup5_200_954_98a30_plddt.png

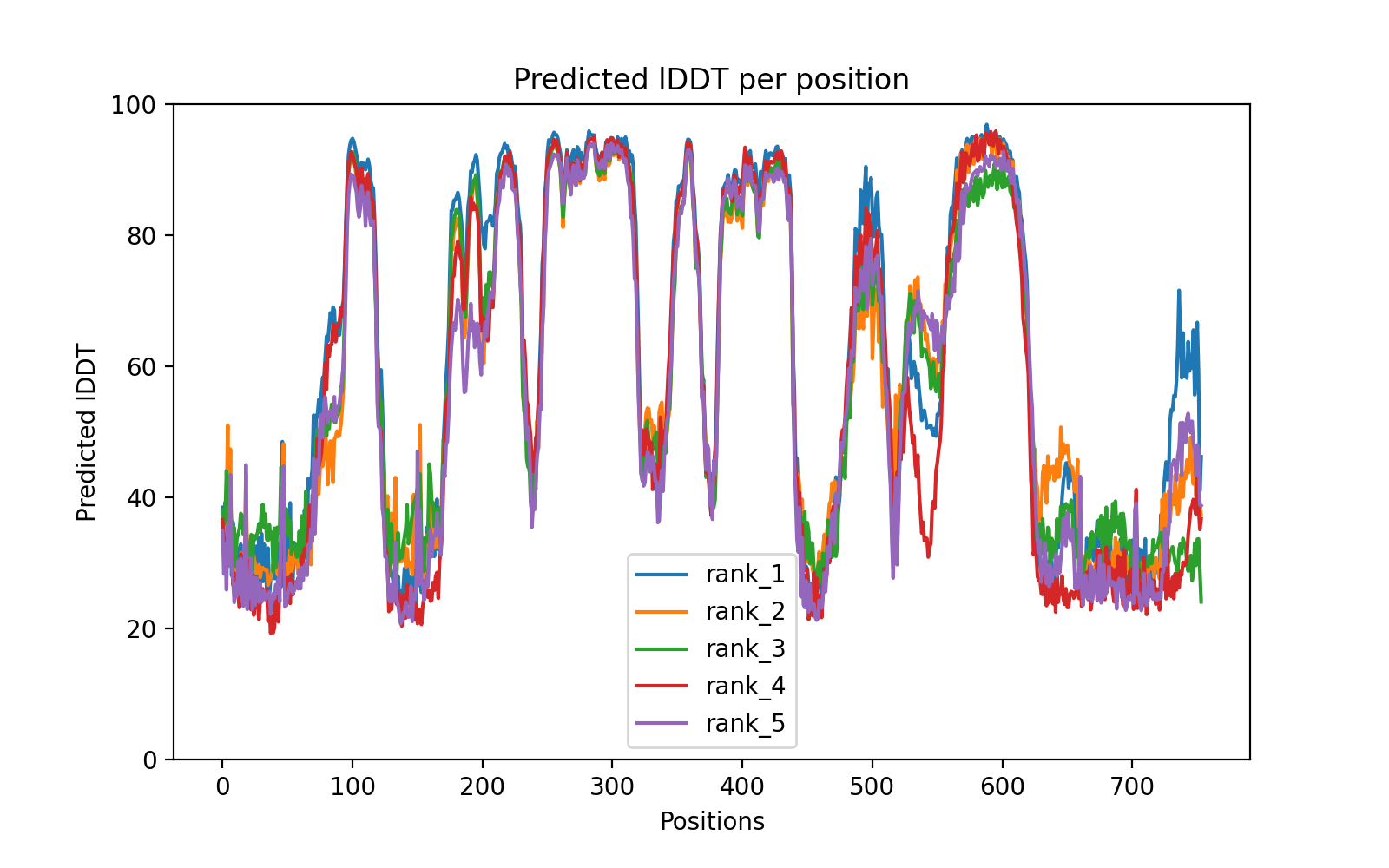

### AncOpis_200_900_b2ee0_coverage.png

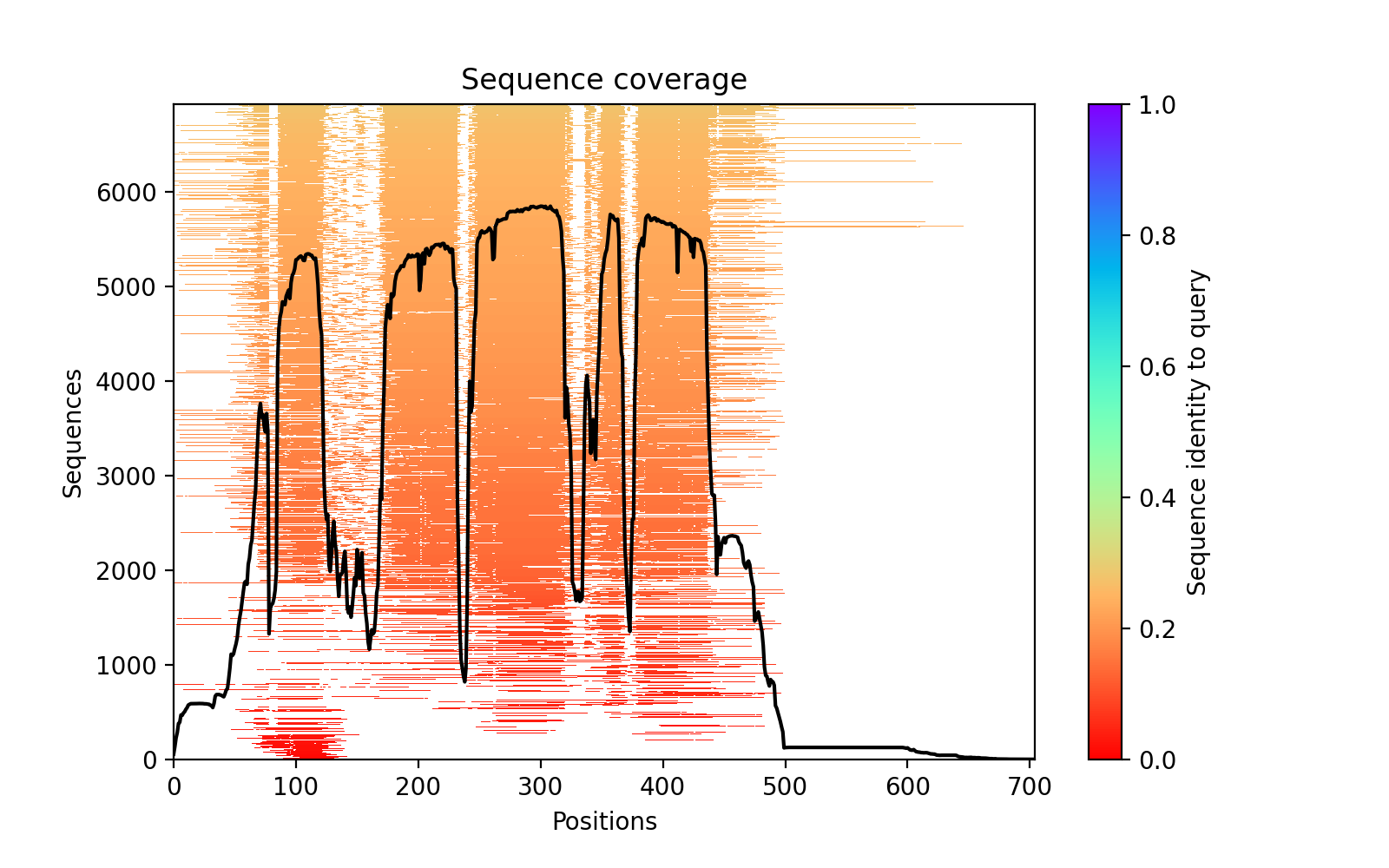

### AncOpis_200_900_b2ee0_PAE.png

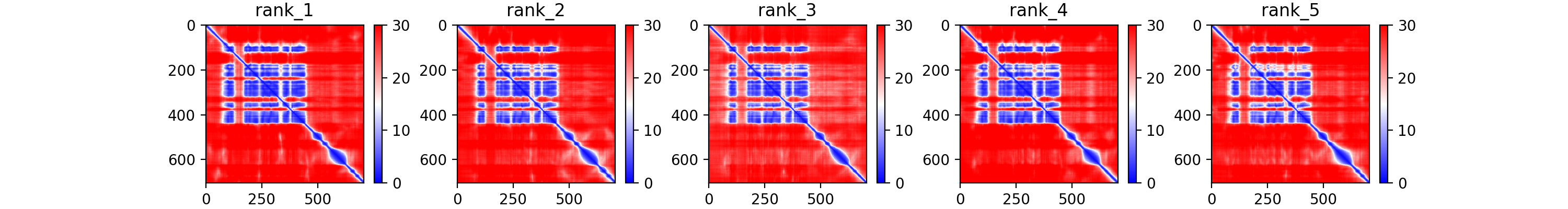

### AncOpis_200_900_b2ee0_plddt.png

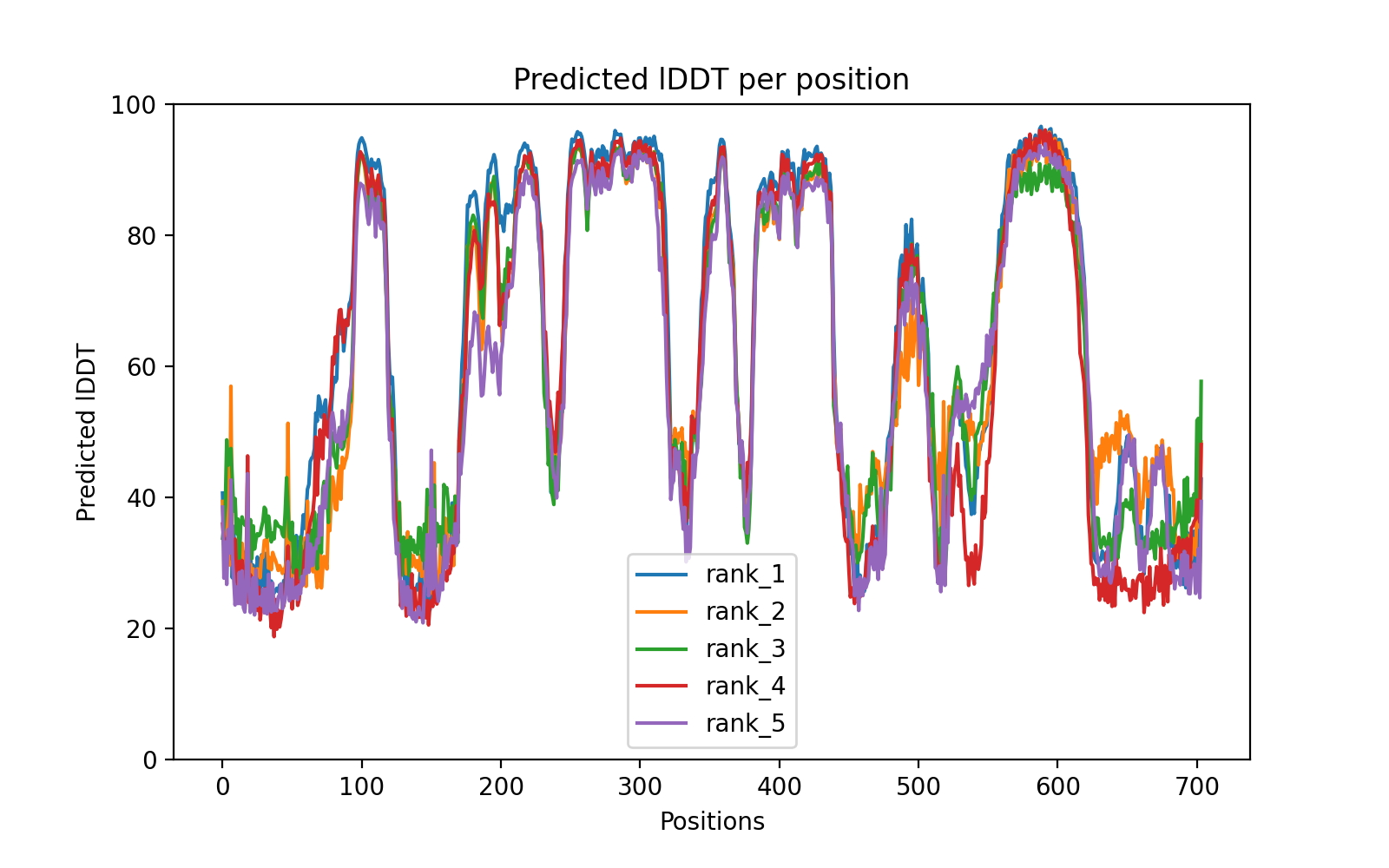

### AncProt_450___47750_coverage.png

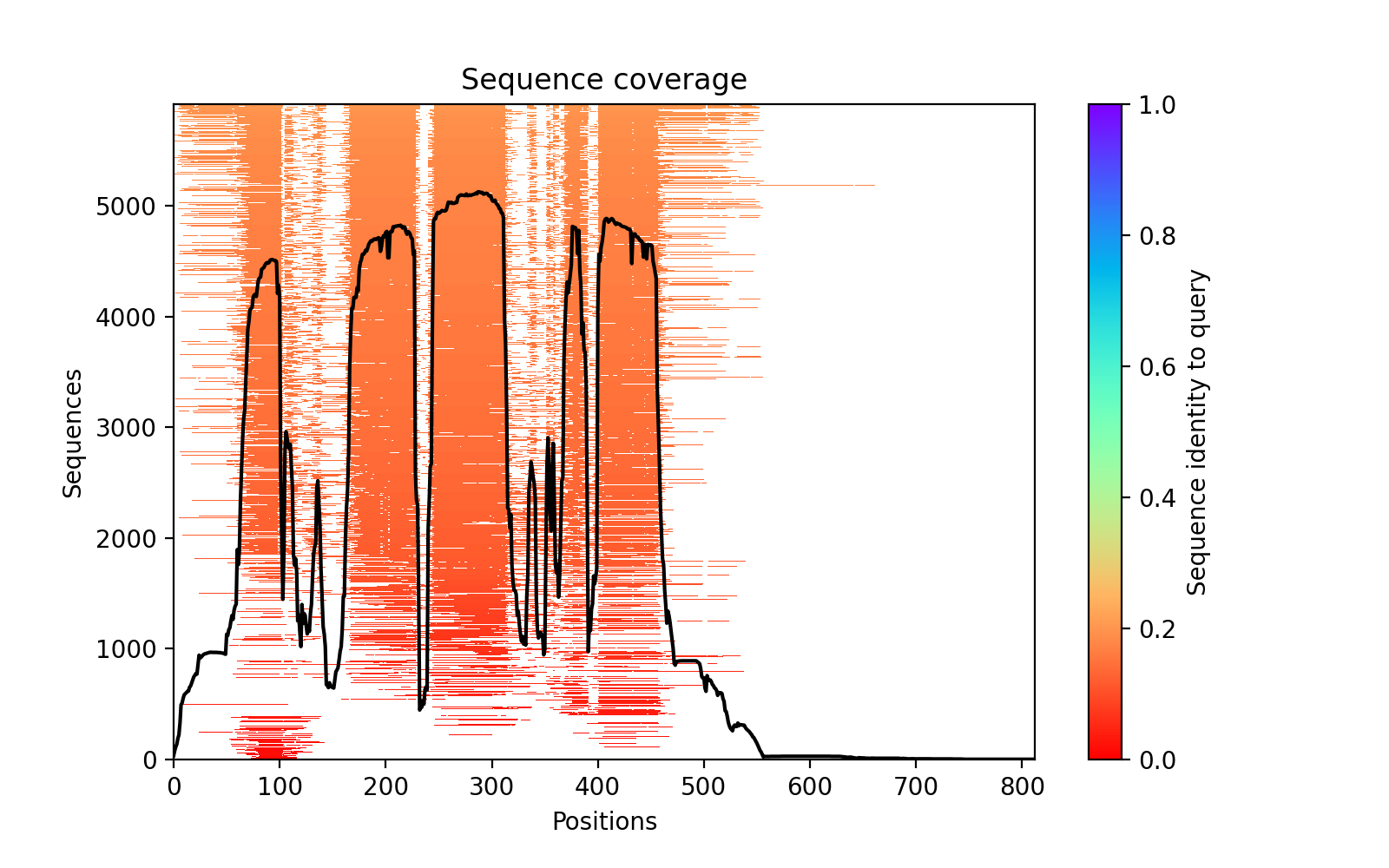

### AncProt_450___47750_PAE.png

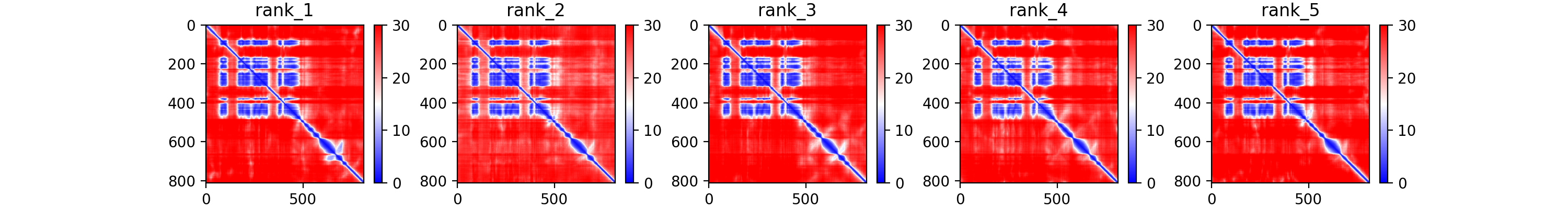

### AncProt_450___47750_plddt.png

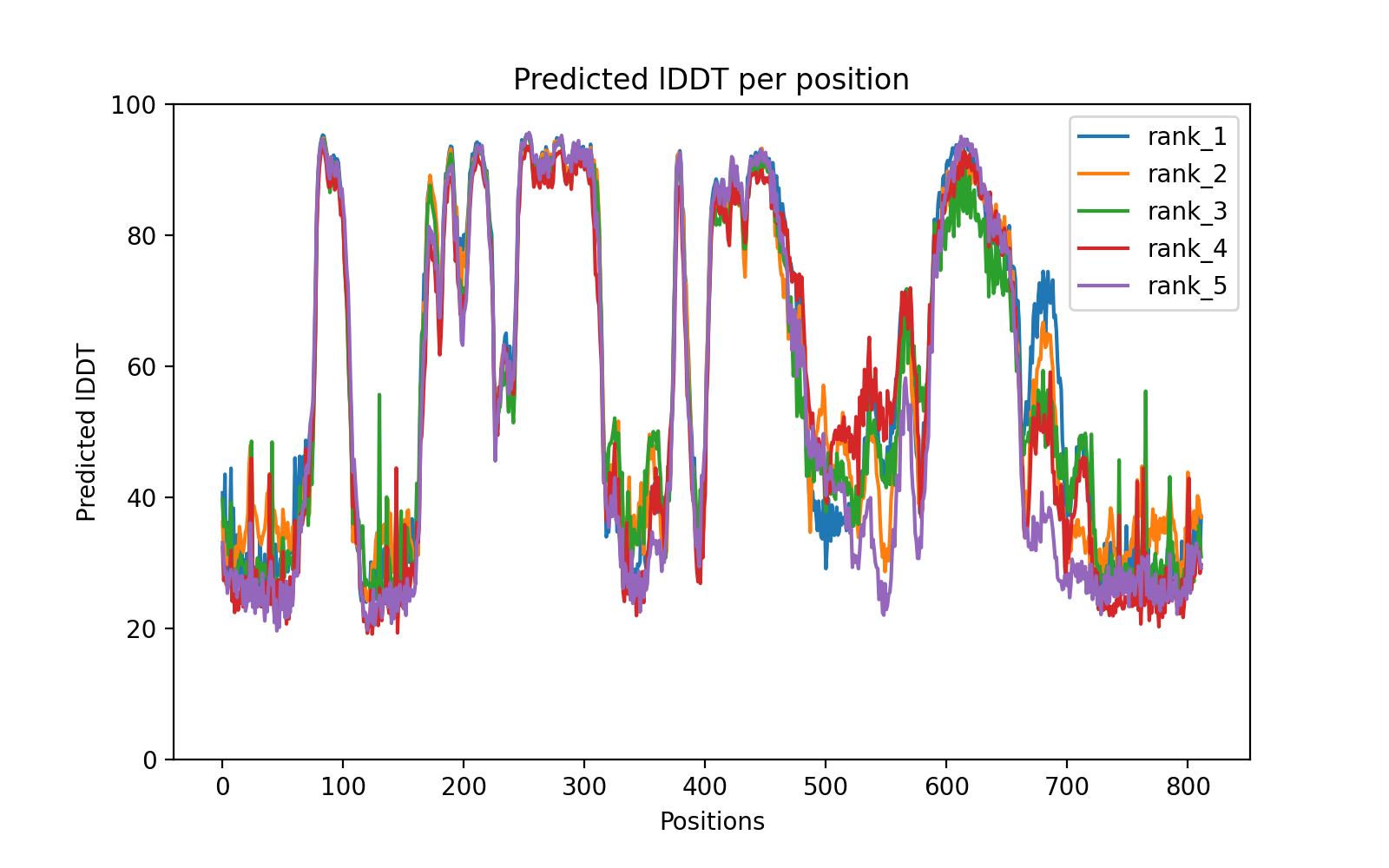

### AncSeptin_N4_385_1281_9397d_coverage.png

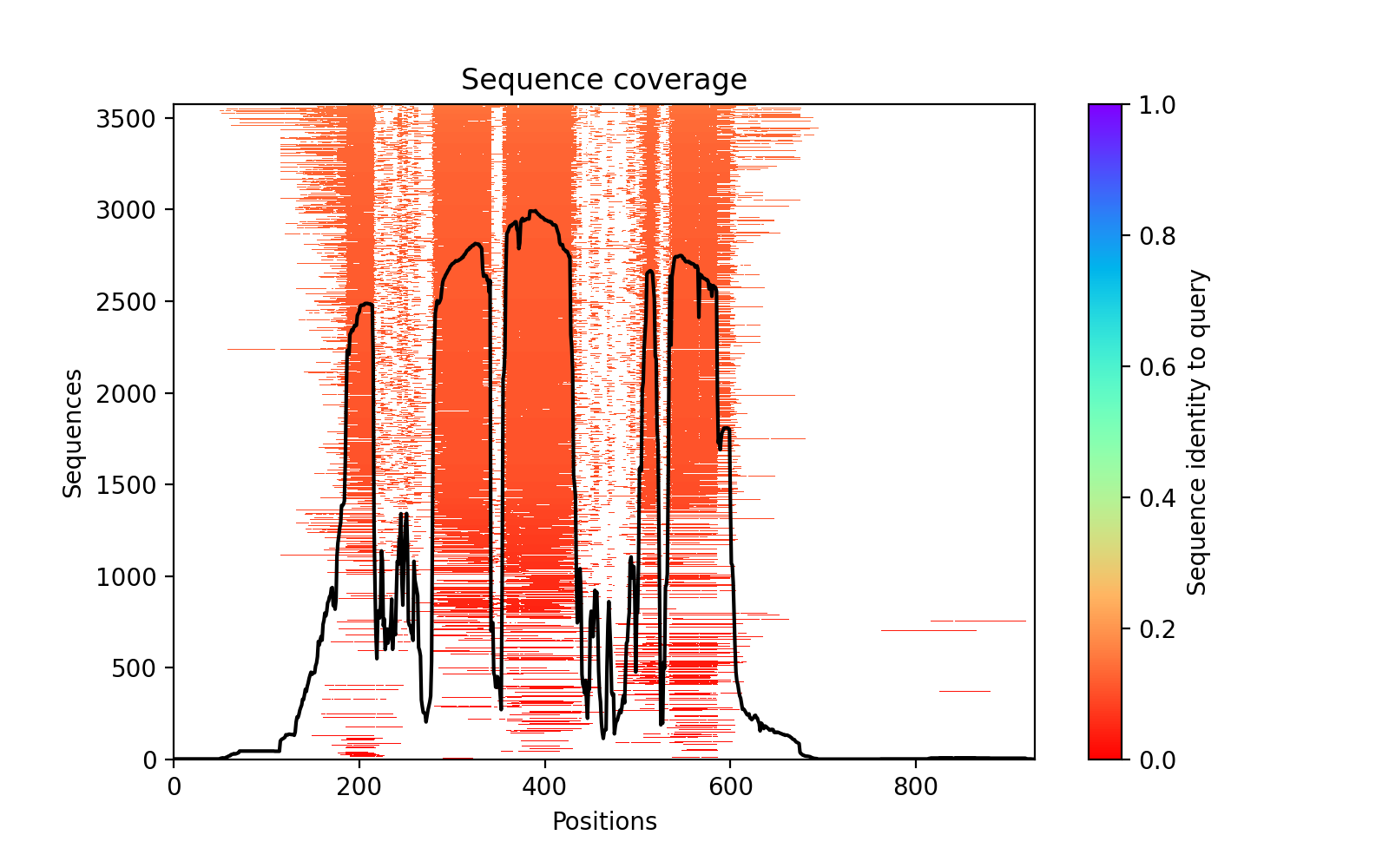

### AncSeptin_N4_385_1281_9397d_PAE.png

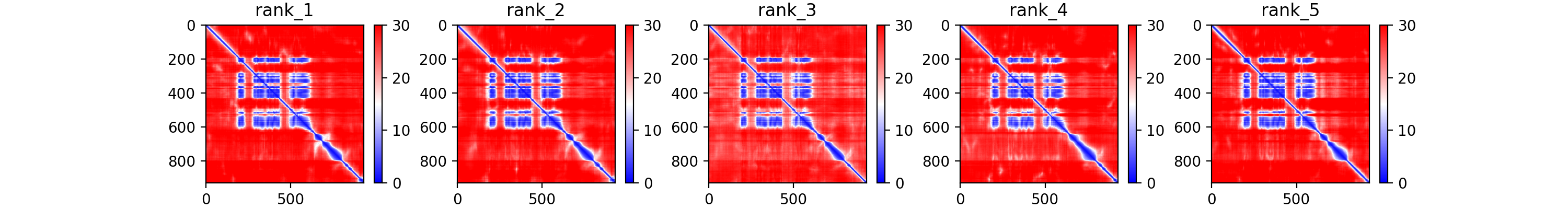

### AncSeptin_N4_385_1281_9397d_plddt.png

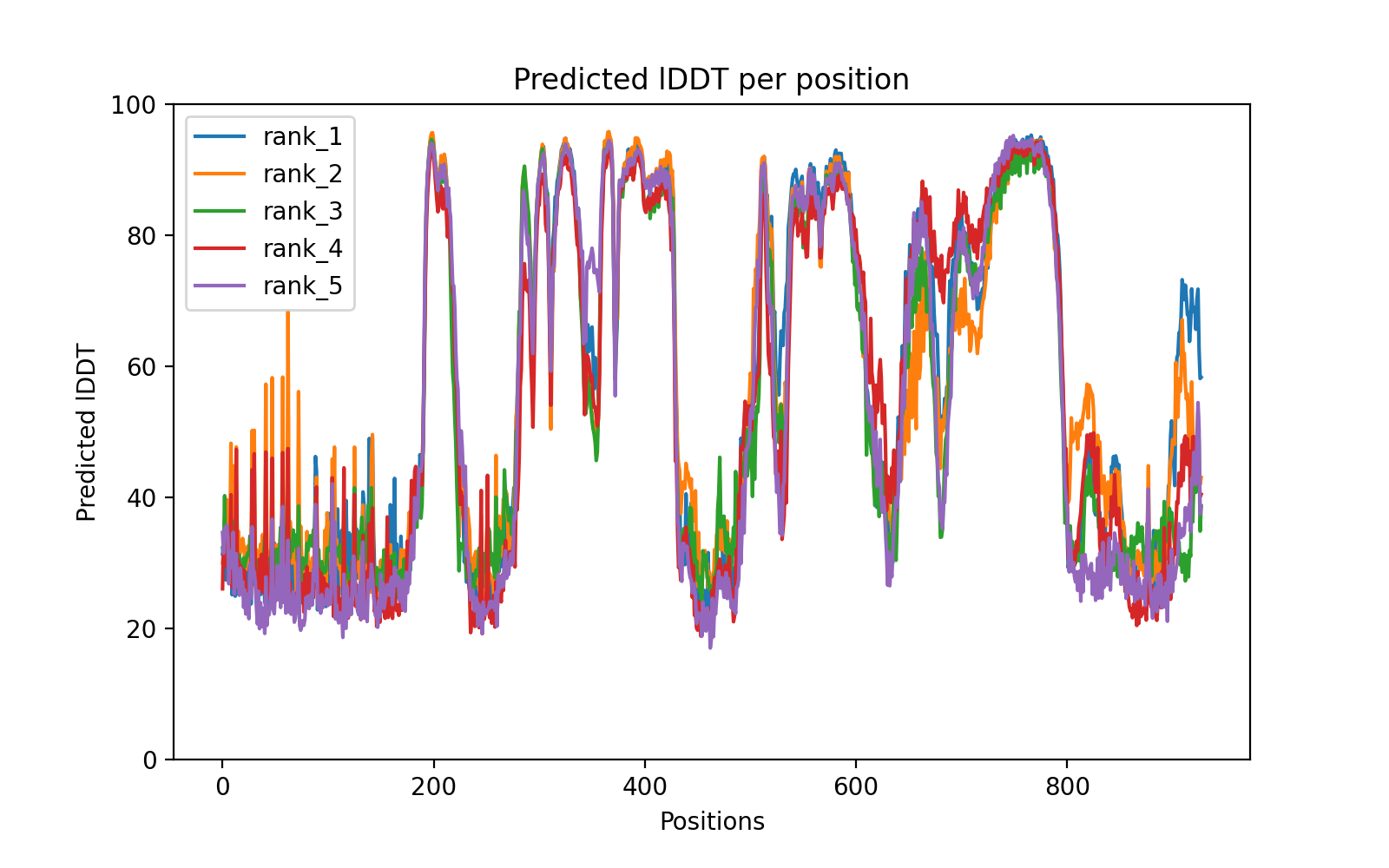

### OpisProtDiv_450_1264_931a9_coverage.png

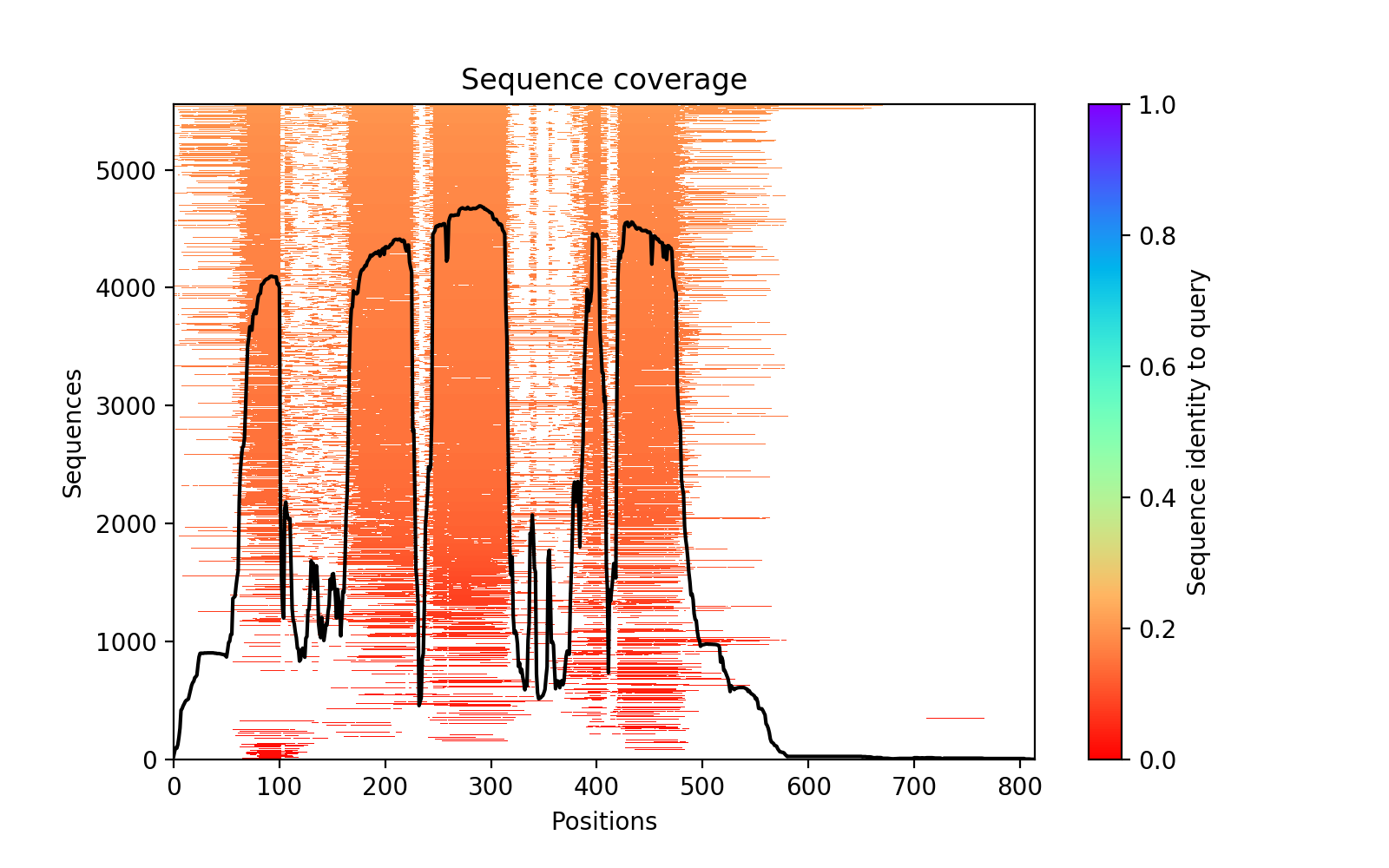

### OpisProtDiv_450_1264_931a9_PAE.png

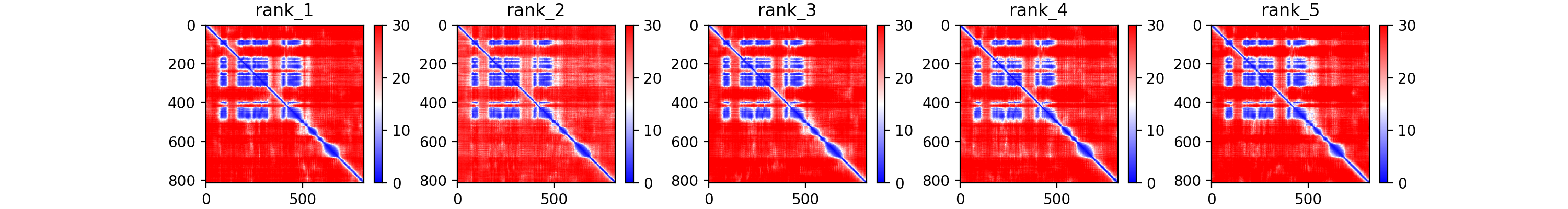

### OpisProtDiv_450_1264_931a9_plddt.png

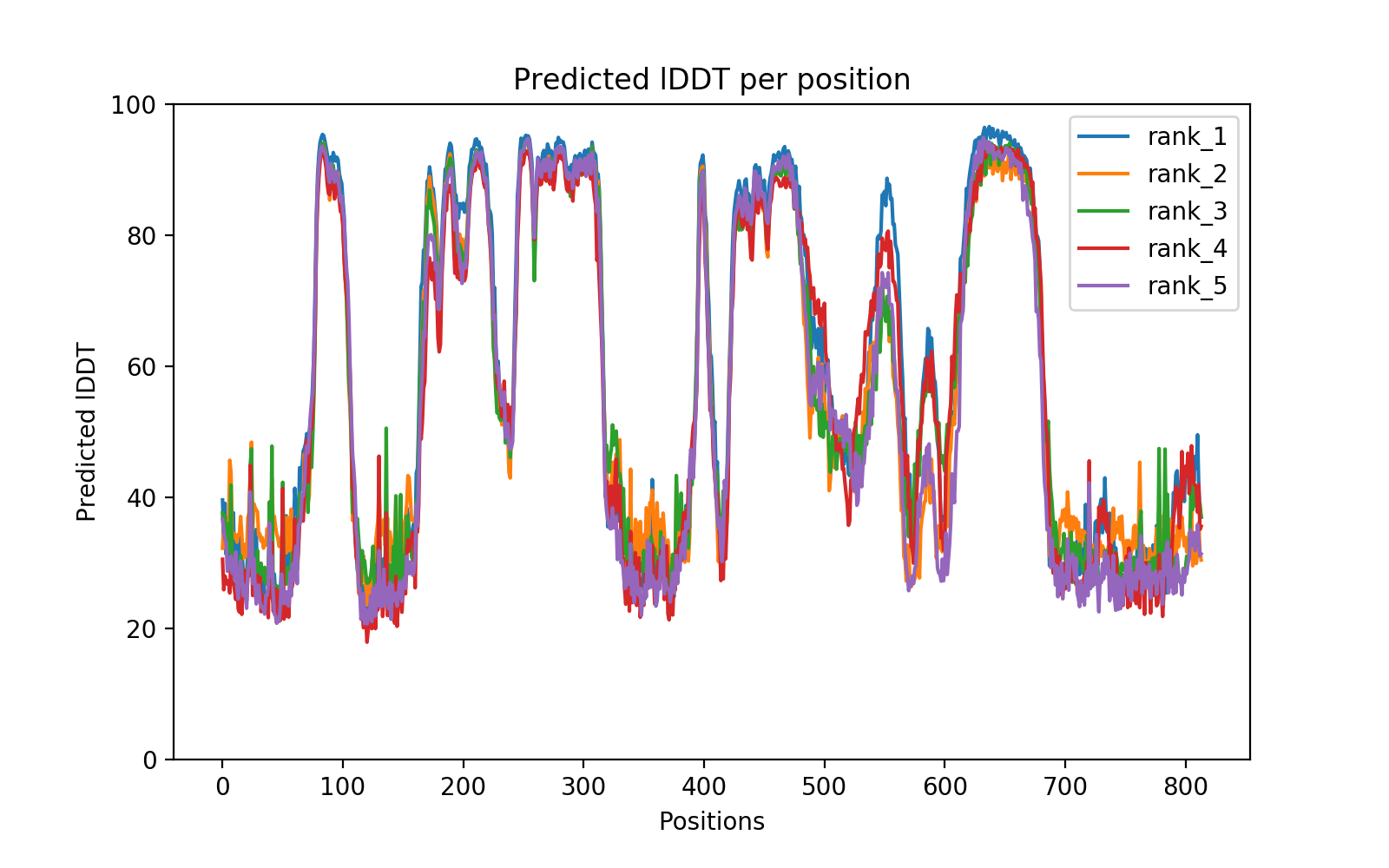
